# DepoCatalog: Mapping the Diversity of 105 Recombinant Klebsiella Phage Depolymerases Across Sequence, Structure, and Substrate Specificity

**DOI:** 10.1101/2025.09.09.675204

**Authors:** Aleksandra Otwinowska, Sebastian Olejniczak, Agnieszka Latka, Maria Pozniak, Grażyna Majkowska-Skrobek, Barbara Maciejewska, Janusz Koszucki, Vyshakh Panicker, Sara Jabłońska, Mathilde Hulsens, Joachim J. Bugert, Régis Tournebize, Stan J.J. Brouns, Flavia Squeglia, Rita Berisio, Yves Briers, Rafal Mostowy, Zuzanna Drulis-Kawa

## Abstract

A validated catalog of 105 recombinant depolymerases from Klebsiella phages covers 58 KL-types. 46 novel enzymes from prophages, jumbo phages, and common phages are linked to any known enzymatic activity against 14 classical serotypes and 12 genome-defined KL-types. Using activity-based profiling, structure prediction, and domain dissection, we developed a function-guided classification and a five-class structural catalog.

This framework reveals highly specific enzymes active against up to three capsule types. K47 CPS was degraded by three diverse protein groups. Structurally similar depolymerases degrading particular CPS were found in distinct phage taxa, with highly conserved enzymes in *Drulisvirus* specific to K1-, K2-types. The exclusive depolymerases were found in siphoviruses targeting K2 and K62 serotypes. A case study of five structurally similar enzymes degrading KL22/KL37/KL111 and KL25/KL119 capsules suggested specificity switching via amino acid changes or C-domain modification.

Klebsiella phage depolymerases catalog sheds light on their diversity, evolution, and potential application.

## Introduction

Capsular polysaccharides (CPS) are produced by encapsulated bacteria as a thick layer surrounding the cell and serving various functions. CPS plays a key role in bacterial virulence, mimicking host cell molecules, and protecting pathogens from host immune recognition and defense mechanisms, such as opsonization, phagocytosis, or complement-mediated killing (Sachdeva et al. 2017, Majkowska-Skrobek et al. 2016 & 2018, Zierkie et al., 2025, Liu et al. 2025). It also prevents bacterial desiccation, enhances the adherence to abiotic and biological surfaces, forms a biofilm matrix, and protects bacteria from recognition and infection by bacterial viruses (bacteriophages) (Majkowska-Skrobek et al. 2021, Wang et al. 2015).

*Klebsiella pneumoniae*, an opportunistic, highly virulent, and multidrug-resistant bacterium, is one of the most common encapsulated Gram-negative pathogens that uses the capsule as a major factor contributing to high invasiveness, immune clearance evasion, and cationic antibiotics sequestration (Sachdeva et al. 2017, Calfee 2017, Bengoechea & Sa Pessoa 2019). To date, more than 163 different *cps* loci encoding CPS biosynthetic enzymes have been identified, illustrating a huge diversity and complexity of genes responsible for the biosynthesis of *Klebsiella* capsule polysaccharides (Wyres et al. 2015, Wick et al. 2018, Lam et al. 2022).

It should be noted, however, that the diversity and genetic richness of genes encoding bacterial capsules have not yet been fully verified in terms of their sugar composition and the structure of CPS. There is currently one comprehensive database available for the structural characterization of polysaccharide serotypes K1–K82, which were experimentally verified and confirmed (K-PAM; Patro et. al. 2020). As a result, the knowledge of the CPS composition of capsule locus KL>100 capsule types is still missing; thus, we are lacking information about their structural similarities and whether they exhibit similar antigenic, virulence, or immunogenic properties.

As was mentioned above, the bacterial capsule forms a physical barrier, inhibiting phage adsorption to the cell receptors hidden beneath. Therefore, many phages developed specific virion-associated enzymes (depolymerases) that degrade the polysaccharide layers to initiate the infection (Pan et al. 2017, Majkowska-Skrobek et al. 2016 & 2018, Latka et al. 2021, Dunstan et al. 2021, Haudiquet et al. 2024). These enzymes are typically receptor-binding proteins (RBPs) enabling precise enzymatic degradation of bacterial polysaccharides (CPS, LPS) and biofilm matrix (Latka et al. 2019, Huang et al. 2024). Klebsiella phages differ in their RBP organization, ranging from single-type RBP to complex branching systems (2-14 RBPs) (Pan et al. 2017, Latka et al. 2019; Ouyang et al. 2022).

Phage-encoded depolymerases exhibit a conserved modular architecture and typically assemble into elongated homotrimers centered around a parallel β-helix core (Steinbacher et al. 1994 & 1996, Seckler 1998, Seul et al. 2014). Their structure comprises three principal domains: (i) an N-terminal structural region, (ii) a central enzymatic domain, and (iii) a C-terminal trimerization module contributing also to substrate recognition (Steinbacher et al. 1997, Freiberg et al. 2003, Barbirz et al. 2008, Squeglia et al. 2020). The catalytic center can be supplemented by a β-barrel insertion domain, which completes the active site architecture (e.g., KP34gp57, Kl-dep, Dpo43, K1-ORF34) (Maciejewska et al. 2023, Gorodnichev et al. 2023, Liu et al. 2020, Tu et al. 2022). The catalytic pocket of phage depolymerases can be located either within a monomer (intra-subunit) or at the interface between adjacent subunits (inter-subunit) (Leiman & Molineux, 2008).

Some enzymatic RBPs (e.g., KP32gp37) are equipped with an intramolecular chaperone at the C-terminus, guiding proper folding and possibly undergoing autoproteolytic cleavage upon trimer assembly, which locks the enzyme in its mature, protease-resistant conformation (Schwarzer et al. 2007, Majkowska-Skrobek et al. 2018, Latka et al. 2021).

Phage-encoded depolymerases are broadly classified into two main enzymatic categories based on their catalytic mechanisms: hydrolases (EC 3) and lyases (EC 4). Both classes facilitate the degradation of polysaccharides into soluble oligosaccharides, thereby disrupting the protective carbohydrate barrier surrounding bacterial cells (Davies and Henrissat 1995, Sutherland 1995, Pires et al. 2016, Latka et al. 2017).

Despite recent advances in the biochemical and structural characterization of phage-derived depolymerases, our understanding of the relationship between sequence, structure, and substrate specificity remains fragmentary. The current database of characterized depolymerases is limited, making it difficult to infer substrate specificity based solely on amino acid sequence similarity (Concha-Eloko et al. 2024, Boeckaerts et al. 2024). Although some enzymes with high sequence similarity display conserved specificity, others with little or no similarity can target the same capsular polysaccharide, challenging the assumption that sequence identity correlates with specificity (e.g., K2 type-specific enzymes). The existence of unrecognized evolutionary lineages of depolymerases highlights the need for experimental validation to accurately define depolymerase specificity. The full extent of capsule cross-reactivity and the determinants of substrate specificity remain poorly understood. Also, prophage-encoded depolymerases remain largely unexplored, since most available data focus on enzymes from virulent phages (Otwinowska et al. 2025). Taken together, these gaps emphasize the need for experimental validation of potential depolymerase candidates, as well as systematic studies integrating biochemical, structural, and functional analyses.

In this study, we present a catalog of 105 phage-borne depolymerases prepared as recombinant proteins that have been experimentally verified for their specificity towards *Klebsiella* capsule degradation. A highlight of this depolymerase catalog is, first, the specificity-guided classification of proteins targeting up to 58 KL-types, with very specific ones degrading only a single type of CPS and broad active enzymes recognizing and cleaving up to three different CPS (K-types). Secondly, integrating the structure prediction with the domain’s dissection analysis, we developed a five-class enzyme catalog, serving as a new conceptual framework for understanding the diversity and evolution of depolymerases. Combining the structure, specificity, and diversity of depolymerases in our catalog, we aim to answer several research questions:

1. What are the structural and specificity diversities of phage depolymerases targeting K. *pneumoniae* capsules?
2. What is the sequence and structure variability of depolymerases targeting the same capsule type? Can we predict their capsular tropism based on amino acid sequence or structure prediction?
3. How does the diversity of depolymerases reflect the genetic variability of *Klebsiella* CPS loci?
4. What is the specificity of the identified depolymerases toward *Klebsiella* capsular types, and how does it vary across different phage origins, including replication cycle and morphotype?
5. How do structural features and domain compositions correlate with the enzymatic activity and specificity of depolymerases?
6. What mechanisms underlie the ability of certain depolymerases to degrade multiple capsular types, and what structural determinants enable this versatility?
7. Can we use this knowledge to design novel depolymerases with a broad substrate range for therapeutic purposes or a narrow one for diagnostic applications?

Our findings expand the known repertoire of phage-borne depolymerases, which is accompanied by the huge diversity of *Klebsiella spp. cps* loci, thus CPS types, and their possible modifications. The unprecedented variability observed in these enzymes provides valuable insights into phage-host interactions, capsular specificity, and the evolutionary dynamics of phage-encoded enzymes. Moreover, this curated enzyme collection provides a valuable resource for further research and application in the field of phage biology, phage engineering, bacterial diagnostics, and antimicrobial strategies.

## Results

### Expanded dataset of Klebsiella phage depolymerases reveals broad serotype coverage and diverse origin

In this study, we have analyzed **105** distinct recombinantly produced depolymerases derived from Klebsiella phages, which were experimentally verified for their specificity towards Klebsiella capsule degradation, covering **58** different KL types. Among them, **59** proteins have already been documented in the literature (Cheetam et al. 2024, Zhao et al. 2024, Latka et al. 2021), and **46** were newly discovered by our team in Klebsiella prophages (20 proteins), Klebsiella jumbo phages (17 proteins), and common Klebsiella phages (9 proteins). Notably, **26** capsule types were targeted exclusively by depolymerases discovered here and had not previously been linked to any known enzymatic activity. This group includes 14 classical serotypes (K9, K14, K19, K22, K28, K32, K37, K38, K39, K46, K52, K60, K61, K68) and 12 KL-types (KL111, KL114, KL116, KL119, KL122, KL127, KL134, KL137, KL143, KL146, KL153, KL158). The KL-types are particularly noteworthy, as they represent genome-defined capsules that are increasingly reported in sequencing data but remain largely unexplored in the context of CPS composition (KL>100) and specific phage-derived enzymes.

Presented enzymes originate from five distinct sources: Klebsiella podoviruses (40 proteins), Klebsiella siphoviruses (9 proteins), Klebsiella myoviruses (11 proteins), Klebsiella jumbo myoviruses phages (25 proteins), and Klebsiella prophages (20 proteins). Each protein has also been analyzed by structure prediction using the AlphaFold3 tool and domain dissection to investigate the modularity of capsule-degrading enzymes, their specificity, and variability.

The enzymatic specificity of the recombinant depolymerases was assessed based on details provided in published papers, whereas proteins prepared within this study were evaluated by a spot assay on a broad *Klebsiella spp*. K-type panel ranging from K1-K82 to KL101-KL170 (with a few exceptions, including KL104, KL106, KL110, KL115, KL117, KL121, KL126, KL128, KL129, KL135, KL136, KL138, KL145, KL147-KL152, KL155-KL157, KL159-KL162, KL164 and KL165) (Supplementary Table S1).

To characterize and prepare a curated enzyme catalog, the key characteristics (targeted KL-type; depolymerase name and ID; annotation; nucleotide sequence; amino acid sequence; length in aa; domain features; catalytic site; cleaved bond; tested KL-types; phage name and replication cycle; phage morphotype; phage genome accession number; reference) for all collected depolymerases have been compiled in Supplementary Table S2.

### Capsule type versus enzyme diversity – taxa conserved enzymes, and up to 3 structurally diverse protein groups targeting one CPS type

The CPS-degrading enzymes were first grouped based on their substrate specificity (58 KL-types targeted) to assess the structural variability and sequence variability within particular capsule tropisms (Figure 1). In Figure 1, each group/subgroup includes only one protein representative targeting a particular capsule type, but the full list is shown in Supplementary Materials S1. All defined protein groups were also subjected to sequence (aa) and structural (TM score) alignments carried out on the N-terminal domain-deficient proteins to avoid interpretation bias caused by the conserved N-termini observed within phage genera (Supplementary Materials S1). That approach enables us to answer the initial research questions about the sequence, structure, and functional diversity of phage depolymerases targeting particular *K. pneumoniae* capsules, and about the variety of enzymes across different phage origins.

Seven K1-specific depolymerases derived from 6 podoviruses (58-98 aa %identity) and one jumbo phage (32-35 aa %identity) formed a structured uniform K1 group 1 (TM score 0.9) with the same organization of domains. It was interesting that 5 podoviruses from the *Drulisvirus* genus shared >95 aa %identity in depolymerase sequence, although they originated from distinct geographic regions and hosts.

The most representative serotype specificity was K2, with 13 specific depolymerases, including six that have been proven also to degrade K13 CPS. It is important to mention that K2 and K13 CPS share the core polysaccharide unit, and K13 is extra decorated with galactose. Depolymerases from group 1 cleave α-D-Glc-(1→3)-β-D-Glc in K2/K13 CPS, whereas those from group 2 degrade β-D-Glc-(1→4)-β-D-Man in K2/K13 CPS (Figure 1, Supplementary Materials S1, Table S2; Dunstan et al. 2021, Lin et al. 2022, Ye et al. 2024). Each group was structurally homogeneous with a high TM score (0.9-1.0), except DpK2 enzyme; thus, we might assume that all these enzymes are also able to digest K13 CPS, even if not tested experimentally. Interestingly, the K2 group 1 was almost identical in terms of amino acid sequence (>97 %identity), and all five were *Webervirus* (siphovirus) proteins originating from geographically distinct regions, which might suggest this group is exclusive to the sipho-morphotype. The differences in TM score between DpK2 and the rest of the K2 group 1 representatives were verified in PyMOL, showing that subtle changes in amino acid content resulted in modified spatial orientation of the C-terminus predicted by AlphaFold 3 (Supplementary Materials S1). Nevertheless, while AlphaFold 3 can occasionally capture conformational changes arising from single or a few amino acid substitutions, its predictions are not intrinsically sensitive or reliably accurate at that scale, especially for peripheral or flexible regions such as C-terminal regions. The observed difference in C-terminal orientation between DpK2 and the other K2 group 1 representatives could therefore reflect model uncertainty rather than a genuine structural shift.

The second K2 group was more diverse, encompassing 5 podovirus proteins, along with one protein derived from sipho-, myo-, and jumbo myovirus, showing the aa % identity at a low level (28%-55%). Similarly to K1 targeting depolymerases, the K2-specific enzymes from four *Drulisvirus*es were almost identical in sequence (96-98 aa %identity), despite the divergent origin of isolation and hosts.

Since K2 group members differ in terms of their C-terminal domain, we checked the structural similarity of the central domain only, and the TM score varied between groups from 0.5 to 0.7, indicating the same fold, which—when supported by sequence similarity—often implies a common evolutionary origin.

The K21 group (TM score >0.9) was formed by two podoenzymes sharing 86% identity, and one jumbo phage enzyme with low aa similarity (39%). The K23-specific group, characterized by nearly structurally identical proteins (TM score 0.9), consisted of three prophage-derived proteins with high amino acid sequence similarity (98%), as well as two enzymes originating from podo- and myoviruses. In several other cases (K23, K28, and KL111), both lytic and prophage-associated enzymes were found to target the same CPS, having (Tm score >0.9; 29-99 aa% identity).

In order to explore the sequence and structural variability of depolymerases targeting the same capsule type, we extended our analysis to include enzymes beyond those specific for K1 and K2. There were other K-types in which we found 2 structurally diverse protein groups: K3, K20, K46, KN4=KL104, with a 1:1 representation, K62 (2:2), and K64 (3:2). Enzymes specific to K62 formed two structure- and sequence-wise coherent groups (1 and 2), originating either from lytic Webervirus (siphovirus) or from prophages also belonging to siphoviruses. These might represent another example of probably sipho-morphotype exclusive depolymerases. Five enzymes degrading serotype K64 formed two groups (1 and 2) with different domain organizations, a TM-score of 0.9 within each group, and high amino acid similarity (98%), but only within proteins derived from podoviruses (group 1). Interestingly, for K47, even 3 structurally diverse groups were observed. Depolymerases originating from podoviruses form groups 1 and 2, with enzymes sharing domain composition, the TM score (0.6), but low aa %identity (31-32%). The Dpo43 podoenzyme stood out from the others with a different domain organization, low structural and sequence similarity, belonging to the K47 group 3.

The other point of analysis was the substrate specificity range of the individual enzyme. It turned out that the vast majority of proteins (91 out of 105) showed narrow activity only to a single capsule type. Six enzymes (K2/K13, K3/KL146, K21/KL163, K25/KL119, K30/K69, K57/K68) exhibit double substrate specificity, likely by targeting the same glycosidic linkage in two structurally distinct polysaccharides (probably valid also for not yet characterized CPS compositions of KL > 100). There was one exceptional protein, KLEO13 gp10, recognizing and cleaving three different CPS types K22/K37/KL111.

### Structure-based classification of Klebsiella phage capsule depolymerases - catalog

A characteristic feature of depolymerases is the β-helical central domain, where the homology to lyase or hydrolase is predicted, and the catalytic pocket is located. This domain is preceded by an N-terminus, which can vary significantly in shape and length (ranging from a couple to hundreds of amino acids), and was assigned a structural function necessary for the attachment to the virion or attachment to the preceding tailspike/tail fiber in the RBP branching system (Stummeyer et al. 2006, Leiman et al. 2007, Prokhorov et al. 2017, Plattner et al. 2019, Latka et al. 2019). On the other end of the β-helix, at the C-terminus, domains were reported to possess the function in host receptor recognition or protein trimerization (Steinbacher et al. 1997, Freiberg et al. 2003, Barbirz et al. 2008, Olszak et al. 2017). To investigate whether there are any purpose-built motifs or domain architectures that correlate with the substrate specificity, we attempted to categorize depolymerases based on the overall protein structure and domain dissection, omitting the N-terminus due to high conservation amongst the closely related phages and morphotypes, and lack of influence on enzymatic activity (Latka et al. 2021).

Depolymerases were classified into five principal structural classes based on their overall architectural features (Figure 2). In Figure 2, each class includes only one protein representative targeting a particular capsule type.

**Class 1, ‘classical depolymerase’**, is the most abundant, with 29 members. In this class, the β-helical central domain, located between cap 1 (the last α-helix preceding β-helix) and cap 2 (first α-helix or loop after β-helix), is composed of 10 to 20 rungs, which are followed by visually separated C-terminal domain(s). This class is subdivided into six subgroups (A-F) that can be distinguished depending on the C-terminal domain structure. Class 1A (6 members, e.g., K47_P560dep) is characterized by the C-terminal domain having its β-sheets arranged perpendicularly to the β-helix, resembling a **‘vertical stand’**. This class includes enzymes specific to various capsular types, each characterized by distinct polysaccharide subunit composition (K9, K23, K47, K57/K68, K62, KN5-KL105). Class 1B proteins have the β-sheets of the C-terminus parallel to those in the β-helix, resembling a sort of **‘pouf’**. Eleven members represent the capsule tropism to K2, K11, K13, K20, K22, K25/KL119, K37, K60, K61, KN3=KL103, KL111, and KL153. Among 6 depolymerases specific to K3, K19, K28, K56, K64, KL137, and KL146, a **‘spout’** shape could be noticed as an addition to parallel β-sheets of the C-terminal domain, forming Class 1C. In one case of Class 1D (K47_Dep42), the monomers of the C-terminal domain did not trimerize as a common structure but as three separate units forming a kind of **‘tripod’** (pLDDT >90) (Supplementary Materials S1). Five depolymerases targeting K51, KN4=KL104, KL134, KL143, and KL158, were grouped in Class 1E, having exceptionally long central domain (∼20 rungs), followed by only a couple of β-strands-long, tapering **‘short C-domain’**. Finally, Class 1F with **‘double C-domain’** was distinguished based on 6 representatives, being also the most diverse subgroup of classical depolymerases. K2/K13_B1dep has a C-domain composed of β-sheets parallel to the β-helix, followed by a needle, which is formed of α-helices. KP32gp38 and K11gp0043 possess two domains, the first

with β-sheets parallel, the second perpendicular to β-helix, named in the published report as CBM-LEC duet (Squeglia et al. 2020, Latka et al. 2025). Remaining representatives of Class 1F, K46_248_38, K54_gp531, and KL127_319_37, possess two visually separate modules at the C-terminus with diverse shapes and folds. Some of the C-terminal domains of Class 1A and 1F were found homologous to carbohydrate-binding module (CBM) and/or lectin, however, with a low score in DALI.

**Class 2** was distinguished based on the presence of an additional structure element, formed by antiparallel β-sheets reaching out of the central β-helix, called **‘insertion domain’**. Insertion domain varied in terms of location: Class 2A - close to cap 1, **‘classical insertion domain’**, Class 2B - in the middle of the enzymatic domain, **‘middle insertion domain’**, Class 2C - close to cap 2, **‘low-lay insertion domain’**. There was one protein (K47_Dpo43) possessing two insertion domains present close to the cap 1 and ending with a small C-terminus. This protein formed Class 2D - **‘double insertion domain’**. Class 2B was also represented by only one protein, K20_Kl-dep, which possessed a ‘pouf-like’ C-terminal domain. Class 2C (K62_0391_11) was featured with a ‘vertical stand-like’ C-terminus, while the structure of the C-terminal domain of five representatives belonging to Class 2A was more diverse, degrading K1, K5, K14, K63, and K64 CPS.

**Class 3, ‘tail fiber domain in depolymerase’** has a short or middle-length central β-helix core and possesses a long β-strand-rich region, visually resembling a tail fiber (as also indicated by DALI search). These enzymes are divided into two subgroups. Class 3A **‘tail fiber domain + classical C-domain’** included enzymes specific to K8, K30, K32, K39, K46, K69, and KL116, whereas Class 3B **‘tail fiber domain + chaperone’** contained enzymes with an α-helix-rich domain with most predicted peptidase domain (pfam13884), and targeting K3, K7, K35, K52, and KL122.

**Class 4** – **‘αH-containing central domain’** depolymerases feature classical β-helix supplemented with α-helices and loops inserted between the β-sheets of the different rungs, a morphology observed for five depolymerases. This class was further divided into two subgroups depending on both further differences in the central domain and the C-terminus. In Class 4A, the central domain is short (9-10 rungs), followed by a double C-domain with the first resembling a short fragment of tail fiber, and the second ‘vertical stand-like’. Interestingly, this kind of special organization was exceptionally presented by depolymerases specific to KN1=KL101 and KN4=KL104, which could suggest a unique polysaccharide subunit composition in these capsule types. In Class 4B proteins degrading K27, K38, and KN2=KL102 CPS, the central domain reassembles a duplication of the central domain from class 4A proteins in the form of a mirror image, which results in a very irregular shape. The C-terminus is also composed of two parts: first longer tail fiber-like fragment, followed by an α-helix-rich chaperone with most of a peptidase domain predicted (pfam13884).

**Class 5** was formed for two representatives (KL108_Dep108.1 and KL114_KP24gp307) having **‘colanidase-like depolymerase’** domain with a long central domain and C-terminus, which is spread out.

The last step was the total comparison of the structure similarities between all central and C-terminal domains (Supplementary Materials S1). Most of the central domains from distinct protein classes shared the structure conformation at the level of > 0.5 TM score, with some exceptions in Class 1E, Class 2 “insertion domain”, and Class 4B (TM score < 0.4). In contrast, the variety of C-terminus domain structure turned out to be very high (TM score < 0.4), with only a few domains similar to each other (TM score 0.5). It suggests that the enzymatic part of the enzyme is much more conserved compared to the C-terminus.

### Capsular Specificity Switching Induced by Amino Acid Changes or C-domain Modification

In this section, we aimed to explore the relationship between architectural similarities and the different substrate specificity of depolymerases. Despite being highly similar in structure (TM score >0.9), five enzymes from Class 1B “pouf” (KLEO13g10, 184_43, KANgp232, KP24gp300, and S2-2) exhibited markedly different specificity profiles in experimental assays.

While 184_43 and KANgp232 depolymerases were active against only a single KL111 capsule type, the KLEO13gp10 was capable of degrading up to three distinct polysaccharides (K22, K37, and KL111). The sugar subunit similarity between K22 and K37 types (differing only in galactose acetylation residue) might suggest the same degradation site; therefore, we assumed that the unknown KL111 polysaccharide should be composed accordingly. We have also included two other proteins in the detailed analysis (KP24gp300 and S2-2), which are specific to the K25 serotype, differing from K22/K37 by the inverted position of two substituting sugars within the CPS repeating unit. Moreover, KP24gp300 proved to be additionally effective in the degradation of KL119 polysaccharide, which has an unknown structure.

This observation prompted us to investigate whether amino acid composition or subtle structural differences, particularly within substrate-binding regions/catalytic pocket or domain interfaces, could clarify such substrate specificity differences. Our analysis went through several steps (Figure 3, Supplementary Figure S1-S5): (1) comparison of available data regarding sugar composition building K22, K37 and K25 CPS; (2) comparison of *cps* clusters involved in the biosynthesis of analysed capsule polysaccharides (K22, K37, KL111, K25, and KL119); (3) analyses of sequence and structure similarities between five assessed proteins, including amino acid conservation, and mapping the active site within the three-dimensional structure of the enzyme by evaluating the surface electrostatic potential.

**Figure 3.**
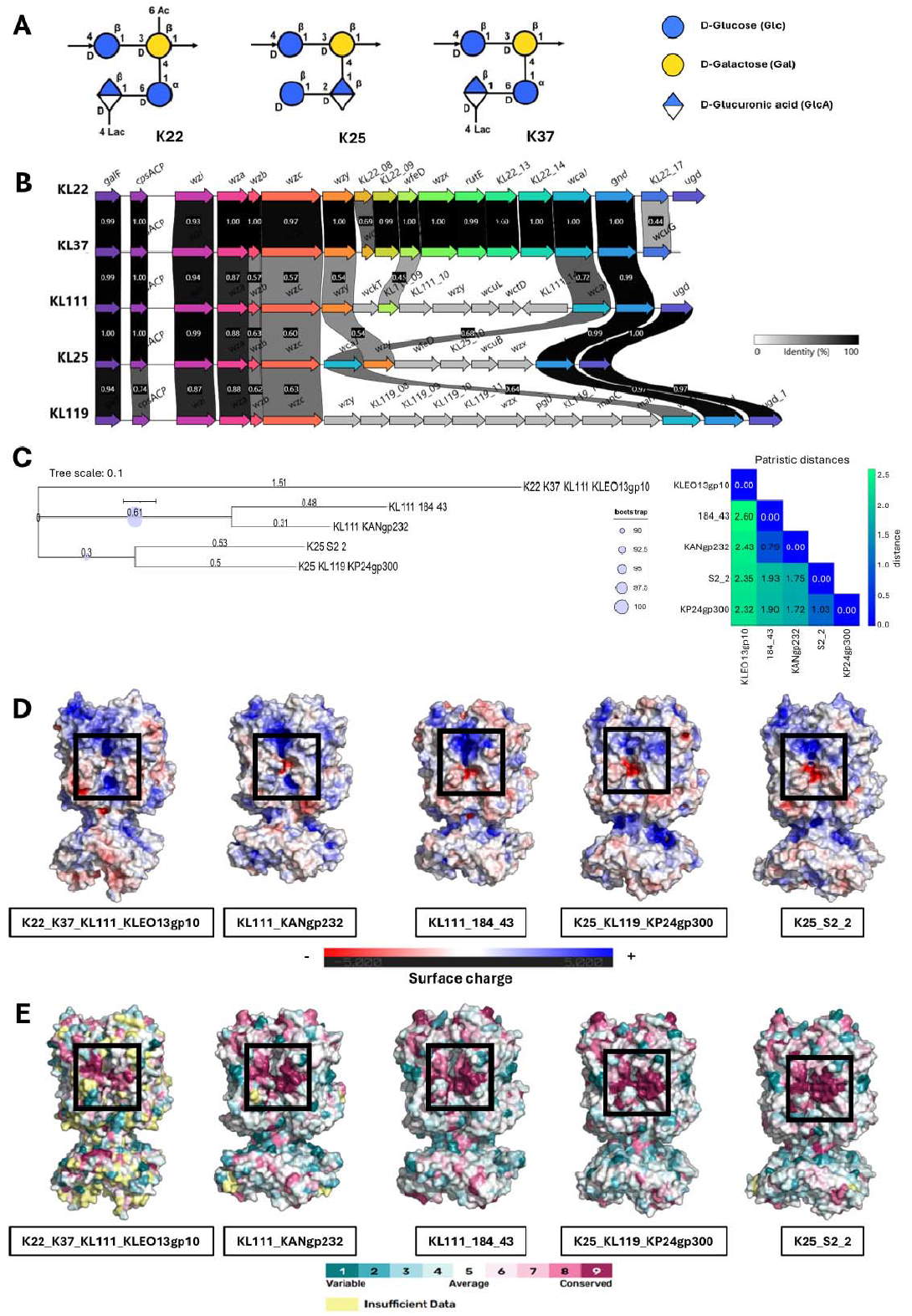
Comparative analysis of five selected phage depolymerases and targeted CPS. (A) Structural representation of three *K. pneumoniae* CPS subunits of K22, K25, and K37 serotypes. Each subunit is shown in a cartoon-style model. Images were sourced from the K-PAM database (https://iith.ac.in/K-PAM/home.html), *a Klebsiella* serotype predictor and surface antigen modeler; (B) Comparative genomic organization of KL22, KL25, KL37, KL111, and KL119 CPS loci. Gene clusters are displayed with annotated similarities, enabling visualization of conserved and variable regions across the loci (Clinker); (C) Phylogenetic tree constructed in IQ-TREE using JTT+G substitution model from multiple sequence alignment of KLEO13 gp10, KANgp232, 184_43, KP24gp300, and S2-2 depolymerases. Bootstrap support visualized as blue dots, where size reflects bootstrap value magnitude (details in MM section). (D) Surface electrostatic potential maps of analysed depolymerases generated in PyMOL with the APBS plugin. Regions highlighted with black squares are putative locations of the active site, based on the most negative and positive charges accumulation and their location in the groove between monomers (details in MM section). (E) Amino acid conservation of the analysed depolymerases was made in ConSurf. Amino acid residues are colored according to conservation scores. Regions highlighted with black squares are possible locations of the active site, based on the grouping of highly conserved amino acid residues.

### Polysaccharide similarity does not reflect cps gene cluster similarity, except for K22/K37

Serotypes K22 and K37 have identical monosaccharide composition in the subunit, except for the presence of an acetyl group at C6 of galactose in serotype K37. When comparing these serotypes with K25, the same sugars (β-D-glucose and β-D-galactose) are observed in the main chain, but there are differences in the side chain. In K22 and K37, the side chain starts with α-D-glucose attached to C4 of the main chain β-D-galactose, then β-D-glucuronic acid is attached to C6 of α-D-glucose. There is also a lactyl group attached to C4 of β-D-glucuronic acid. In K25, the side chain is organized in a reversed pattern. Additionally, there is no derivative of lactyl acid attached to β-D-glucuronic acid in K25 (Figure 3A).

Comparisons of the KL22, KL25, KL37, KL111, and KL119 CPS synthesis loci (Figure 3B) showed the presence of two parts: core conserved genes (*galF, cpsACP, wzi, wza, wzb, wzc*) and a more diverse, variable region, responsible for the biosynthesis of specific sugars. Loci KL22 and KL37 are almost identical, with differences limited to two genes - *wcmA* and *wcuG*, with similarity of 69% and 44%, respectively. There is also a presence of the *ugd* gene in the KL22 locus, which is absent in the KL37 locus. The remaining genes have a high level of identity (93% - 100%), which correlates with the similarity of their corresponding CPS structures. In the KL111 locus, two genes – *wzy* and KL111_09-show similarity (54% and 45%) to *wzy* and *wfeD* from KL22 and KL37 loci, respectively. The KL111 locus *wzy* gene also shares 54% similarity with the *wzy* gene from the KL25 locus. The *ugd* genes present in all, except the KL37 locus, have similarity in the range of 97% - 100%. Among all analyzed loci, two genes are present in all of them – *wcaJ* and *gnd*, with variable similarity ranging from 64% to 100%. Based on the 5 KL-loci comparison, it would not have been possible to predict the similarity in CPS composition among 22/37, 25, 111, and 119 K types. Moreover, there is no obvious relation between enzyme specificity and the KL-locus structure.

### Structural and Sequence Limitations in Predicting Protein Substrate Specificity

Comparisons of amino acid sequences among these depolymerases showed that the highest level of sequence identity (55% and 42%) occurred between depolymerases degrading the same serotype – KL111 (KANgp232 and 184_43) and K25 (KP24gp300 and S2_2), respectively. In the remaining cases, the sequence identity ranged from 29% to 36%. The most dissimilar in terms of sequence identity was depolymerase KLEO13gp10 active against three serotypes (K22, K37, KL111), whose aa %identity to the other enzymes was only 28-32%. Detailed data are summarized in Figure S2 (Supplementary Figure S1).

Phylogenetic tree and patristic distance analyses (Figure 3C) revealed clustering patterns. Depolymerases 184_43 and KANgp232 (both KL111) form the closest pair (distance ∼0,79), indicating strong relatedness. Similarly, S2_2 (K25) and KP24gp300 (K25 and KL119) are closely associated (distance ∼1.03). In contrast, KLEO13gp10 (K22, K37, and KL111) is the most distantly related to all others (2.32 to 2.60). KLEO13gp10 did not group with either cluster, instead showing a weak association to the K25 proteins (closest to KL24gp300 at ∼2.32, and S2_2 at ∼2.35). Its greatest divergence was from 184_43 (∼2.60) and KANgp232 (∼2.43), indicating stronger separation from the KL111 proteins. KLEO13gp10 may represent a distinct lineage, more loosely connected to the K25 group than to KL111. Bootstrap replicates confirmed its unstable placement, as KLEO13gp10 did not form a supported clade with any group, likely due to limited phylogenetic signal or its high divergence.

The three-dimensional structures of all five depolymerases (excluding the N-terminal domain) were compared (Supplementary Figure S2), and the TM-score ranged from 0.89 to 0.98, indicating high structural similarity. Analysis of structural alignments (Supplementary Figure S2) confirmed the agreement in the spatial arrangement of their structures. Further comparisons were performed separately for central domains and TM-score ranged from 0.93 to 0.98, while for the C-terminal domains, the values were lower – 0.74 for KLEOgp10 against others, 0.83-0.86 between K22 and K25 specific proteins, and up to 0.93-0.95 within the same specificity, indicating greater structural similarity in the central domain and more variability in the C-terminus (Supplementary Figure S3).

Analysis of surface electrostatic potentials (Figure 3D) revealed a significant accumulation of negative and positive charges occurring in the space between monomers in all analyzed depolymerases, suggesting that this region may harbor the active site (intersubunit position). However, in KLEO13gp10, there were fewer negatively charged amino acids in this region, but they were more located on the solvent-exposed surface of a single monomer - also present in S2_2 depolymerase. Other groups of charged residues were observed at the narrowing between the central and C-terminal domain, most prominently in KP24gp300, but also evident in the remaining depolymerases. Additionally, a distinct cluster of negatively charged residues was identified on the underside of the KLEO13gp10 C-terminal domain, which appeared weaker in other analyzed enzymes.

Analysis of amino acid residue conservation (Figure 3E, Supplementary Figure S4-S5) revealed the presence of highly conserved amino acid residues within the intermonomer region corresponding to the above-described putative active site location. Several conserved motifs (Supplementary Figure S4-S5) were also identified, indicating a possible universal catalytic mechanism for most of the analyzed enzymes. Many of these motifs were located in the intermonomer region. In the C-terminal domain, conserved motifs were mainly observed in the depolymerases KANgp232, 184_43, KP24gp300, and S2_2, whereas KLEO13gp10 displayed a different conservation pattern and didn’t share these motifs. It should be noted that the conservation profile of KLEO13gp10 was derived from a smaller number of homologous sequences (n=9), which may limit the resolution of residue-specific conservation. Nonetheless, the clustering of conserved residues in the intermonomer region across analyzed enzymes supports its functional importance.

## Discussion

Phage depolymerases exhibit remarkable structural and functional diversity. The extensive diversity of *Klebsiella* capsules, shaped by variations in sugar composition, stoichiometry, and *cps* locus organization, is paralleled by the high variability of RBPs equipped with capsule-specific depolymerase domains (Latka et al. 2019). These enzymes are typically highly specific, recognizing and degrading the capsular polysaccharides of a single serotype. Exceptions are rare and usually limited to serotypes with closely related polysaccharide structures, such as K2/K13 and K30/K69, which share an identical sugar backbone, often resulting in cross-reactive enzymatic activity (Pertics et al. 2021, Hsieh et al. 2017). To date, 59 recombinant *Klebsiella* phage depolymerases have been reported before, targeting 32 distinct capsular types, with the most significant number of depolymerases reported for serotypes K1 and K2 (Supplementary Table S2).

Predicting substrate specificity from sequence data alone remains difficult due to the limited number of experimentally validated depolymerases. While some enzymes with high sequence similarity share substrate preferences, others with low sequence identity can still target the same capsule type (Latka et al. 2021). Structural modeling provides more reliable insights, especially when focusing on the catalytic and substrate-binding domains. However, even structurally similar enzymes can differ in activity due to subtle changes in key residues in the sugar-interacting pockets. Machine learning tools and bioinformatics pipelines are being developed to improve predictive accuracy (Boeckaerts et al. 2024, Concha-Eloko et al. 2024, Gaborieau et al. 2024). Still, these models require larger datasets of characterized enzymes to be truly effective. Hybrid approaches combining sequence, structure, and biochemical data offer the most promise. Ultimately, while predictions are improving, they are not yet sufficient to replace empirical testing.

It must be stressed that all 105 enzymes presented here were expressed as recombinant proteins and experimentally confirmed for their specificity in degrading *Klebsiella* capsular polysaccharides (published studies and this work), most using a large panel of K-types. Notably, 46 of these enzymes targeting 26 unique K-types are novel discoveries generated within this project and originating from three distinct phage sources: Klebsiella prophages, Klebsiella jumbo phages, and Klebsiella common lytic phages. By analyzing the unique structural, functional, and phylogenetic diversity of capsule depolymerases encoded by Klebsiella phages, we established a comprehensive **catalog** encompassing depolymerases targeting **58** distinct **capsule types**. Based on domain architecture, we proposed a systematic classification scheme comprising **five principal classes and several sub-classes**. This framework provides clear insights into the remarkable evolutionary diversification of **Klebsiella** phage depolymerases, which appears to mirror the extensive heterogeneity of capsule types within the host species.

Crucially, this classification may serve as a blueprint for standardization and future systematic investigations. It enables the categorization of newly discovered depolymerases and facilitates the definition of additional classes or subgroups, thereby progressively enhancing the resolution with which we can map the natural diversity of these enzymes. Notably, while our dataset covers depolymerases targeting approximately one-third of the currently described capsule locus types, it underscores the likelihood that substantial diversity remains uncovered.

### What are the sequence, structural and functional variations of phage depolymerases targeting Klebsiella pneumoniae capsules, and how do these differences correlate with capsule type specificity, phage taxonomy, lifestyle (virulent vs. temperate), and virion morphology?

Scientific understanding of the origin of enzymatic RBPs can be traced back to research conducted in the 1980s with the statement that tailspike enzymes degrading cell surface polysaccharides can be produced by tailed phages belonging to the (former)*Myoviridae, Siphoviridae*, and *Podoviridae* families (Rieger-Hug and Stirm 1981, Leiman & Molineux 2008).

Nevertheless, most identified depolymerases are encoded by phages exhibiting a podovirus morphotype. *K. pneumoniae* podoviruses typically show simple RBP structures, with most encoding only one depolymerase-bearing RBP. However, some *Przondovirus* phages have developed dual-RBP systems enabling recognition of two, and in the case of KP32, three capsular types (RBP1 for K3 and RBP2 for K21/KL163) (Latka et al. 2019). Examples include Klebsiella phages K11 (recognizing K11 and K31), K5-4 (K5 and K8), KN3-1 (K56 and KL103), and KP32 (K3 and K21/KL163), all of which encode two RBPs with distinct enzymatic specificities (Latka et al.2021, Hsieh et al. 2017, Pan et al. 2019, Majkowska-Skrobek et al. 2018, Liu et al. 2020). A particularly interesting case is Klebsiella phage vB_KpnP_IME205, which encodes two depolymerases (Dpo42 and Dpo43) that both target the K47 capsular serotype but act on a different subset of K47 strains, indicating subtle intra-serotype variations in this capsule composition (Liu et al. 2020).

Sipho-type phages infecting *Klebsiella* display a simple RBP architecture, usually including a single RBP (Latka et al. 2019). Only eight depolymerases have been described in the literature. Interestingly, five of them are active against the K2 capsular type; among these, four proteins (Depo32, DpK2, BMacgp22, and B1dep) share over 98% amino acid identity, strongly indicating they are variants of the same depolymerase (Cai et al. 2019, Dustan et al. 2021 & 2023, Pertics et al. 2021). The remaining three siphoviral depolymerases (Dep1011, K62-Dpo30, and KP36gp50) target serotypes K5, K62, and K63, respectively (Li et al. 2024, Pan et al. 2024, Majkowska-Skrobek et al. 2016).

Myo-type jumbo *Klebsiella* phages exhibit the most complex branching RBP architecture, often comprising multiple depolymerases and therefore have accordingly the broadest host ranges among all known phages targeting this species (Latka et al. 2019). The most elaborate RBP system was described in the jumbo phage vB_KpM_FBKp24, with 14 distinct RBPs, 11 of which exhibit confirmed depolymerase activity (this study). Together, these enzymes enable recognition of 13 different capsular types — including two with dual activity (K2/13, K19, K25/KL119, K35, K46, K61, K64, KL114, KL134, KL137, KL158) highlighting the exceptional host range of this phage (Ouyang et al. 2022; this study).

Analysis of the depolymerase catalog revealed that several CPS types (specifically K2, K20, K46, K47, K62, K64, and KL104) were targeted by two structurally distinct groups of enzymes. In most cases, these enzymes were encoded by phages belonging to different taxonomic lineages. This observation might suggest that Klebsiella phages have independently evolved structurally divergent enzymatic solutions to degrade the same capsular serotype, a clear example of convergent evolution. The recurrence of this phenomenon across multiple CPS types underscores the strong selective pressure exerted by capsule structures and highlights the evolutionary plasticity of phage-encoded depolymerases in adapting to similar functional challenges imposed by the capsule structure.

Morphological differences also influence evolutionary dynamics; for example, myoviruses may evolve tail-associated depolymerases differently than siphoviruses or podoviruses. It is proposed in the literature that recombination events are common in phage evolution and contribute to the mosaic nature of RBP genes (Casjens &Molineux 2012; Hendrix et al. 2002). Environmental pressures, such as host capsule diversity, drive the need for depolymerase innovation. Overall, evolutionary processes vary widely depending on phage type, genome size, and ecological context (Krupovic & Bamford 2011).

Indeed, within several capsule-specific groups (including K1, K2 group2, K5, K11, K21, K23, K28, K30, K57, K63, K64, and KL111), we identified depolymerases encoded by both lytic and prophage with a podovirus, myovirus/jumbo morphotype, that share a conserved domain organization and almost identical structure (TM score 0.9) despite notable differences in amino acid sequence identity (30-40 aa%). This pattern suggests that these enzymes may have evolved independently across distinct taxonomic lineages yet retained a conserved spatial configuration and substrate specificity. Such structural convergence, coupled with sequence divergence, points to strong functional constraints that preserve enzymatic architecture critical for capsule degradation, potentially driven by parallel evolutionary pressures across diverse phage backgrounds.

Significant aa% identity differences for a given specificity were seen in the depolymerase sequence, but only across phage genera. In contrast, *Drulisvirus* depolymerases from K1 group 1 and K2 group 2 turned out to be very conserved in terms of the structure and aa sequence, even if derived from geographically distant sources and propagated on different bacterial hosts.

We also found an interesting case of taxa-specific enzymes in lytic phages and prophages. It was the sipho-morphotype exclusive depolymerases - 5 Webervirus proteins in K2 group 1 and 4 in K62 group 1 (at least one confirmed as a Webervirus protein). Of course, we do not exclude the possibility that this conclusion is based solely on a few representative proteins described so far. As new phages are discovered, it may turn out that this group will expand to include representatives from other taxa as well. Nevertheless, it is unlikely to be a coincidence that identical enzymes have been isolated from phages originating from distant environments and diverse bacterial hosts.

### What is the specificity range of identified depolymerases toward Klebsiella capsular types?

In theory, the ability to degrade multiple capsular types could be largely attributed to flexible or double substrate-binding regions (carbohydrate-binding motifs) involved in the interactions with bacterial polysaccharides. Broad active enzymes often contain adaptable catalytic domains that allow them to interact with diverse polysaccharides (Leiman & Molineux, 2008).

Nevertheless, our catalog shows double K-type tropism only for CPS sharing the same main unit (K2/K13; K22/K37; K30/K69 and K57/K68, Figure 1), meaning that enzymatic RBPs are very specific towards the substrate, not tolerating structurally diverse polysaccharides. The only exception we observed was the KLEO13gp09 depolymerase, able to degrade 3 substrates (K22/K37/KL111), but so far, the sugar composition of KL111 CPS is still unknown.

### Can we predict their capsular tropism based on amino acid sequence or structure prediction?

The structure-based catalog of *Klebsiella* capsule depolymerases showed that the domain composition of enzymes with different capsule tropisms may be the same. Subtle spatial variations, primarily localized in the C-terminus and conformational differences in the active sites, appear to facilitate recognition of distinct glycosidic linkages and sugar residues. Additionally, specific structural motifs or flexible loop regions adjacent to the active site may adapt to accommodate diverse capsular architectures. Collectively, these features contribute to the broad specificity and functional versatility observed among depolymerases encountered among members with the same structural class (Figure 2). These findings highlight the complexity of enzyme-substrate interactions and suggest that even minor structural modifications can have significant functional implications on substrate specificity. We also observed high structural dissimilarities between most of the C-terminal domains, suggesting their key role in the specific recognition of particular CPS (Supplementary Material 1). Therefore, as long as the database of proteins with experimentally confirmed specificity remains limited, the prediction of substrate specificity is uncertain except in cases where amino acid sequence identity to a characterized enzyme is exceptionally high.

Looking at the structural versatility with a particular class and between the five created classes, we clearly see that each protein representing separate K-specificity is characterized by its unique and well-defined structural conformation. This underscores the extensive variability and intricate nature of both the structural architecture and substrate specificity of these enzymes.

### What drives the diversity of phage depolymerases: domain shuffling or point mutations?

In theory, both domain shuffling and point mutations contribute to modular enzymes’ diversification, with domain shuffling playing a more prominent role in generating novel functional variants (Knecht et al. 2020). The modular architecture of depolymerases allows for recombination events, particularly involving the catalytic and substrate-binding domains (Latka et al. 2021, Latka et al. 2025). Domain shuffling between N-terminus and Central+C-terminal parts can result in the acquisition of external enzymes with novel properties, exchanging the host range(Steven et al. 1988, Stummeyer et al. 2005).

But there is a big question mark if the evolution enables rapid adaptation to new capsule types by separate enzymatic or C-terminus domain exchange? In contrast to recombinations, point mutations serve to fine-tune enzymatic properties within existing domain frameworks. Mutations in key residues can modulate activity, stability, and specificity, thereby affecting phage infectivity. This is what we found in K22/K37/KL111 and K25/KL119 targeting depolymerases with an almost identical structure of the central enzymatic domain and conserved catalytic pocket, but different distribution of negatively charged moieties responsible for sugar binding.

The diversity of the capsule as phage receptor (K swap/modifications) drives the need for such diversification. Ultimately, the interplay between domain-level recombination and residue-level mutations ensures a dynamic and adaptable depolymerase repertoire (Hampton et al. 2020, Latka et al. 2021).

A similar strategy was reported on LPS-specific depolymerases found in *Salmonella* and *E. coli* phages, which present “*an evolutionary model in which phages employ two distinct strategies to adapt their tailspikes to the substrate diversity represented by bacterial LPS. First, they retain a conserved and robust protein fold while modifying key active site residues to alter substrate specificity. Second, they incorporate foreign domains, already functionally optimized for a specific LPS serotype, into the tailspike structure”* (Leiman & Mollineux, 2008).

### Can novel depolymerases be engineered to broaden their substrate range for therapeutic or diagnostic applications?

Beyond their role in phage biology, depolymerases have significant therapeutic and diagnostic applications. Due to depolymerase high specificity they hold potential for diagnostic use, enabling the rapid serotyping of bacterial isolates (Li et al. 2021). Their substrate specificity could be leveraged to differentiate bacterial strains based on capsular composition.

In the therapy approach, depolymerases are used for biofilm degradation, for bacteria sensitization to capsule-independent phages, and to increase bacterial susceptibility to immune recognition and clearance upon capsule degradation (Majkowska-Skrobek et al. 2016, 2018 & 2021, Wang et al. 2023, Topka-Bielecka et al. 2021, Liston et al. 2018). Capsule degradation also improves antibiotic penetration, making previously resistant strains more vulnerable to standard treatments (Wu et al. 2019). Therefore, the emerging tools for protein engineering could broaden the depolymerase substrate range (for therapeutic purposes) as a promising approach for combating multidrug-resistant bacterial infections. The synthetic modular enzymes have been widely studied for phage endolysins, where combining domains from different enzymes or directed mutagenesis, researchers could create chimeric enzymes with expanded or altered specificity (Gerstmans et al. 2018, Binte Muhammad Jai et al. 2020). Structural insights and computational modeling guide the rational design of such enzymes, allowing for targeted modifications.

There was also a series of experiments done by our team regarding chimeric Klebsiella phage RBPs. The domains of phage KP32 depolymerase, which naturally target the K3 capsular type, were successfully replaced with domains from K63- and K21-specific phage RBPs, resulting in the specificity swap (Latka et al. 2021). The functional autonomy of the C-terminal domains was also confirmed, stressing their role in substrate binding and enzymatic activity (CBM) and trimerization (LEC) (Latka et al. 2025). In addition to domain swapping and modular grafting, genetic miniaturization of depolymerases has emerged as an effective strategy for simplifying enzyme architecture while retaining catalytic function. A truncated version of the KP34gp57 depolymerase was constructed by removing the N-terminal conserved peptide and C-terminal trimerization modules, yielding a monomeric active “mini-enzyme” composed solely of the central catalytic β-helix and the β-barrel insertion domain. Notably, this approach is only applicable for depolymerases with an intrasubunit catalytic site since depolymerases with an intersubunit catalytic cleft lose their activity upon loss of the trimeric state (Maciejewska et al. 2023, Latka et al. 2025).

These results provide strong experimental support for the concept that phage depolymerases function as modular platforms, amenable to precise genetic manipulation for the generation of tailored anti-capsular agents. Nevertheless, uncovering the underlying design rules will require a larger body of experimental data, complemented and accelerated by AI-driven analytical tools. Moreover, analyzing the depolymerase catalog, we can see that modifying or broadening the substrate specificity of these enzymes will be very challenging.

In summary, through the comprehensive cataloging and functional validation of 105 phage-derived depolymerases targeting a large panel of CPS types and the creation of *Klebsiella* capsule depolymerase catalog, this study can lay the groundwork for deeper exploration of phage-host interactions, evolutionary dynamics, biological functions, and the translational potential of Klebsiella phage depolymerases in therapeutic and biotechnological applications.

## Materials and Methods

### In silico identification of depolymerase

The following criteria were used for putative depolymerase identification: (1) the protein must be longer than 500 amino acid residues; (2) the protein must be annotated as tail fiber/tail spike protein in the searched database; (3) the protein must show homology to domains annotated as lyases (e.g. hyaluronate lyases (hyaluronidases), pectin/pectate lyases, alginate lyases, K5 lyases) or hydrolases (e.g. sialidases, rhamnosialidases, levanases, dextranases or xylanases), recognized by BlastP (Altschul et al. 1990), HMMER (Finn et al. 2011) or HHpred (Zimmermann et al. 2018); (4) a typical β-helical structure (Latka et al., 2019) should be predicted by AlphaFold3 (Abramson et al. 2024). BlastP was performed against the non-redundant protein sequences (nr) database using standard parameters (expect threshold: 0.05; word size: 5; matrix: BLOSUM62; gap costs: existence 11, extension 1; conditional composition score matrix adjustment). HMMER was used in quick search mode against Reference Proteomes with significance E-values: sequence 0.01 and hit 0.03. HHpred homology detection prediction was run using the PDB_mmCIF70 database and the following parameters (MSA generation method: HHblits=>UniRef30; maximum number of MSA generation iterations: 3; E-value cutoff for MSA generation: 1e-3; minimal sequence identity of MSA hits with query (%): 0; minimal coverage of MSA hits (%): 20; secondary structure scoring: during_alignment; Alignment Mode: Realign with MAC: local:norealign; MAC realignment threshold: 0.3; number of target sequences: 250; minimal probability in hitlist (%): 20). Protein clustering was performed by BlastP with 80% sequence coverage and 50% sequence identity (Altschul et al. 1990). Structural predictions were obtained using AlphaFold3 (Abramson et al. 2024), followed by visualization in PyMOL (Schrödinger, LLC, v3.0). The depolymerases derived from the prophages were identified in the KASPAH and KLEBPAVIA *Klebsiella spp*. genome collections in our previous study (Otwinowska et al. 2025). Prophage genome detection was performed using VirSorter2 (Guo et al. 2021) and PhySpy (Akhter et al. 2012). BlastN (Altschul et al. 1990) was used for genome clustering with 90% sequence coverage and 50% sequence identity. Gene functional annotation was executed (30% sequence coverage and 95% sequence identity) with COG_KOG (Tatusov et al. 2000), PDB (wwPDB consortium, 2019), PFAM (Mistry et al. 2021), PHROGS (Terzian et al. 2021), and UNICLUST (Mirdita et al. 2017) databases. The morphotypes of prophages were assigned using the VIRFAM tool (Lopes et al. 2014).

### Bacterial Strains

*Klebsiella spp*. K-type panel used in this study originate from the Collection de l’Institut Pasteur (CIP), Paris, France; The National Collection of Type Cultures (NCTC), the UK Health Security Agency (UKHSA); collection of the Department of Pathogen Biology and Immunology (Wroclaw, Poland); and the *Klebsiella* Acquisition Surveillance Project at Alfred Health (KASPAH) (Melbourne, Australia) (Gorrie et al. 2022) provided by Kath Holt the London School of Hygiene and Tropical Medicine (LSHTM) Department of Infection Biology, London, UK. The full *K. pneumoniae* K-type panel includes 134 strains representing 119 distinct serotypes (Supplementary Table S1). All bacteria were cultured in Tryptone Soya Broth or Agar (TSB or TSA, Oxoid, Thermo Fisher Scientific, Waltham, MA, USA) at 37 °C and stored at -70°C in a 20% glycerol-supplemented Trypticase Soy Broth (TSB, Becton Dickinson, and Company, Cockeysville, MD, USA).

The *Escherichia coli* strains (Thermo Fisher Scientific, Waltham, MA, USA) used for recombinant protein production: One Shot™ TOP10 for plasmid propagation and One Shot™ BL21 Star™ (DE3) for recombinant protein expression, were cultured in LB broth, Miller (Luria-Bertani) (Difco, Becton, Dickinson and Company (BD), Franklin Lakes, NJ, USA) at 37 °C supplemented with 100 µg/mL ampicillin (IBI Scientific, Dubuque, IA, USA).

### Cloning, expression, and purification

The recombinant depolymerases were obtained through two different cloning approaches. A subset of genes was molecularly cloned in-house using the *Champion pET101* cloning/expression vector (Thermo Fisher Scientific, Waltham, MA, USA) with a C-terminal His tag (6x) and propagated in *E. coli* One Shot™ TOP10 strain. The remaining constructs were synthesized and prepared in expression vectors by commercial service GeneUniversal (Newark, DE, USA). The accuracy of all clones was verified by sequencing (Genomed, Warsaw, Poland) using the specific primers (Supplementary Table S3). Following transformation into *E. coli* One Shot™ BL21 Star™ (DE3), vector-bearing cells were cultured in 500 mL of LB broth supplemented with 100 µg/mL ampicillin at 37 °C with agitation until reaching an optical density (OD600nm) of 0.5–0.6. Recombinant protein expression was induced at 18 °C for 16 h by adding isopropyl-β-D-thiogalactopyranoside (IPTG) to a final concentration of 1 mM. The cultures were then harvested by centrifugation (8000 × g, 15 min, 4 °C), and the resulting pellets were resuspended in lysis buffer (500 mM NaCl, 20 mM NaH_2_PO_4_, pH 7.4). The exact expression conditions for each protein are provided in Supplementary Table S3. After sonication, the lysates were centrifuged (16,000 × g, 30 min, 4 °C), and the supernatants were collected and filtered through a 0.22 µm filter (Sarstedt AG & Co., Numbrecht, Germany). His-tagged proteins were purified using Bio-Scale Mini Profinity IMAC cartridges (Bio-Rad, Hercules, CA, USA) in combination with an FPLC system (Bio-Rad, Hercules, CA, USA). The column was equilibrated with washing buffer (500 mM NaCl, 20 mM NaH_2_PO_4_, pH 7.4), followed by sample loading and washing with 32 column volumes of the same buffer. The proteins were subsequently eluted using an elution buffer (500 mM NaCl, 20 mM NaH_2_PO_4_, 500 mM imidazole, pH 7.4). Some recombinant proteins derived from Klebsiella prophages were prepared by a commercial company (GenScript Biotech, Rijswijk, Netherlands) with the protocol details provided in Supplementary Table S3. Finally, protein concentrations were determined spectrophotometrically (NanoPhotometer NP80, Implen GmbH, Schatzbogen, München, Germany) using a molar extinction coefficient calculated for each protein with the ProtParam tool (Gasteiger et al. 2005).

### SDS-PAGE protein analysis

Sodium dodecyl sulfate-polyacrylamide gel electrophoresis (SDS-PAGE) was conducted following the method of Laemmli (Laemmli, 1970) using a 12% polyacrylamide gel. Next, the protein samples were mixed with Laemmli buffer (Bio-Rad, Hercules, CA, USA) in a 3:1 ratio, respectively, and analyzed in SDS-PAGE. The protein samples were heated at 99 °C for 7 min. Molecular weight standard Precision Plus Protein All Blue (10-250 kDa) (Bio-Rad, Hercules, CA, USA) was used. The protein bands were visualized by GelCode Blue staining (Thermo Fisher Scientific, Waltham, MA, USA).

### Enzymatic functionality assays

Detection of enzyme activity was performed by spot assay on the *K. pneumoniae* serotype panel listed in Supplementary Table S1. Bacteria were grown until reaching an optical density (OD_600nm_) of 1.0, and then,1 ml of bacterial culture was directly transferred onto TSA plates. After drying, 10 µL of a serial two-fold dilution of recombinant enzyme in phosphate-buffered saline (PBS; 137 mM NaCl, 2.7 mM KCl, 10 mM Na_2_HPO_4_, 1.8 mM KH_2_PO_4_, pH 7.4). 10 µL of PBS buffer (negative control) was applied to the bacterial lawn in spots. Following an overnight incubation at 37 °C and an additional 24 h incubation at room temperature, plates were examined for the presence of clear zones (halo). The minimum concentration of protein required to produce a detectable halo on the bacterial lawn was determined by serial dilution of the enzyme to establish the minimal halo-forming unit (MHFC).

### The structural classification of depolymerases

The structural classification of depolymerases was established by delineating three distinct domains: the N-terminal domain, the central β-helical enzymatic domain, and the C-terminal domain (Latka et al. 2019). Structural models were generated using AlphaFold3 (Abramson et al. 2024) and subsequently visualized with PyMOL (The PyMOL Molecular Graphics System, Version 3.0, Schrödinger, LLC). Domain boundaries were assigned based on the analysis of secondary structural elements within the predicted structures, following the classification criteria established by Huang et al. (2024) (Figure 4).

**Figure 4.**
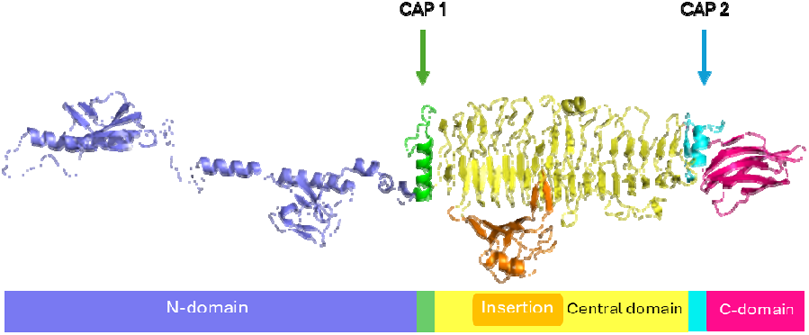
Identification and delineation of depolymerase domains based on Huang et al. 2024.

The N-terminal domain was defined as the region preceding the terminal α-helix (cap 1), which serves as the structural demarcation between this domain and the central enzymatic domain. The central β-helical enzymatic domain was identified as the region initiating from cap 1 and adopting a right-handed parallel β-helix fold, a characteristic structural motif of depolymerases. In cases where an insertion domain was present within the β-helical core, it was distinguished by the occurrence of antiparallel β-sheets and α-helices forming an extended protruding loop. The termination of the central domain was defined by the presence of cap 2, the final α-helix of the β-helical fold. In the absence of cap 2, the domain boundary was delineated at the loop structure preceding the C-terminal β-sandwich domain. The C-terminal domain was assigned to the region downstream of cap 2 (or the corresponding loop structure in its absence) and was characterized by the presence of a β-sandwich fold. In some cases, more than one C-terminal domain could be distinguished per protein: a) both adapting β-sandwich folds separated by loops or linkers; b) the ultimate C-terminal β-sandwich was preceded by an elongated β-strand-rich motif, not forming a right-handed parallel β-helix fold, which was treated as a separate domain. In other cases ultimate C-terminal domain was rich in alpha helices, forming a visually separated domain.

As an attempt to assign additional functions, C-terminal domain(s) or atypical parts of β-helix were selected from the 3D structures and analyzed with the DALI server (Holm et al. 2023) using PDB search for domains previously detected as tailspikes. In particular, ‘CBM’ (carbohydrate-binding module), ‘LEC’ (lectin-like domain), ‘neuraminidase’, and ‘colanidase’ were of interest. Additionally, some regions between the β-helix and the C-terminus were found to be similar to the ‘tail fiber protein’. If at least the Z-score was > 2, the DALI hit was reported (however, since RMSD was not always below 3.0 Å, the results are not reported in detail). Proteins whose C-terminus was rich in alpha-helices were analyzed with PHYRE2.2 (Powell et al. 2025) in search of a C-terminal ‘chaperone’. The delineation was made based on BlastP analysis run with standard parameters (as listed above), where the ‘peptidase’ was detected and treated as a ‘chaperone’ start.

### Sequence and structural alignment of proteins

We used BLASTP (v2.16.1+, Altschul et al. 1990) with default settings to compare protein sequences. Multiple sequence alignments were done using MAFFT (v7.525; Katoh & Standley, 2013), and a phylogenetic tree was built with IQ-TREE (v2.4; Nguyen et al. 2015), applying the JTT+G substitution model. To evaluate the reliability of the branches, we ran 1000 ultrafast bootstrap replicates. The resulting tree was visualized through the Interactive Tree of Life web server (iTOL v7; Letunic & Bork, 2024). For comparative analysis of capsular polysaccharide synthesis (CPS) loci across different genomes, we used Clinker (v0.0.31; Glichrist & Chooi, 2021), which allowed for fast visualisation of gene clusters and their synteny. Protein structures were compared using Usalign (v2024.07.30; Zhang et al. 2022), and we carried out all structural alignments and visualisations in PyMOL (v3.0; Schrödinger, LLC, 2010). We also used PyMOL, along with the APBS plugin (v3.4.1), to calculate and display electrostatic surface potentials. Default PDB2PQR preparation was used to add hydrogens and assign partial charges and atomic radii. Electrostatic potential maps were calculated using APBS with a grid spacing of 0,50 Å. Molecular surfaces were then visualised in PyMOL using a color range of +/-5.00 kT/e. To examine evolutionary conservation of amino acid residues, we used the ConSurf webserver (v2025; Glaser et al. 2003, Landau et al. 2005, Ashkenazy et al. 2010, Ashkenazy et al. 2016). Three-dimensional depolymerase structures in PDB format were submitted, and conservation scores were calculated using default parameters. The server automatically retrieved homologous sequences, generated a multiple sequence alignment, and computed Bayesian residue-specific conservation scores. The number of homologs gathered for each protein was: KLEO13gp10 = 9, 184_43 = 49, KANgp232 = 62, KP24gp300 = 62, and S2-2 = 52. These scores were then mapped onto the three-dimensional structures in PyMOL to highlight regions likely to be functionally important.

## Funding

This work was supported by Narodowe Centrum Nauki, Poland (Polish National Science Centre) in the frame of HARMONIA project UMO-2017/26/M/NZ1/00233 (AL, GMS, BM, FS, RB, ZDK), OPUS-LAP project UMO-2022/47/I/NZ1/01450 (MP, BM, SJ, MH, AL, YB, ZDK), SONATA Bis project UMO-2020/38/E/NZ8/00432 (AO, SO, SO, BM, RM) This work was financed by the Polish National Agency for Academic Exchange (JK, RM), the EMBO Installation Grant (JK, VRP, RM), the Agence Nationale de la Recherche for a FR-DE AMR Bilateral grant ANR-20-AMRB-0004-01 (RT) and AL was partially financed by the Research Foundation–Flanders, Belgium (FWO: 1240021N, 1251224N). GMS, BM, ZDK, JJB, RT were working in KLEOPATRA consortium project, supported by (UMO-2022/04/Y/NZ6/00123 Narodowe Centrum Nauki, Poland; ANR-22-AAMR-0006-06 from the Agence Nationale de la Recherche; 1KI2302A The Federal Ministry of Education and Research) under the framework of the JPIAMR – Joint Programming Initiative on Antimicrobial Resistance.

## Author contributions

The following contributions can be distinguished by different authors based on the CRediT (Contributor Roles Taxonomy) system:

Conceptualization: ZDK

Methodology: AO, SO, AL, GMS, BM, JK, VRP, FS, RB, Validation: AO, SO, RB, FS, YB, RJM, ZDK

Formal Analysis: AO, SO, AL, BM, ZDK

Investigation: AO, SO, AL, MP, GMS, BM, SJ, MH, FS, ZDK Resources: AL, MP, GMS, BM, JK, JJB, RT, SJJB, RJM

Data Curation: AO, SO, AL, BM, ZDK Writing – Original Draft: AO, SO, AL, ZDK Writing – Review & Editing: all authors Visualization: AO, SO, AL, ZDK Supervision: ZDK

Project Administration: ZDK

Funding Acquisition: ZDK, RJM, YB, JJB, RT, RB, AL

## Acknowledgements

We sincerely thank Kathryn E. Holt from Department of Infection Biology, London School of Hygiene & Tropical Medicine, London, United Kingdom/ Department of Infectious Diseases, School of Translational Medicine, Monash University, Melbourne, Victoria, Australia for generously providing access to the KASPAH strain collection, and we also acknowledge Institut Pasteur’s Biological Resources Center (CRBIP) (28 rue de Docteur Roux, Paris 15, France) for providing serotyped *Klebsiella spp*. CIP strains, which were instrumental in testing the capsular specificity of Klebsiella phages and their virion-associated depolymerases.

---

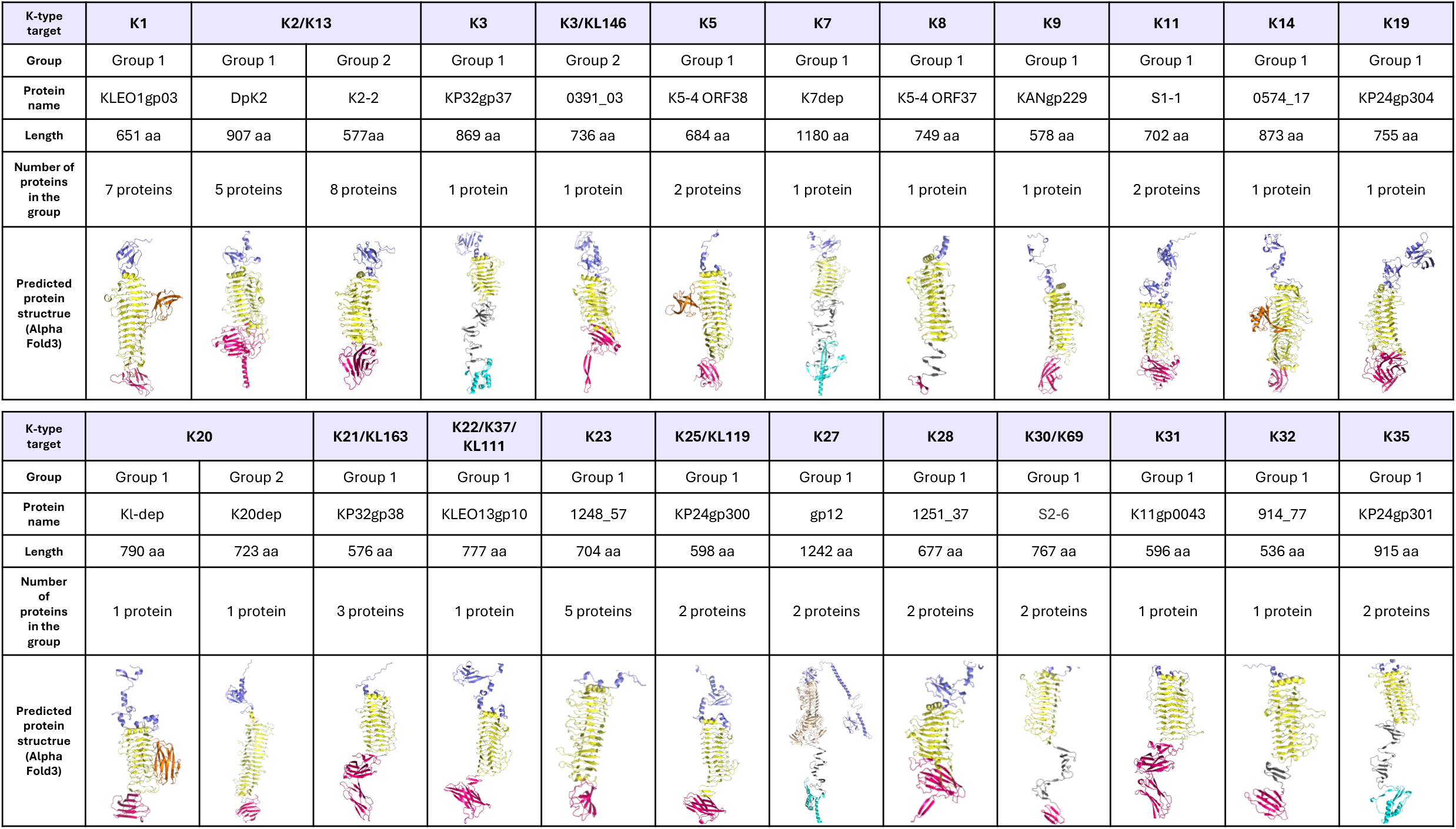

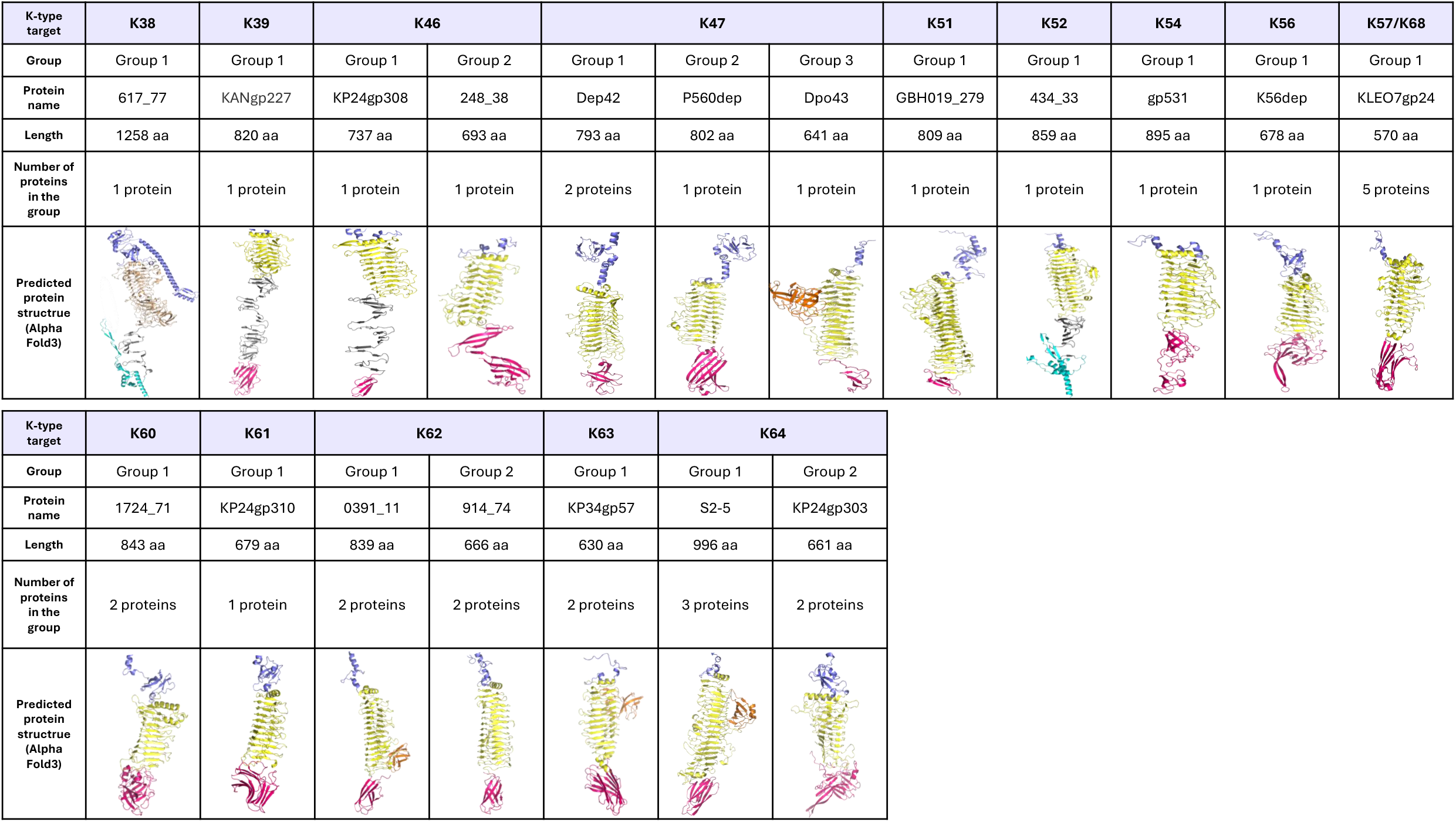

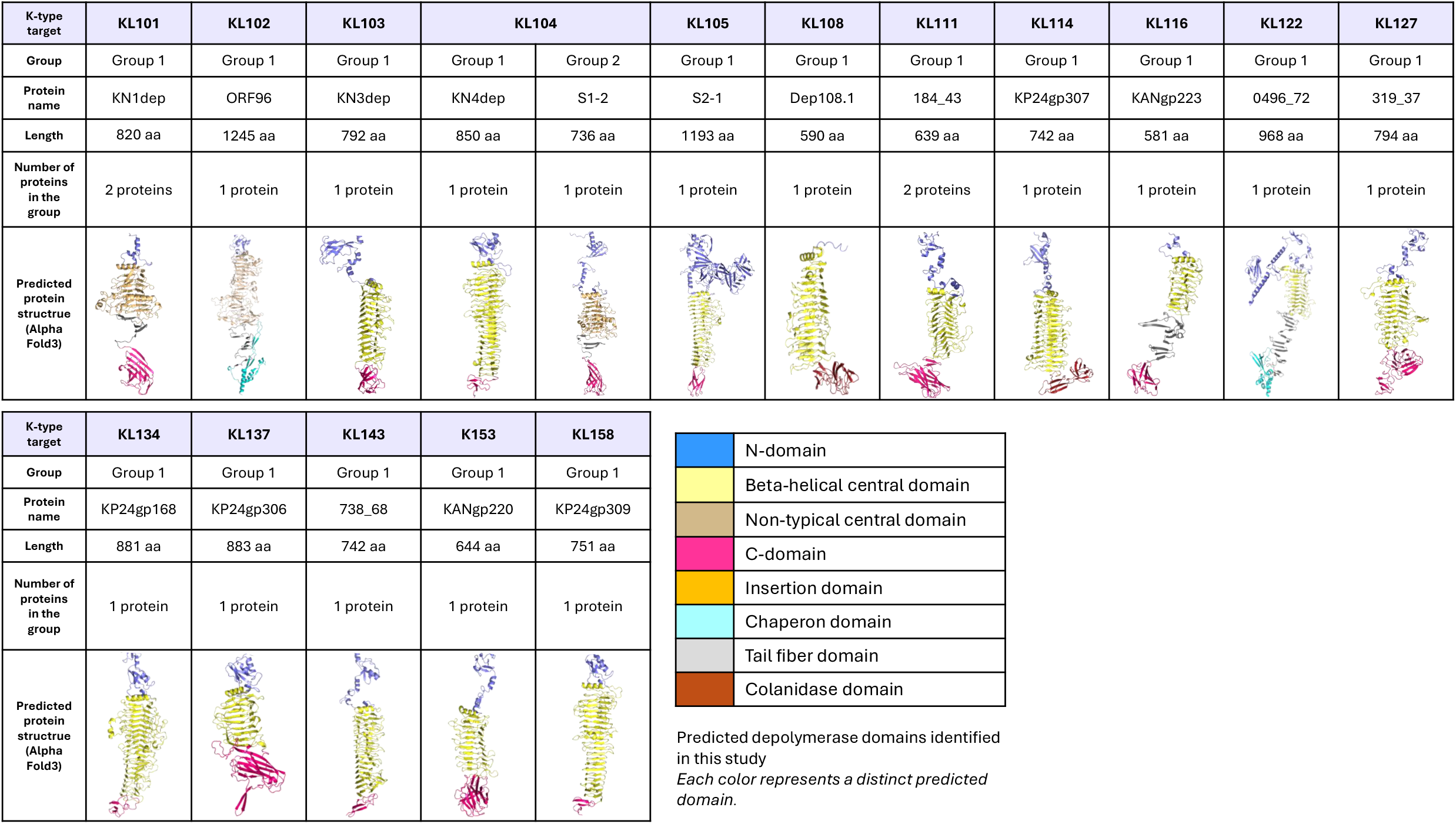

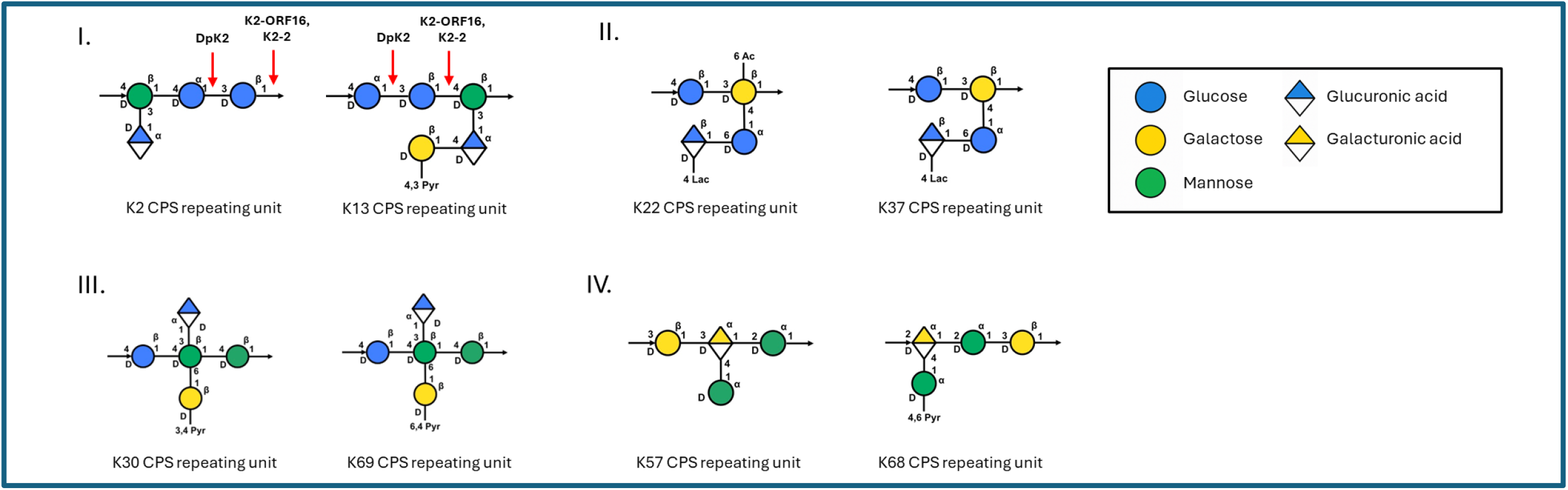

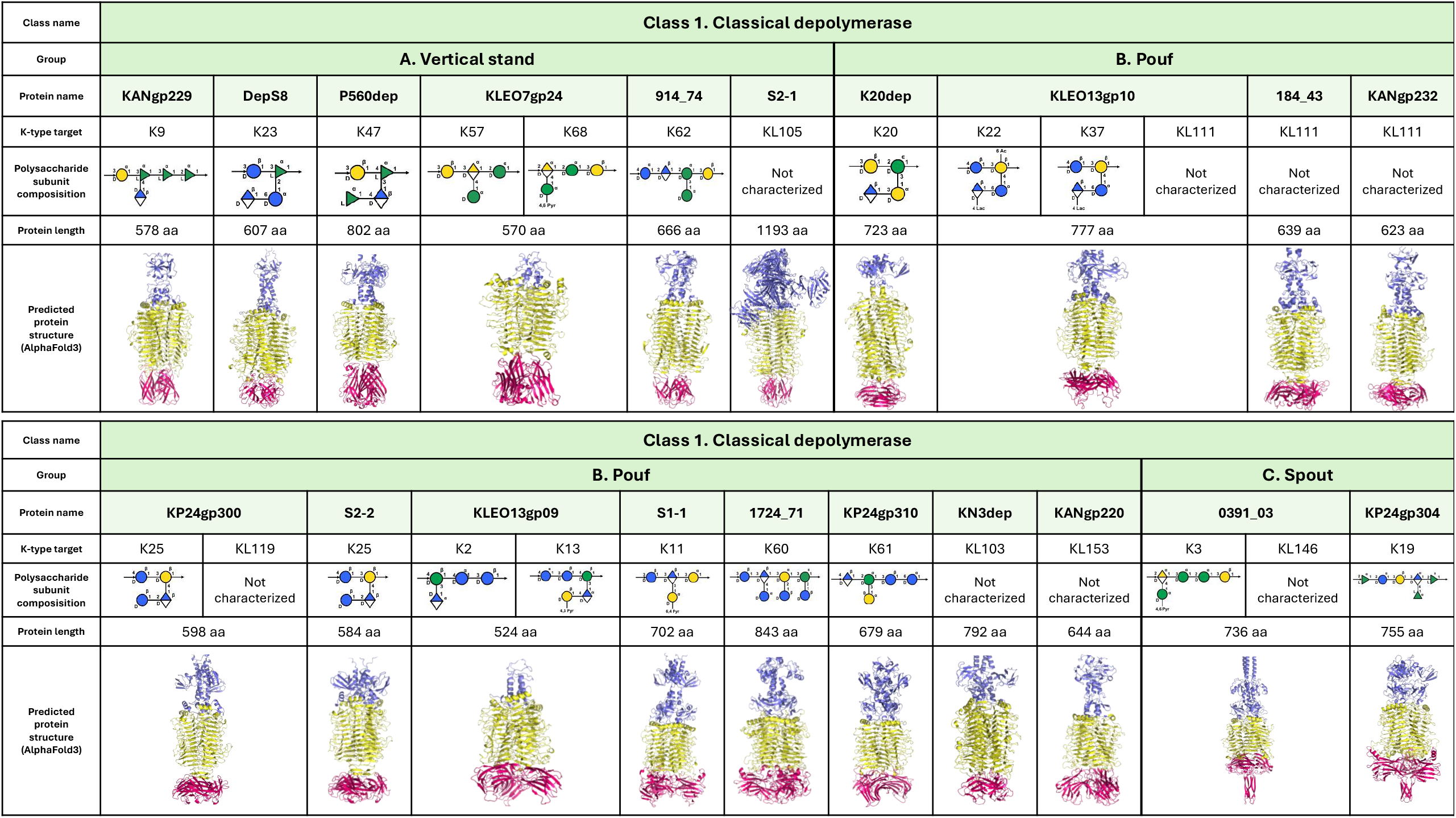

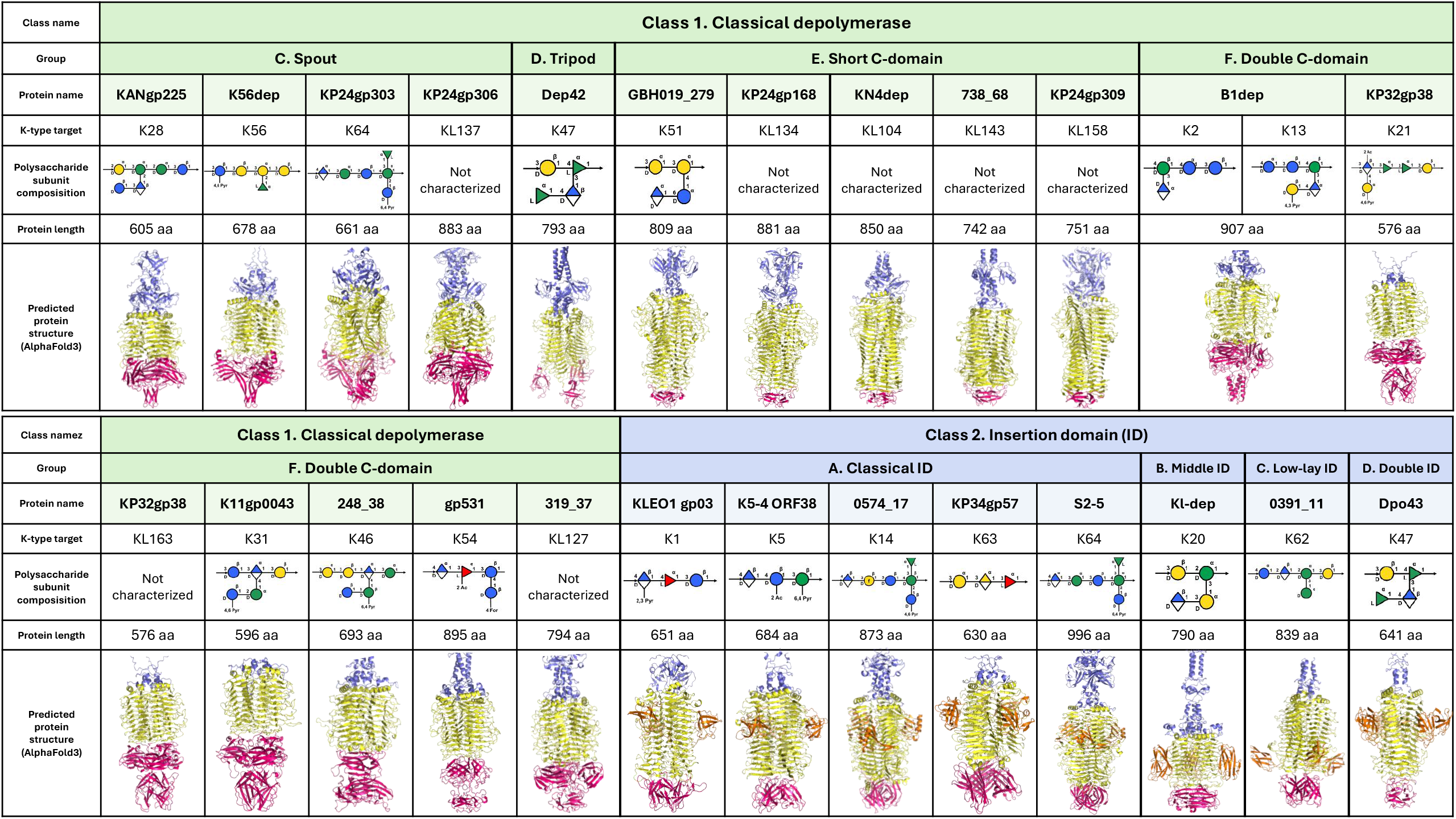

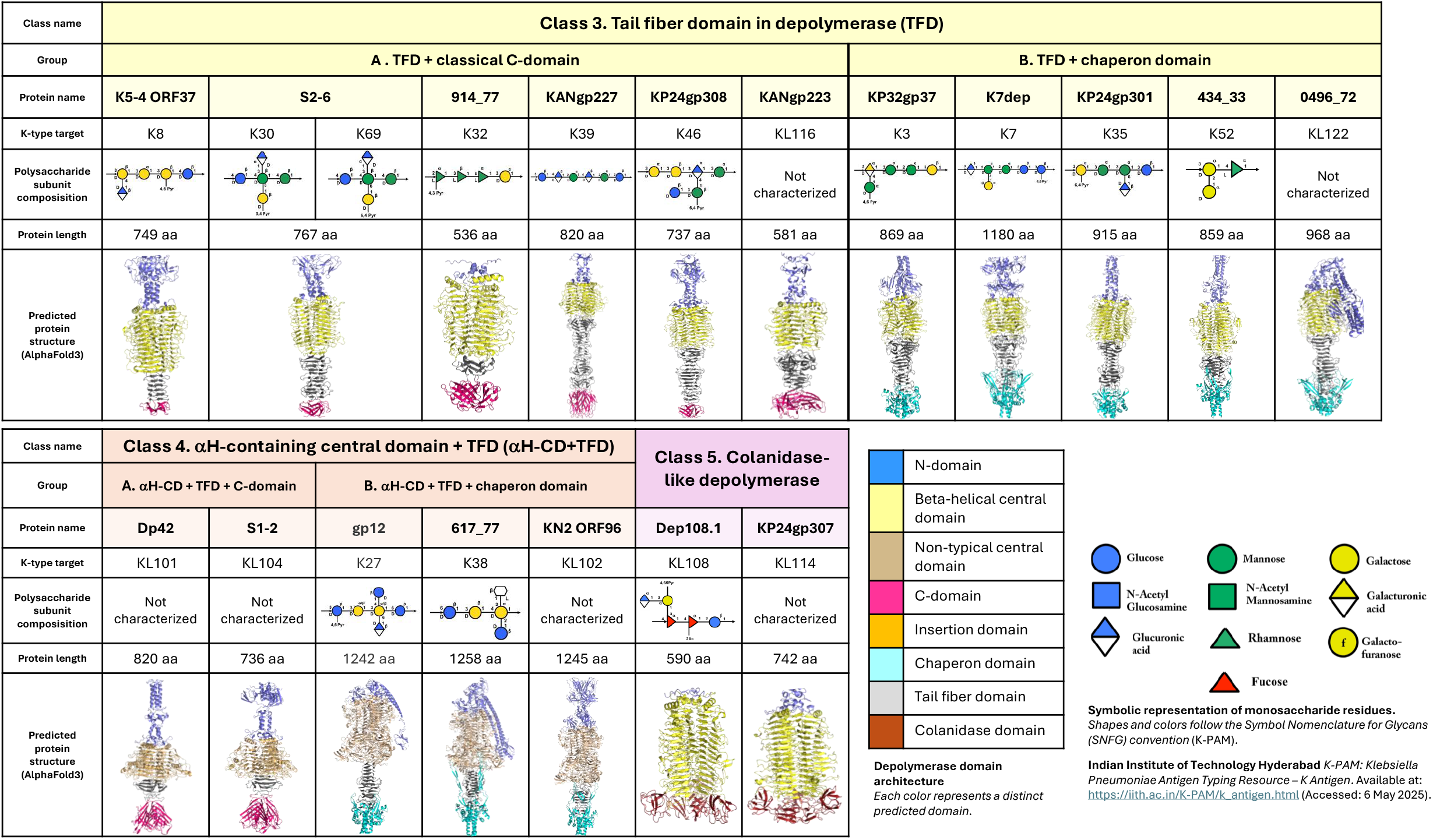

## Notes

### Competing Interest Statement

The authors have declared no competing interest.

