## Supplementary materials for "DepoCatalog: Mapping the Diversity of 105 Recombinant Klebsiella Phage Depolymerases Across Sequence, Structure, and Substrate Specificity"

### Depolymerase classification based on serotype specificity

**Domain architecture legend**

| | |
|---|---|
| | N-domain |
| | Beta-helical central domain |
| | Non-typical central domain |
| | C-domain (including i.e. CBM, LEC domains) |
| | Insertion domain |
| | Chaperon domain |
| | Tail fiber domain |
| | Colanidase domain |

**Symbolic representation of monosaccharide units used in polysaccharide structure diagrams**

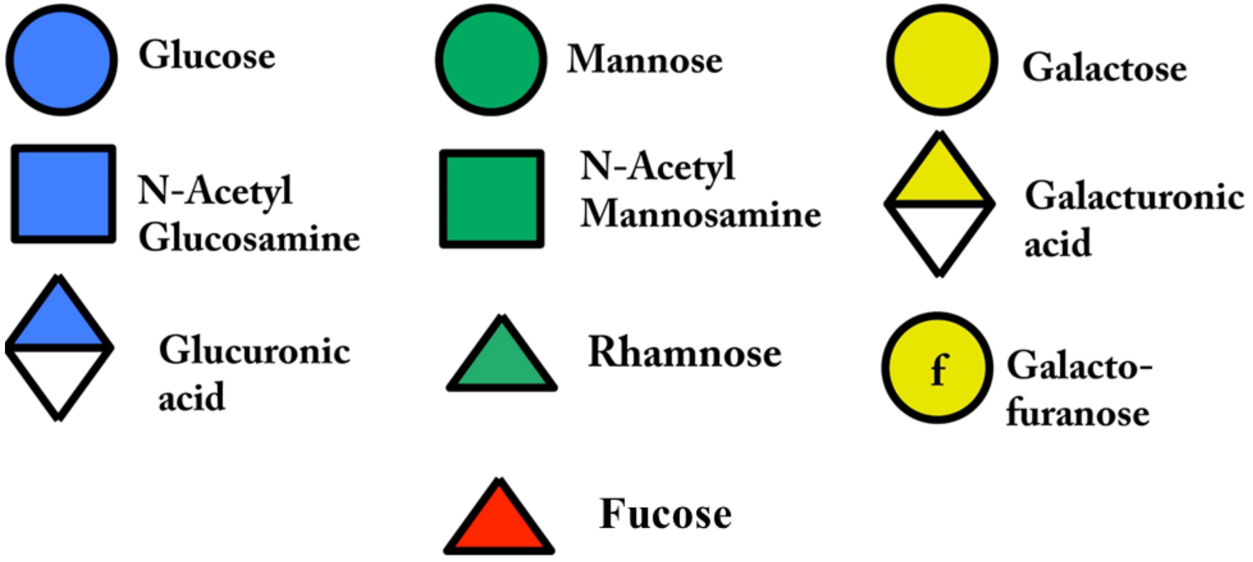


(*K-PAM: Klebsiella Pneumoniae Antigen Typing Resource – K Antigen*. Available at: https://iith.ac.in/K-PAM/k_antigen.html (Accessed: 6 May 2025).

# K1

**Cleavage site K1-ORF34**
(Tu et al. 2022)

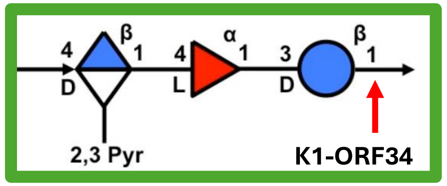


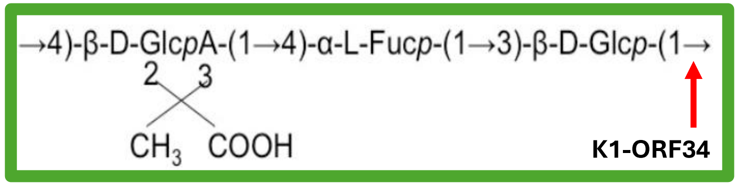


**Active Site Residues K1-ORF34**
**Tyr311,His 373 and Arg397**
(Tu et al. 2022)

| **K-type target** | **K1** | | | | | | |
|---|---|---|---|---|---|---|---|
| **Group** | Group 1 | | | | | | |
| **Protein name** | S2-4 | GBH001_056 | K1-ORF34 | gp47 | KLEO1gp03 | Kpv71_52 | gp09 |
| **Length** | 888 aa | 651 aa | 651 aa | 651 aa | 651 aa | 651 aa | 644 aa |
| **Number of protein in the group** | 7 proteins | | | | | | |
| **Predicted protein structure** | 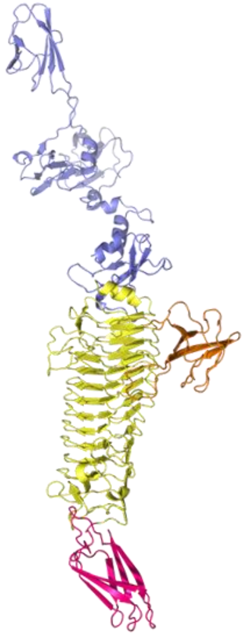 | 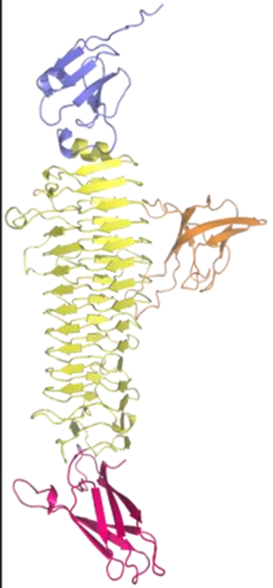 | 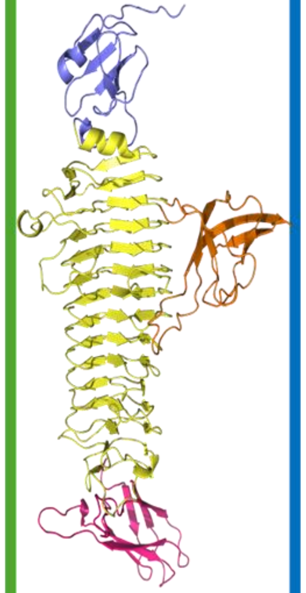 | 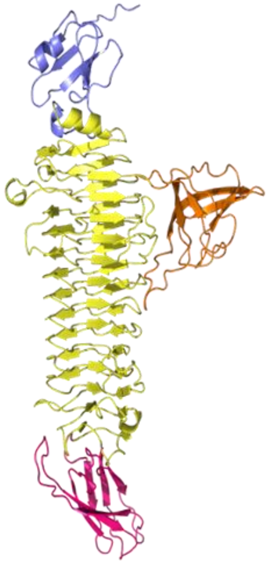 | 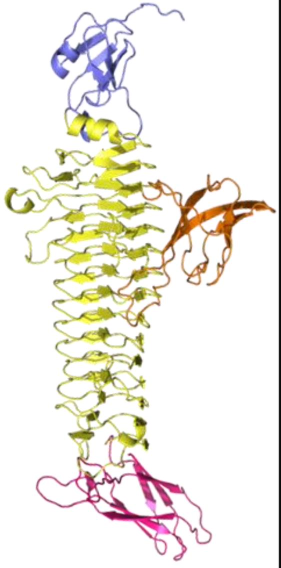 | 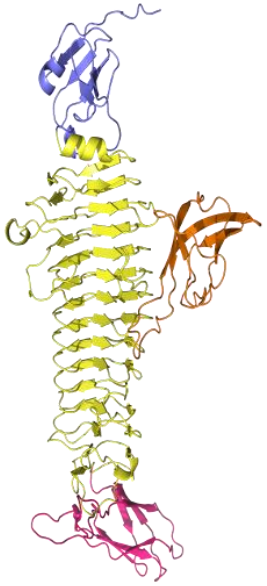 | 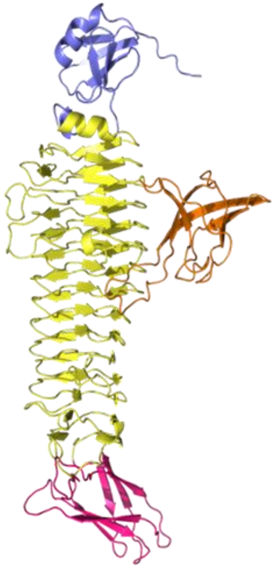 |

K1 group 1
TM-score;
Central + C-term

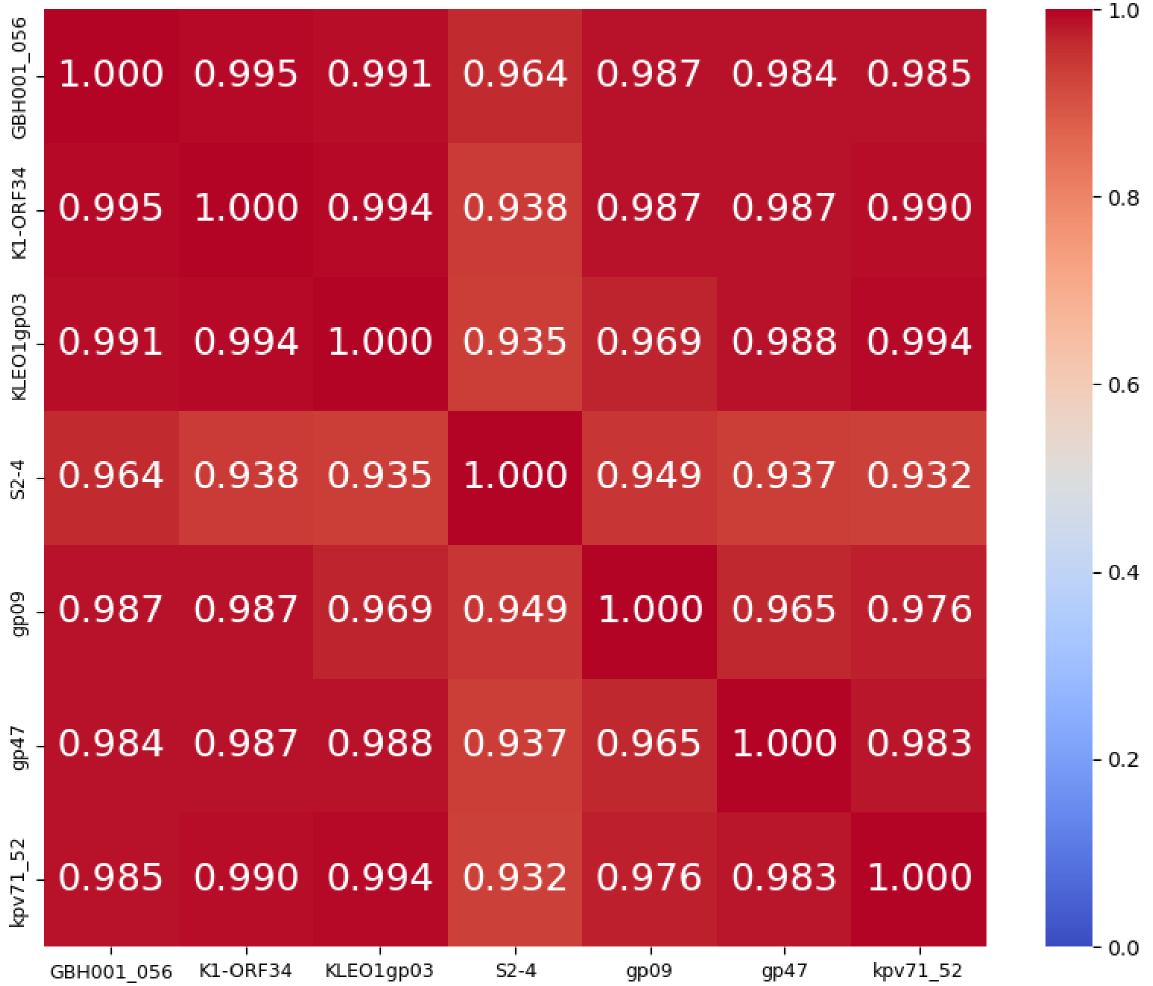


K1 group 1
%identity; cover ≥ 99%
Central + C-term

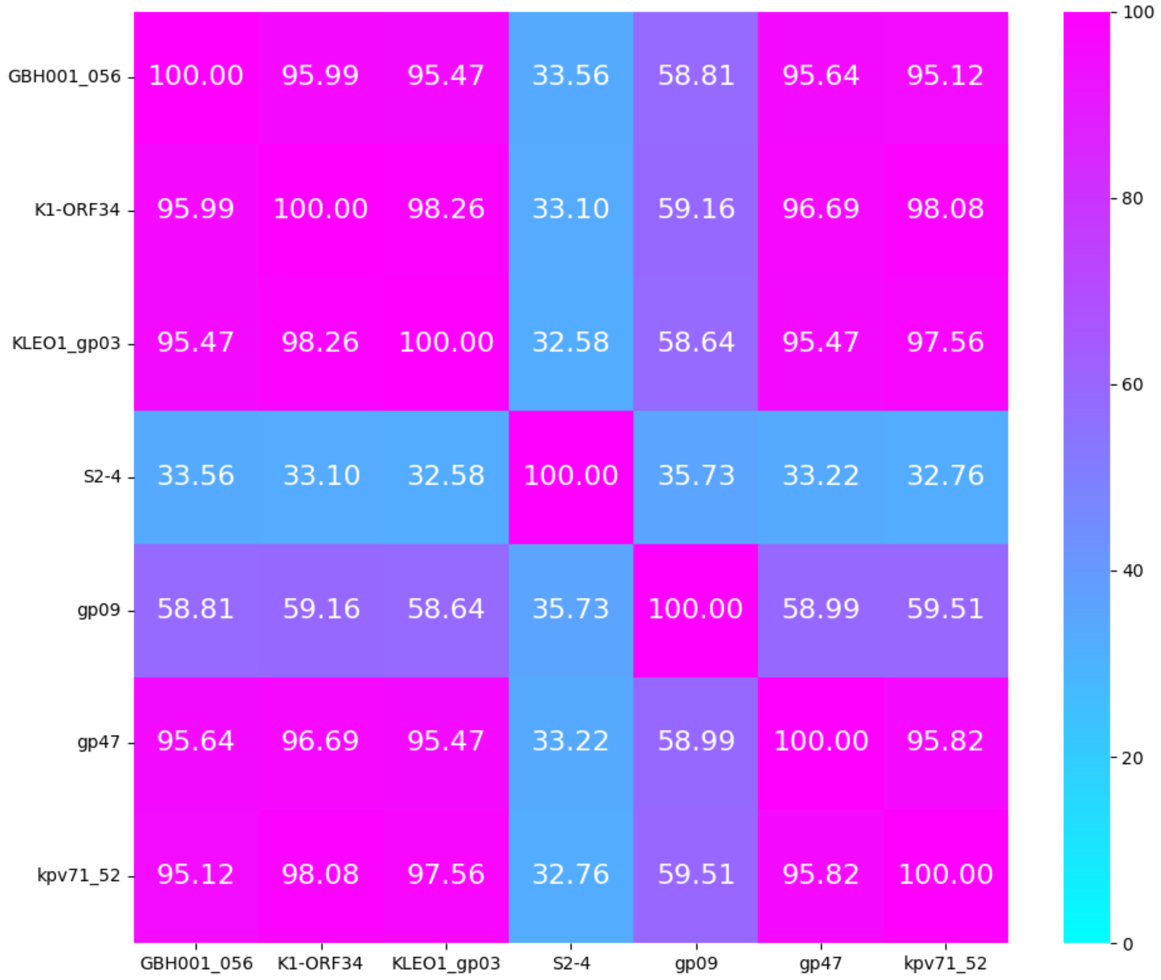

# K2

| K-type target | K2 | |
|---|---|---|
| **Group** | Group 1 | Group 2 |
| **Protein name** | DpK2 | KP24gp196 |
| **Length** | 907 aa | 660 aa |
| **Number of proteins in the group** | 5 proteins | 8 proteins |
| **Predicted protein structure** | 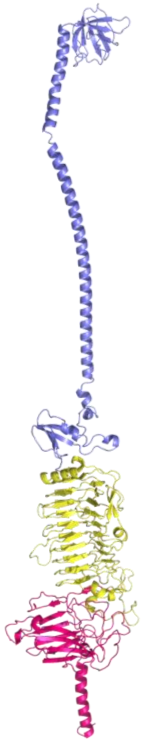 | 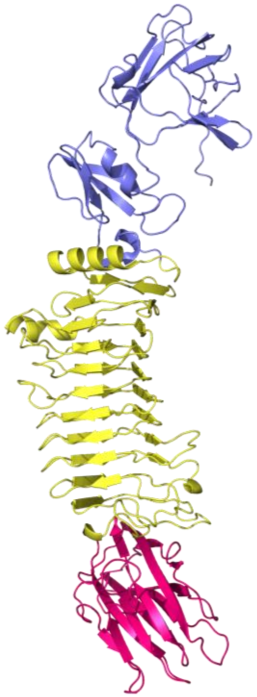 |

#### Cleavage sites
#### K2 depolymerases
(Dunstan et al. 2021, Lin et al. 2022, Ye et al. 2024)

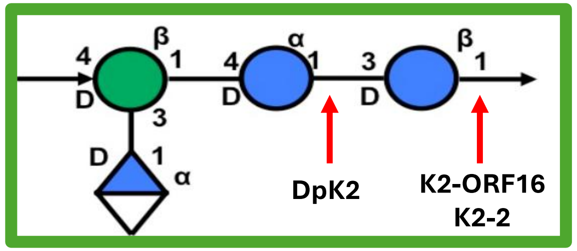


**K2**
CPS repeating unit

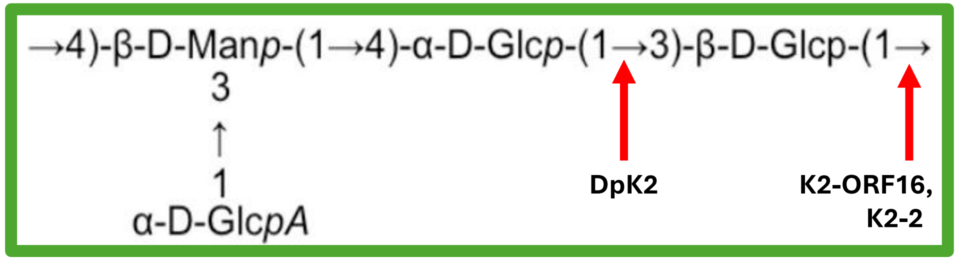

# K2

| K-type target | K2/K13 | K2* | K2* | K2* | K2/K13 |
|---|---|---|---|---|---|
| Group | Group 1 | | | | |
| Protein name | B1dep | DpK2 | Depo32 | BMacgp22 | gp81 |
| Length | 907 aa | 907 aa | 907 aa | 907 aa | 897 aa |
| Number of proteins in the group | 5 proteins | | | | |
| Predicted protein structure | 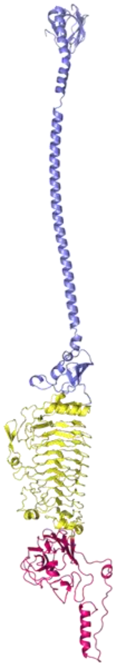 | 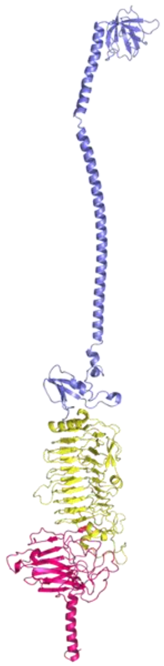 | 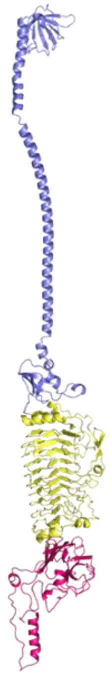 | 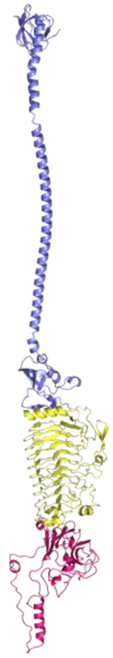 | 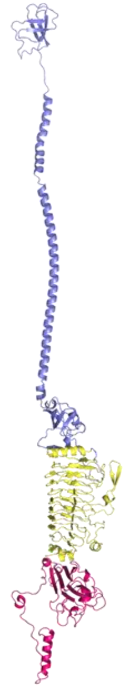 |

*protein was not tested on K13

**Cleavage site DpK2**
(Dunstan et al. 2021)

**K2**
CPS repeating unit

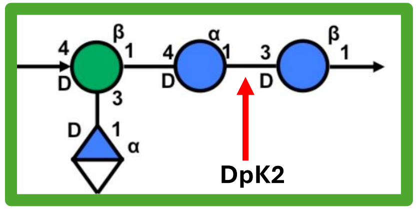


→4)-β-D-Man*p*-(1→4)-α-D-Glc*p*-(1→3)-β-D-Glc*p*-(1→
3
↑
1
α-D-Glc*pA*
DpK2

Active Site Residues **Depo32**
(Cai et al. 2023)
**Glu423, Glu545, Asp497, Asp546**

K2 group 1
TM-score
Central + C-term

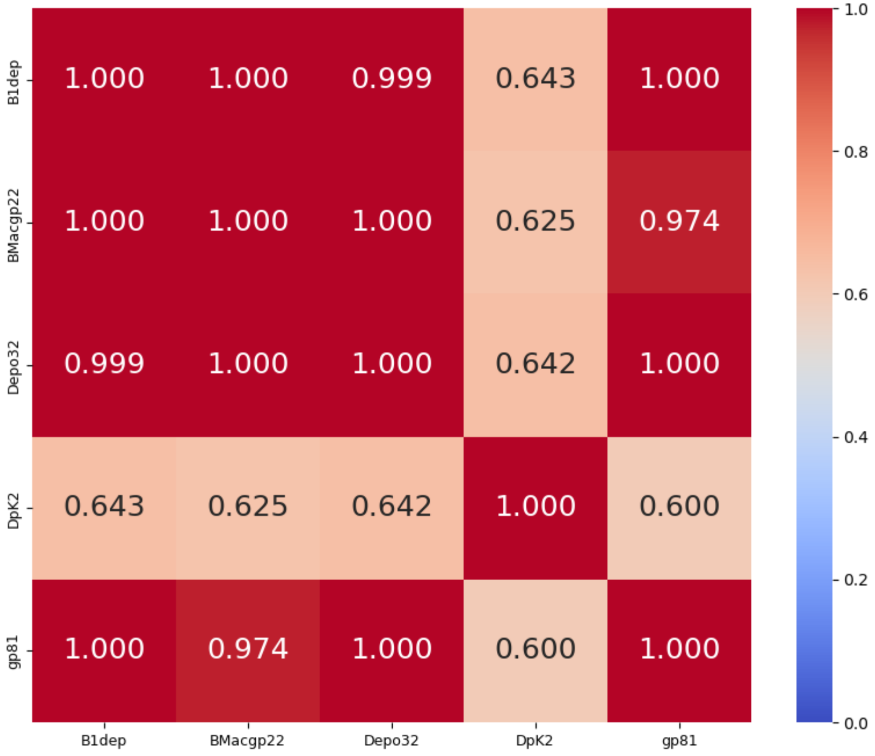


K2 group 1
%identity; cover ≥ 98%
Central + C-term

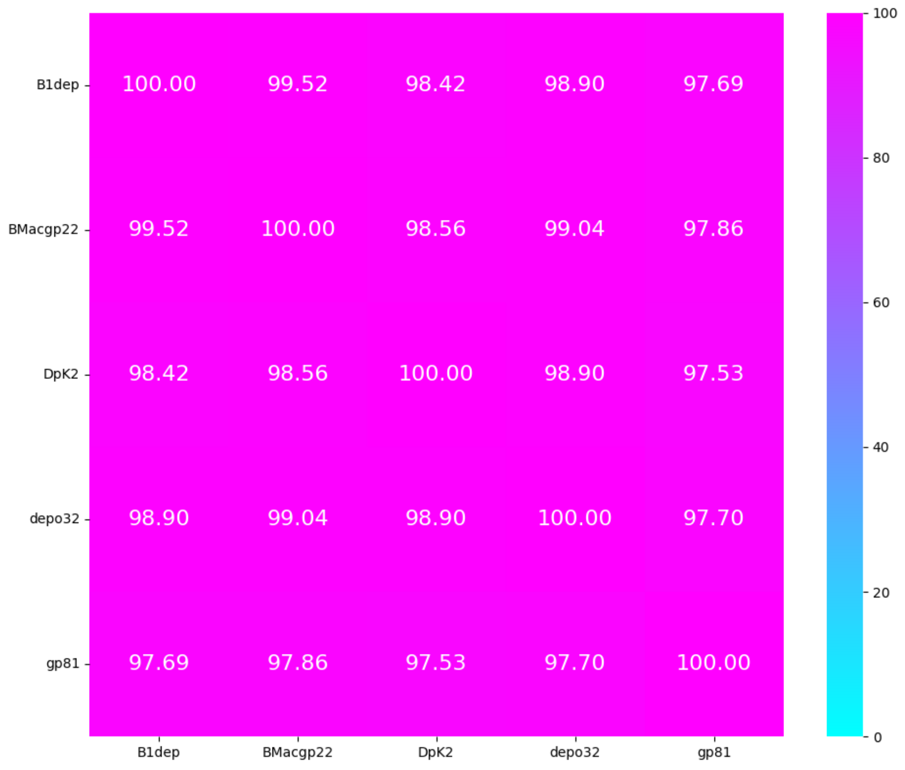


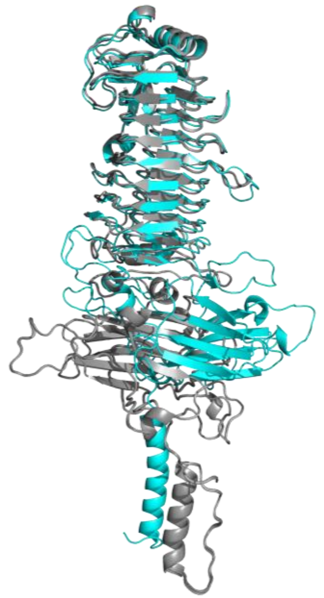

**Structural comparison of the central β-helix and C-terminal domains within the K2 group 1 depolymerases.** The cyan structure represents DpK2, while the grey structures correspond to B1dep, BMacgp22, gp81, and Depo32.

# K2

**Cleavage site**
(Lin et al. 2022, Ye et al. 2024)

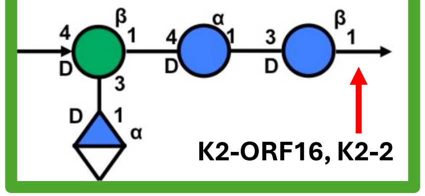


→4)-β-D-Man*p*-(1→4)-α-D-Glc*p*-(1→3)-β-D-Glc*p*-(1→
3
↑
1
α-D-Glc*p*A

K2-ORF16, K2-2

Active Site Residues **K2-2**
(Ye et al. 2024)
**Glu267, Glu323**

| K-type target | K2* | K2* | K2/K13 | K2/K13 | K2* | K2* | K2/K13 | K2/K13 |
|---|---|---|---|---|---|---|---|---|
| **Group** | Group 2 | | | | | | | |
| **Protein name** | NPatgp22 | K2-ORF16 | KP24gp196 | Kpv74_56 | Dep1979 | GBH038_054 | K2-2 | KLEO13gp09 |
| **Lenght** | 763 aa | 668 aa | 660 aa | 577 aa | 577 aa | 577 aa | 577 aa | 524 aa |
| **Number of proteins in the group** | 8 proteins | | | | | | | |
| **Predicted protein structure** | 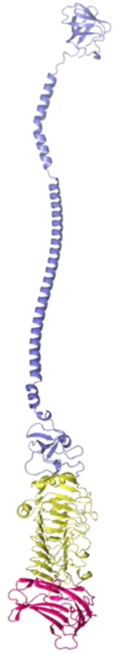 | 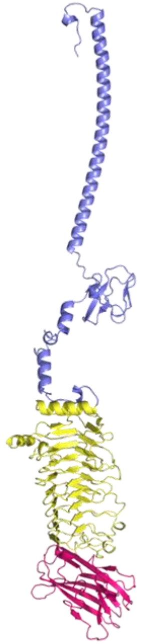 | 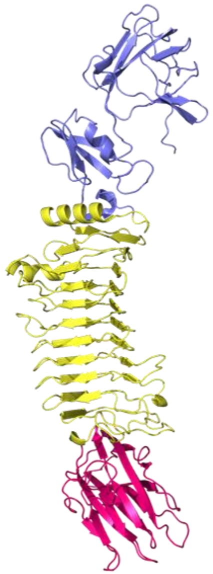 | 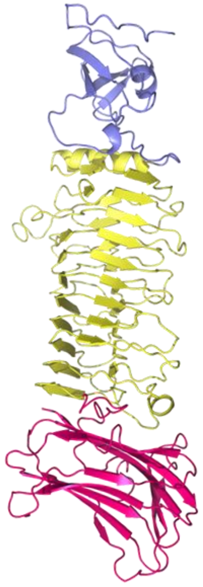 | 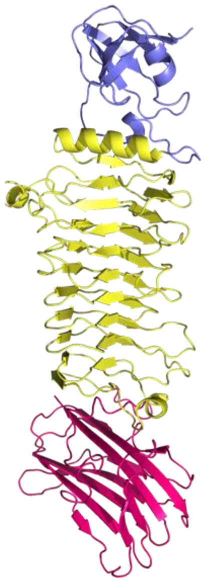 | 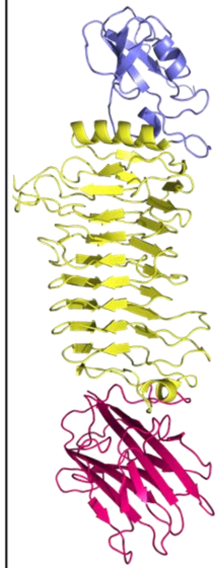 | 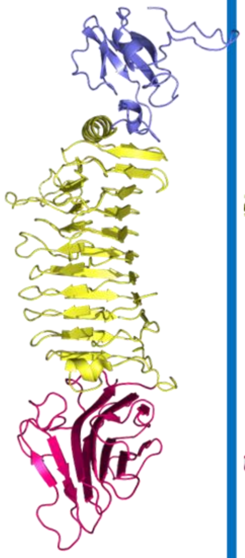 | 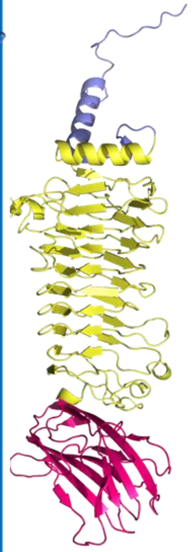 |

*protein was not tested on K13

K2 group 2
TM-score
Central + C-term

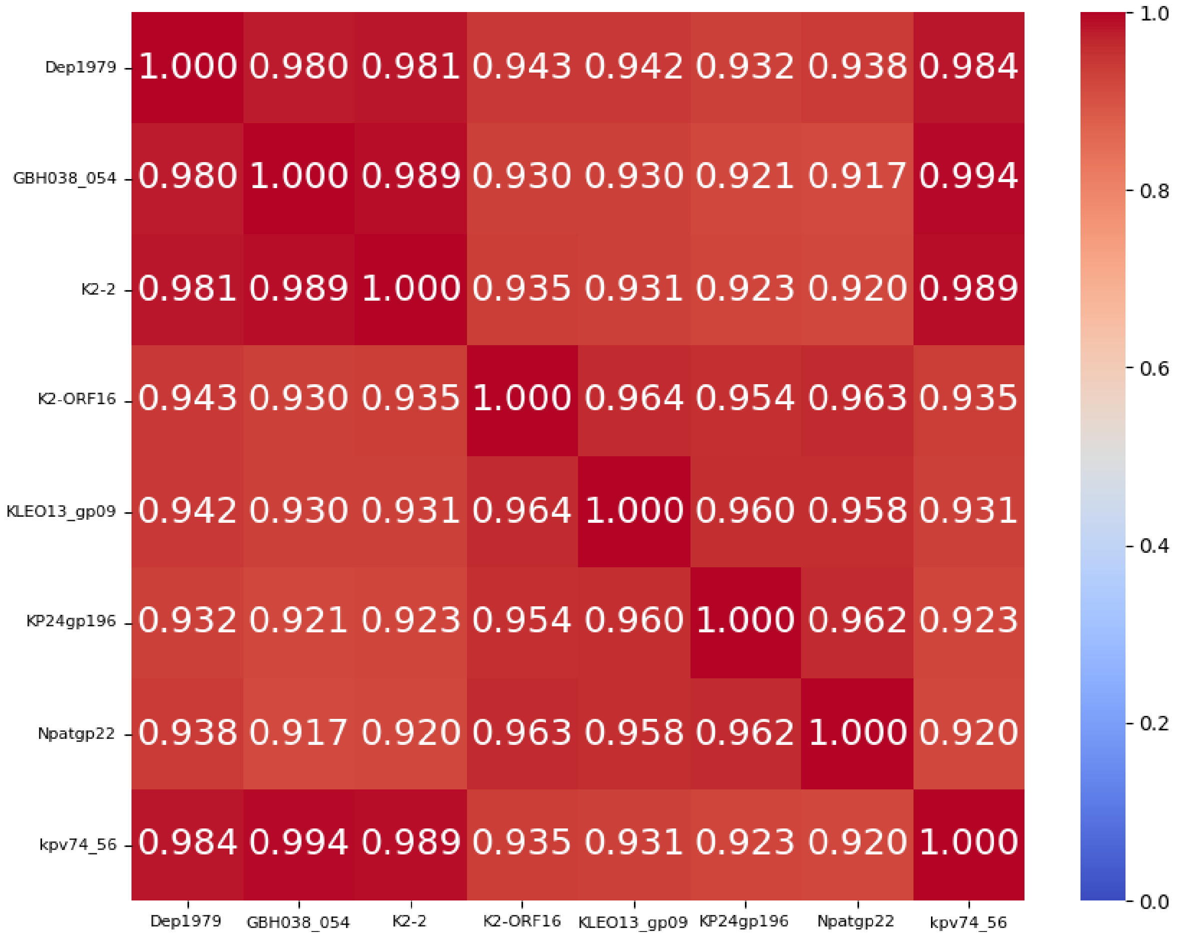


K2 group 2
%identity; cover ≥ 93%
Central + C-term

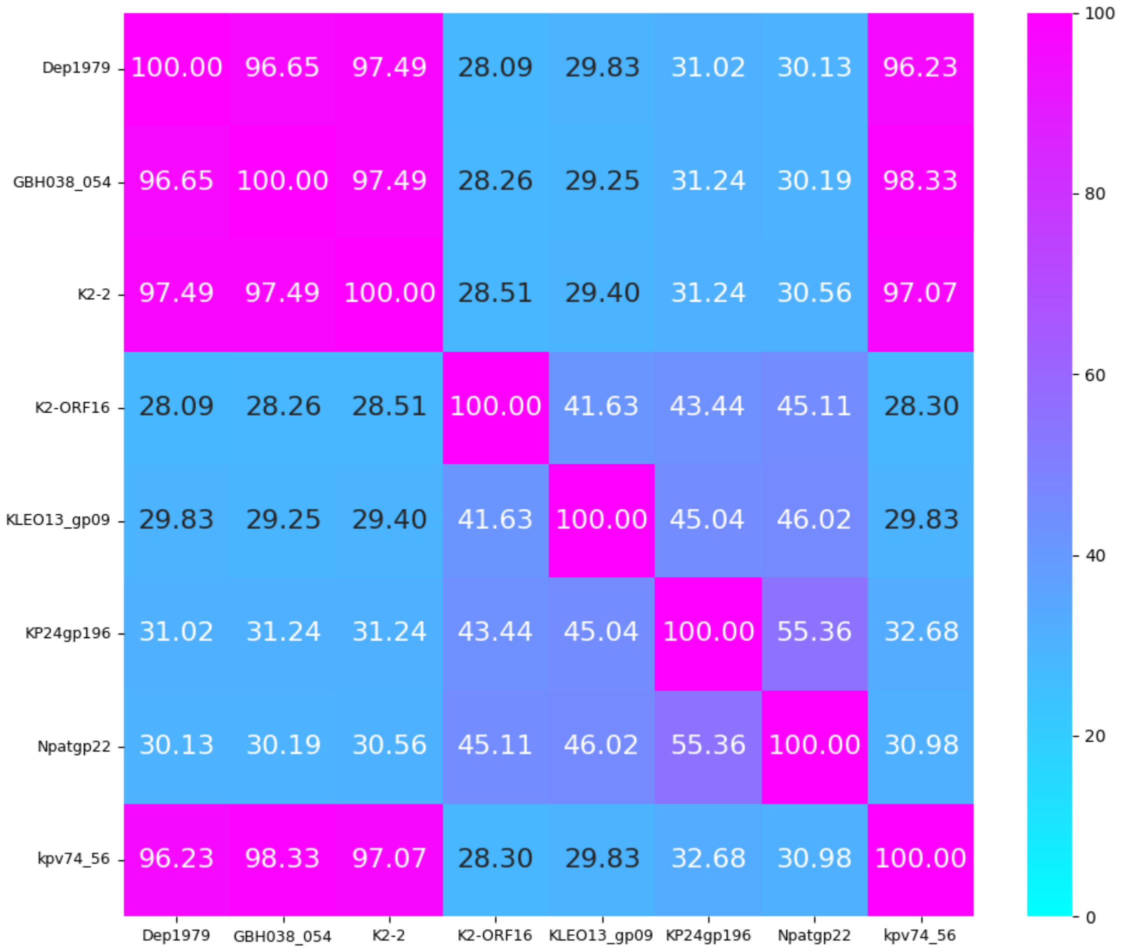

### K2 group 1 + group 2
### TM-score
### Central

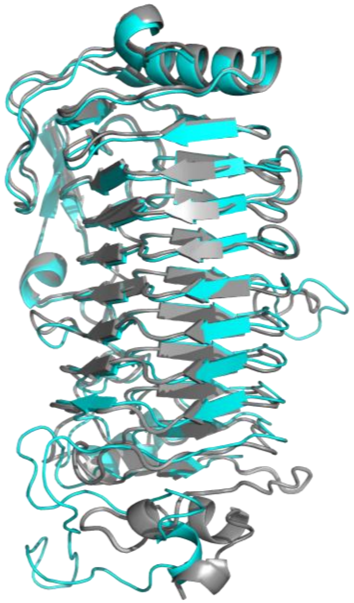

**Structural comparison of the central β-helix domain within the K2 group 1 depolymerases**. The cyan structure represents DpK2, while the grey structures correspond to B1dep, BMacgp22, gp81, and Depo32

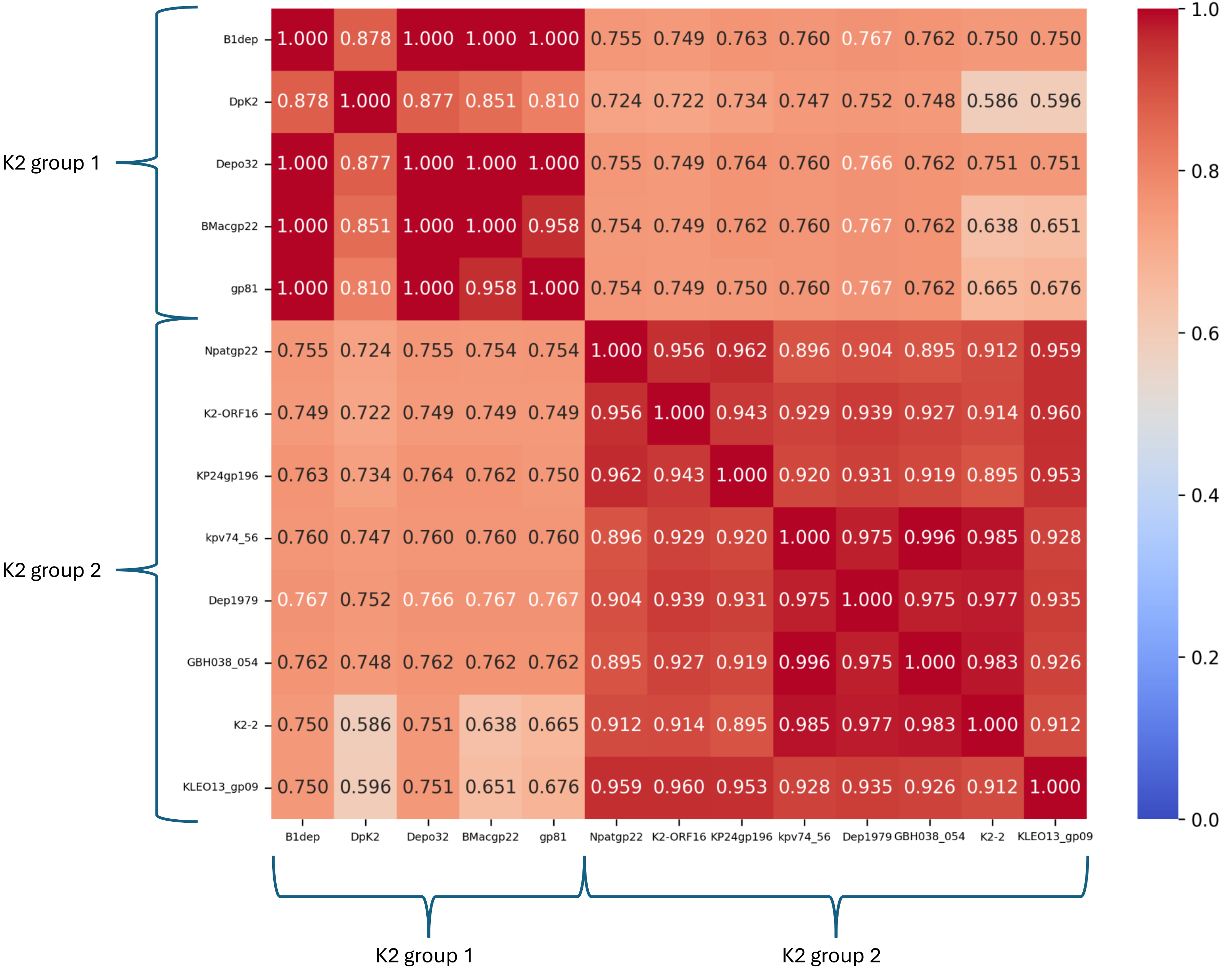

# K3, K5,K7, K8

| K-type target | K3 | K3/KL146 | K5 | | K7 | K8 |
|---|---|---|---|---|---|---|
| Group | Group 1 | Group 2 | Group 1 | | Group 1 | Group 1 |
| Protein name | KP32gp37 | 0391_03 | dep1011 | K5-4 ORF38 | K7dep | K5-4 ORF37 |
| Lenght | 869 aa | 736 aa | 857 aa | 684 aa | 1180 aa | 749 aa |
| Number of proteins in the group | 1 protein | 1 protein | 2 protein | | 1 protein | 1 protein |
| Predicted protein structure | 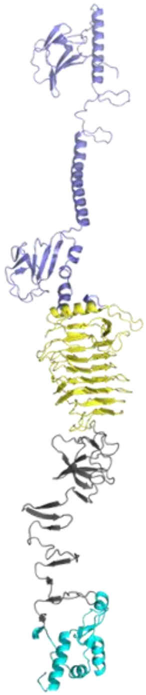 | 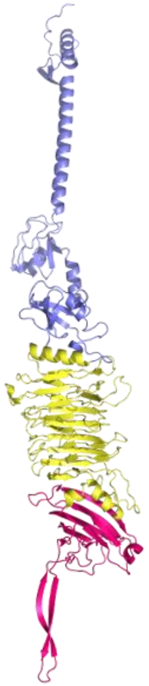 | 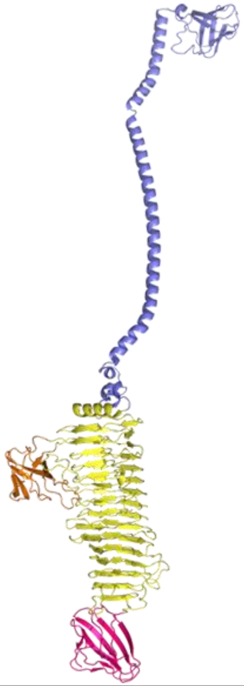 | 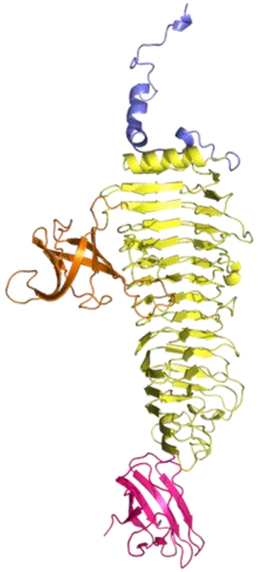 | 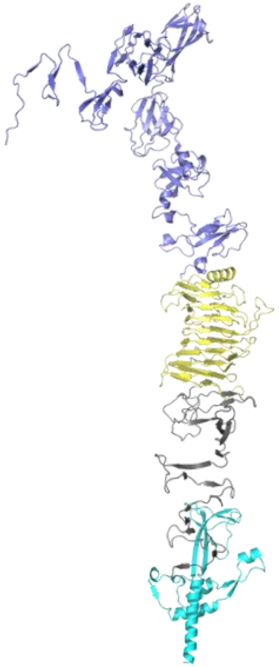 | 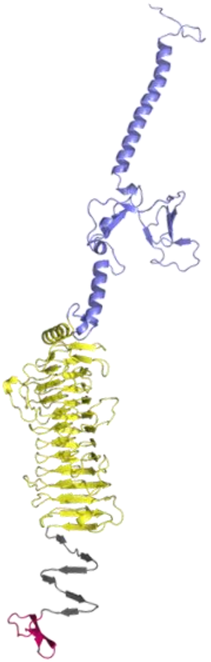 |

K3 group 1 + group 2
TM-score
Central
**KP32gp37 with tail fiber domain (grey part)**

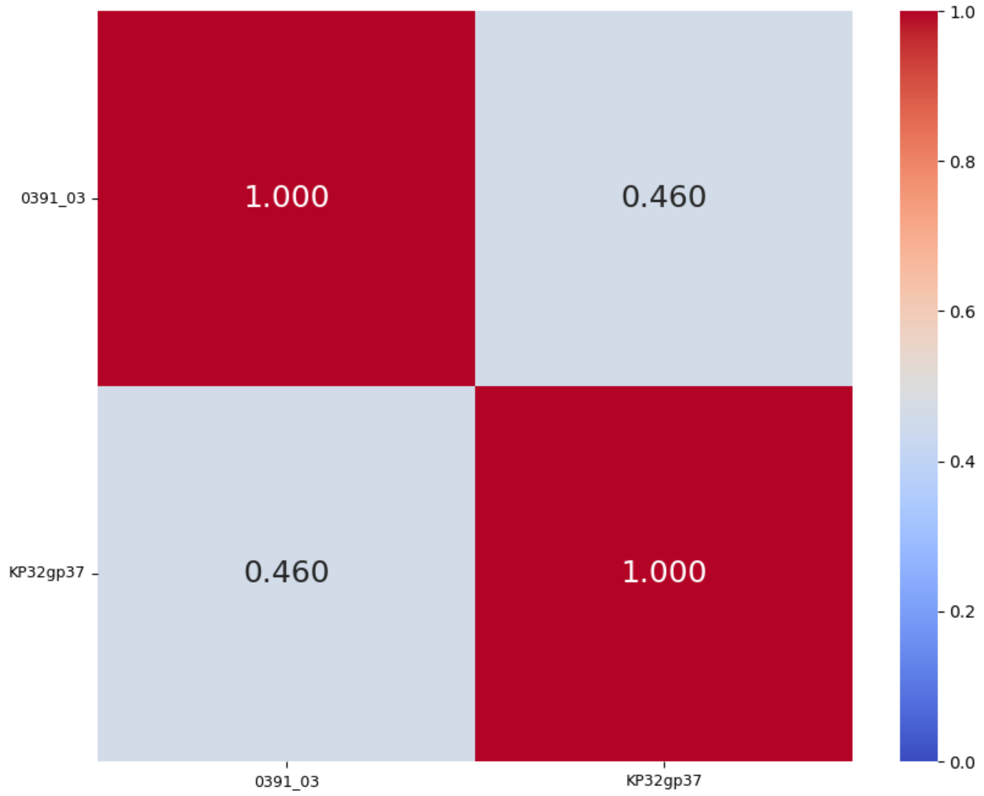


K3 group 1 + group 2
TM-score
Central
**KP32gp37 without tail fiber domain (grey part)**

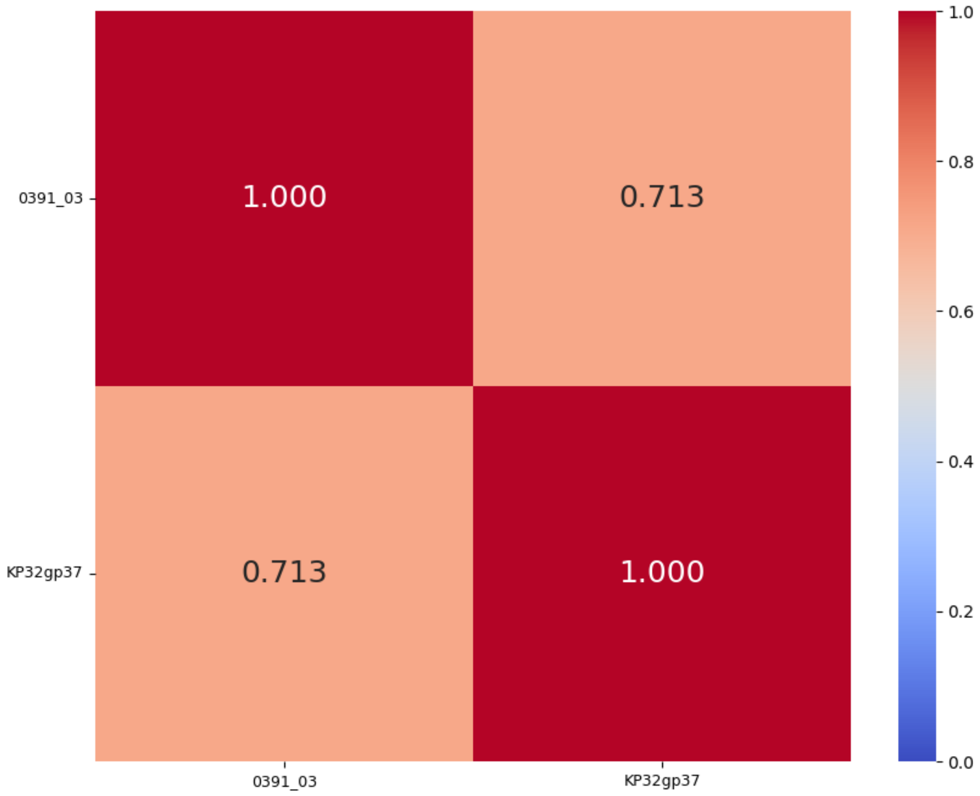

K5 group 1
TM-score
Central + C-term

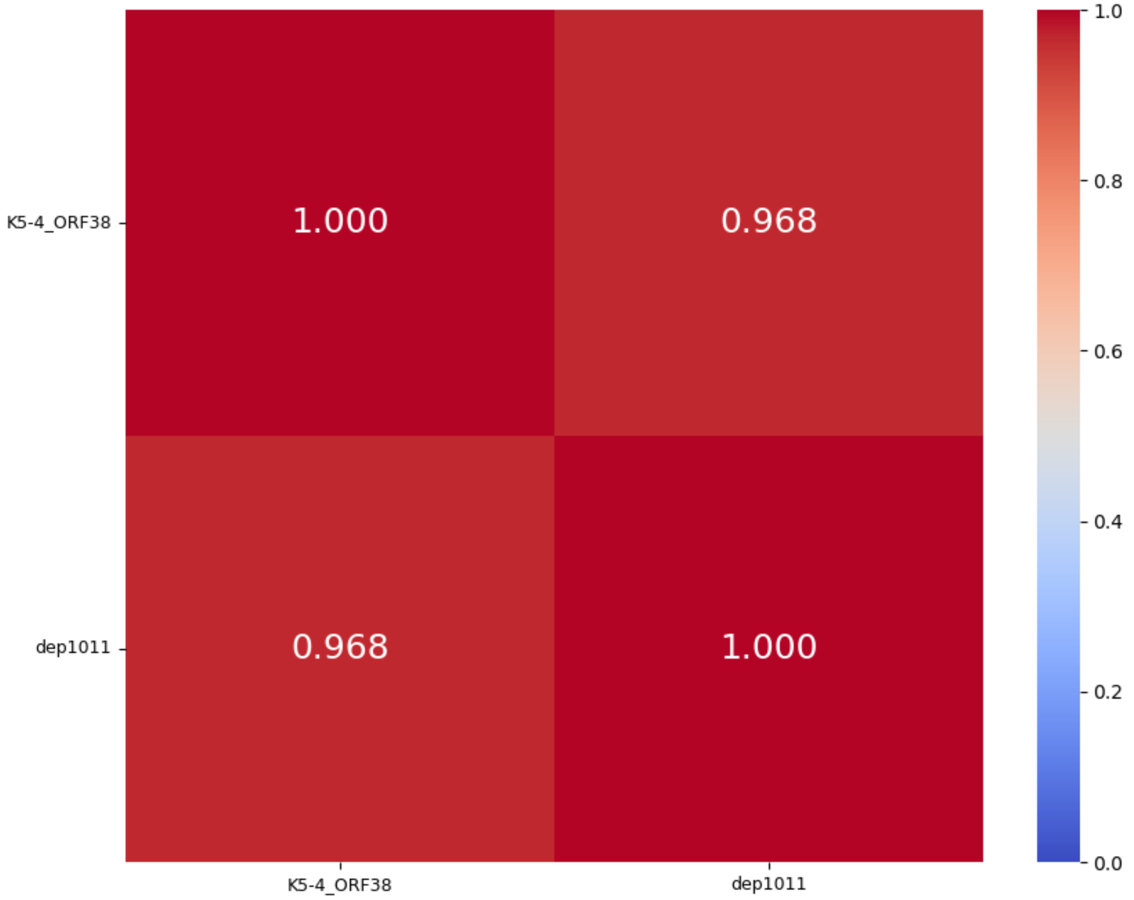


K5 group 1
%identity; cover ≥ 96%
Central + C-term

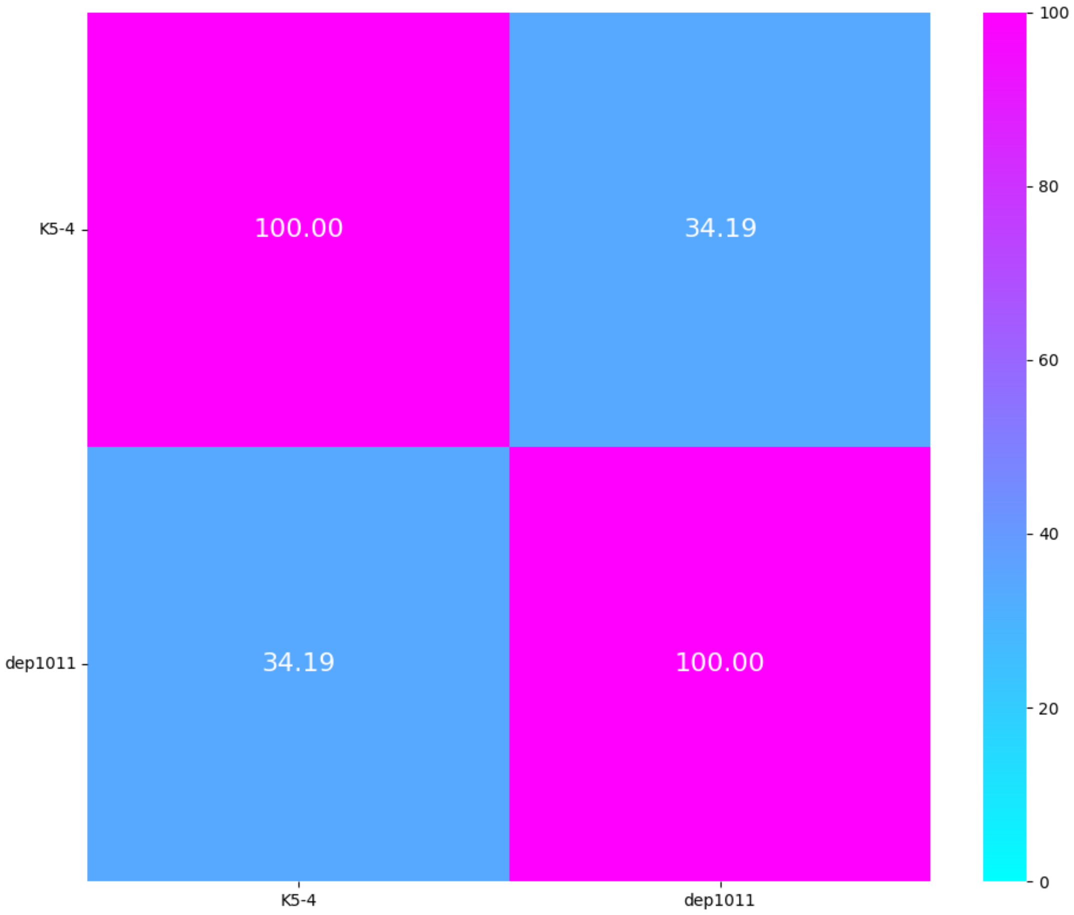

# K9, K11, K14, K19, K20

| K-type target | K9 | K11 | | K14 | K19 | K20 | |
|---|---|---|---|---|---|---|---|
| **Group** | Group 1 | Group 1 | | Group 1 | Group 1 | Group 1 | Group 2 |
| **Protein name** | KANgp229 | K11gp17 | S1-1 | 0574_17 | KP24gp304 | Kl-dep | K20dep |
| **Lenght** | 578 aa | 875 aa | 702 aa | 873 aa | 755 aa | 790 aa | 723 aa |
| **Number of proteins in the group** | 1 protein | 2 proteins | | 1 protein | 1 protein | 1 protein | 1 protein |
| **Predicted protein structure** | 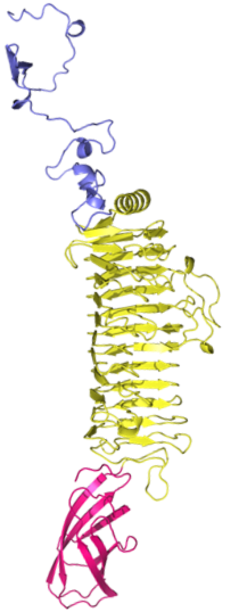 | 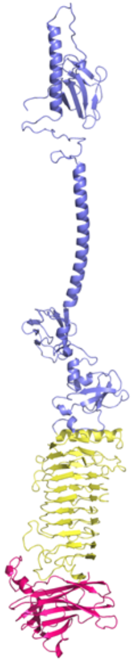 | 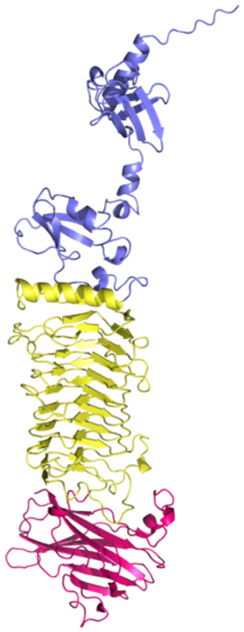 | 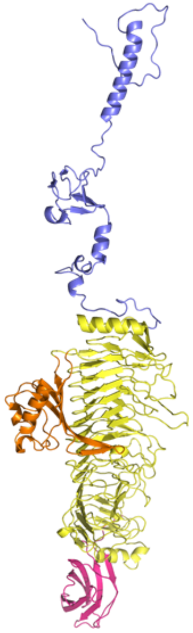 | 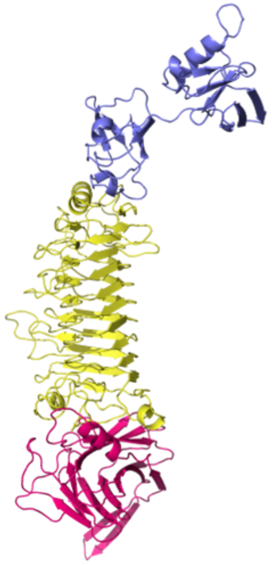 | 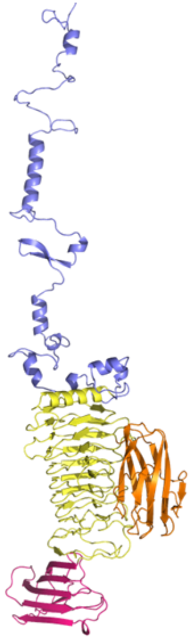 | 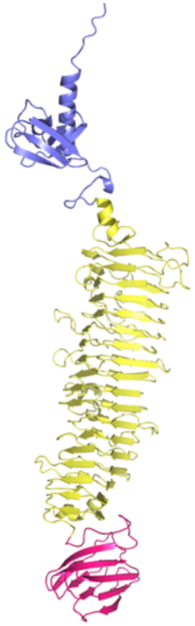 |

K11 group 1
TM-score
Central + C-term

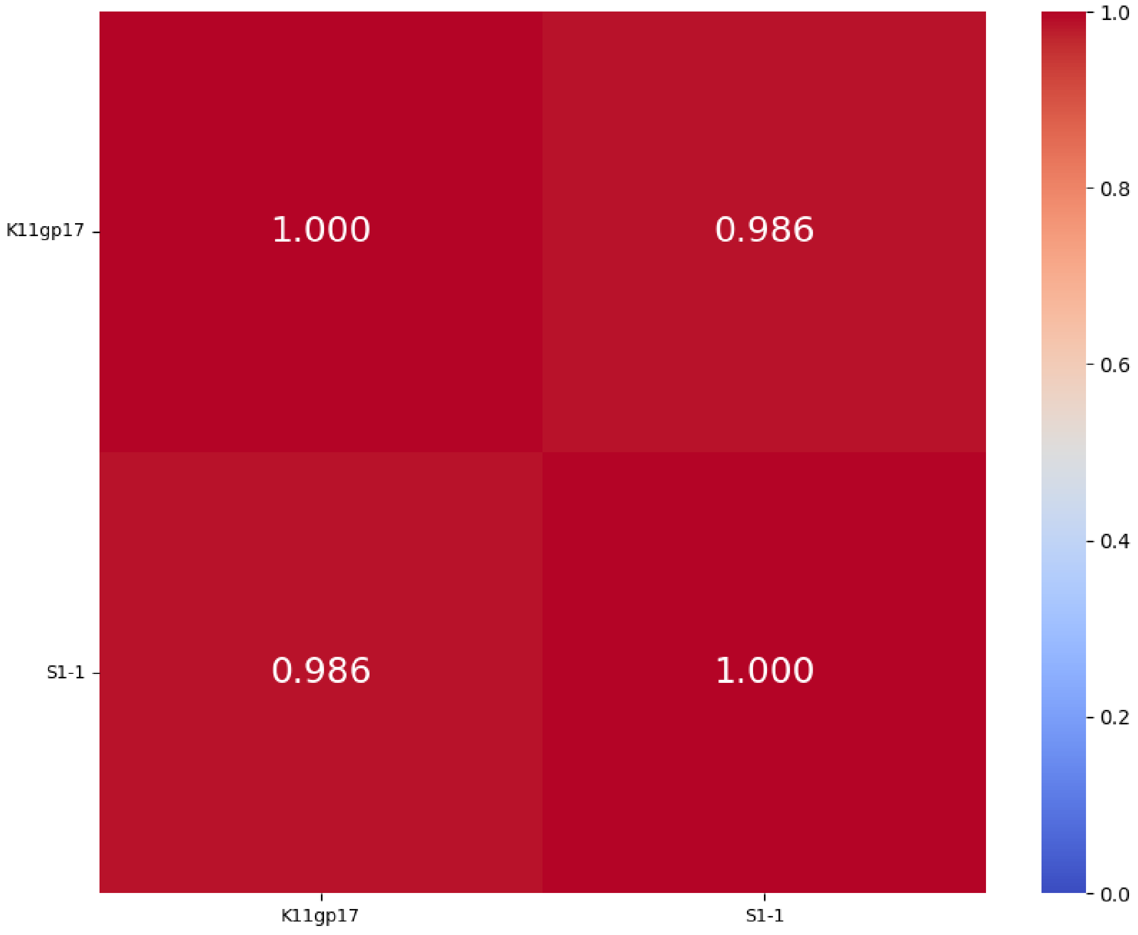


K11 group 1
%identity; cover ≥ 99%
Central + C-term

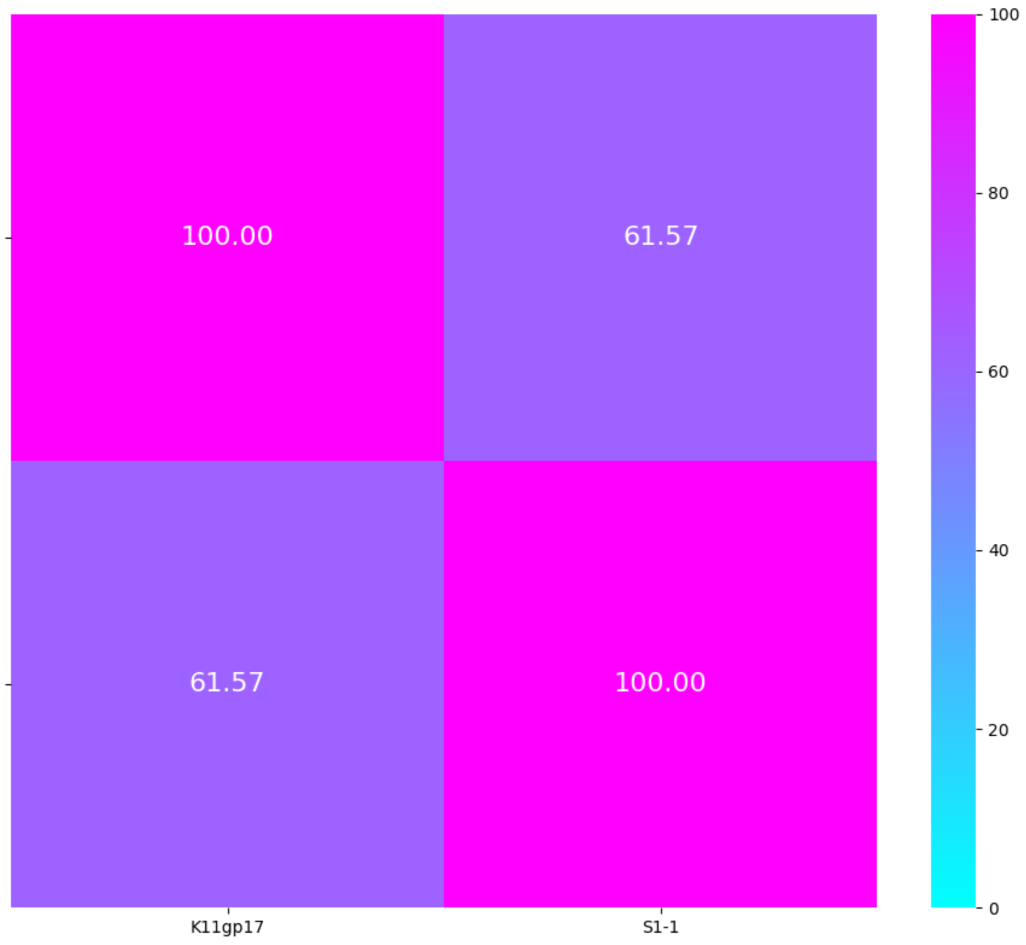

# K21

| K-type target | K21* | K21/KL163 | KL21* |
|---|---|---|---|
| Group | Group 1 | | |
| Protein name | S1-3 | KP32gp38 | RBP2 |
| Lenght | 651 aa | 576 aa | 575 aa |
| Number of proteins in the group | 3 proteins | | |
| Predicted protein structure | 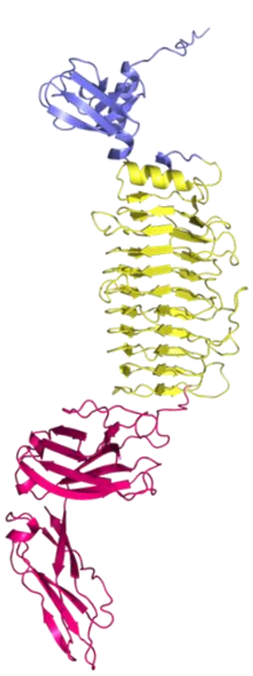 | 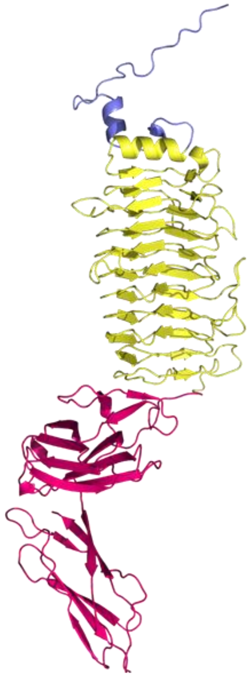 | 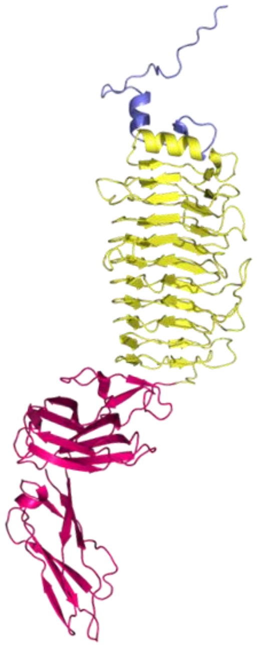 |

*protein was not tested on KL163

#### Cleavage site RBP2

(Lukianova et al. 2023)

**K21**
CPS repeating unit

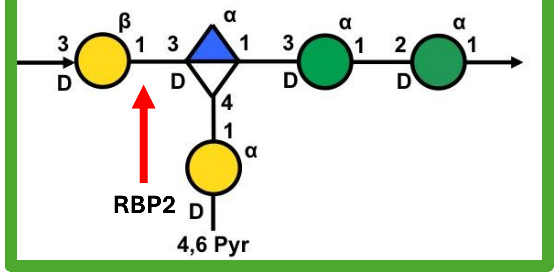


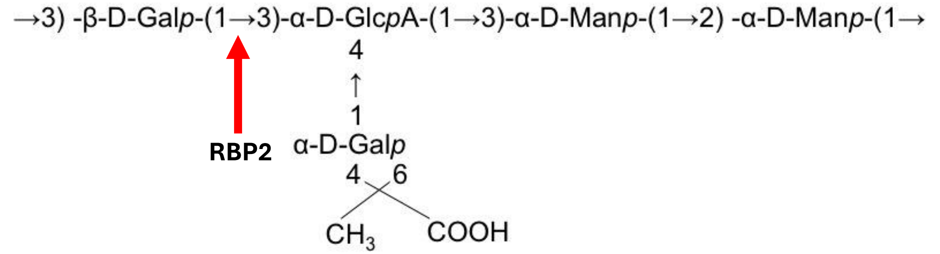


Active Site Residues **KP32gp38**

(Squeglia et al. 2020)

**Glu239, Asp229, Asp241, Glu170**

K21 group 1
TM-score
Centra + C-term

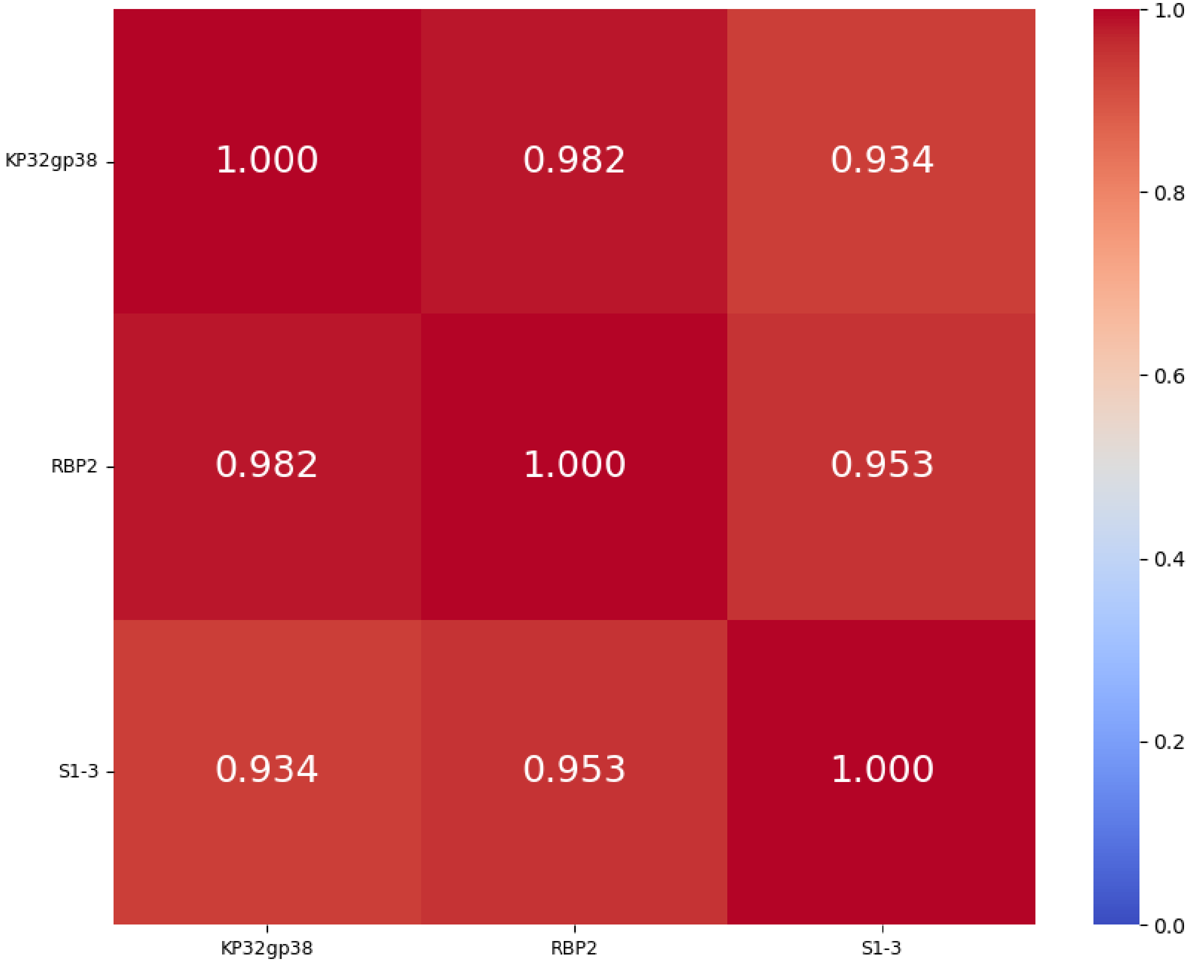


K21 group 1
%identity; cover ≥ 98%
Centra + C-term

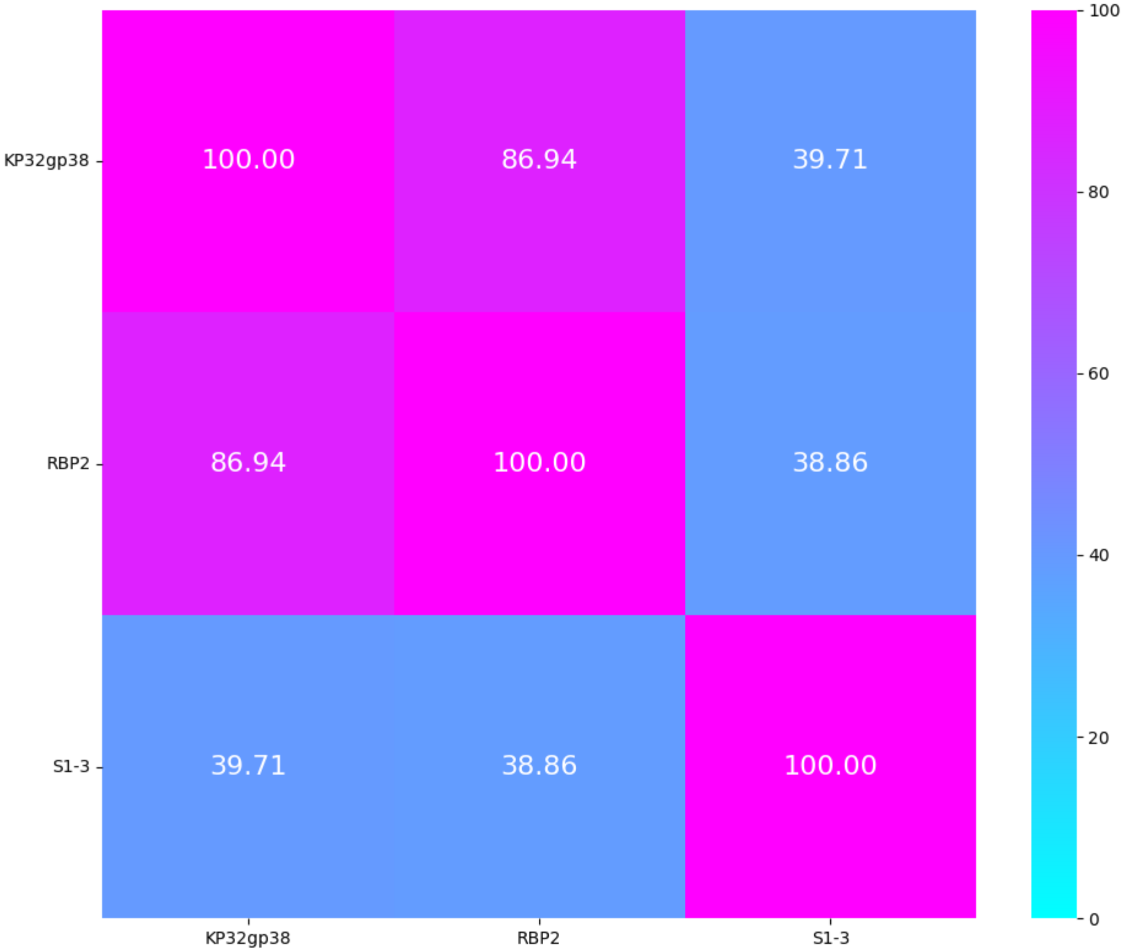

# **K22**/K37/KL111, **K23**

| **K-type target** | **K22/K37/K111** | **K23** | | | | |
|---|---|---|---|---|---|---|
| **Group** | Group 1 | Group 1 | | | | |
| **Protein name** | KLEO13gp10 | 1409_59 | 1441_47 | 1248_57 | DepS8 | Dep622 |
| **Lenght** | 777 aa | 852 aa | 704 aa | 704 aa | 607 aa | 555 aa |
| **Number of proteins in the group** | 1 protein | 5 proteins | | | | |
| **Predicted protein structure** | 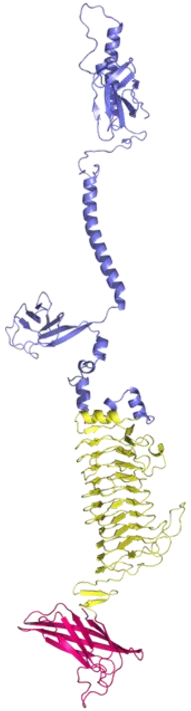 | 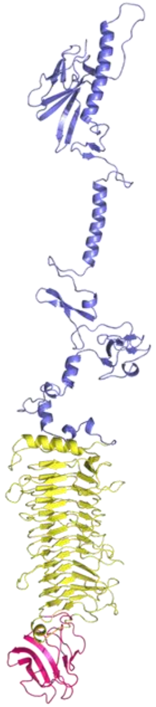 | 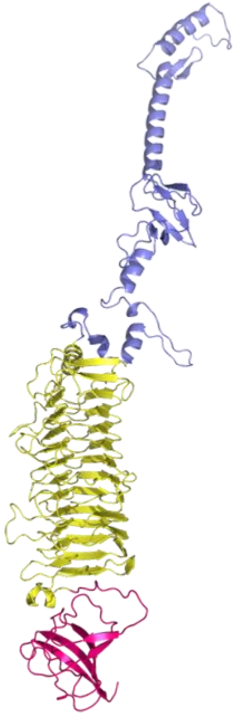 | 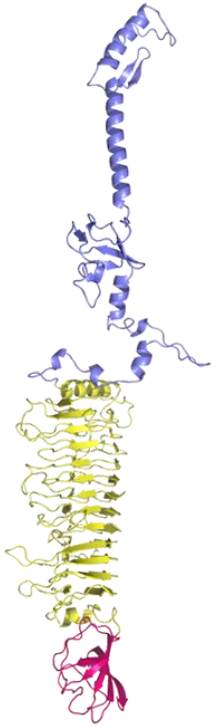 | 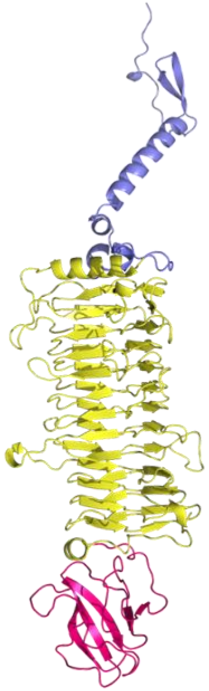 | 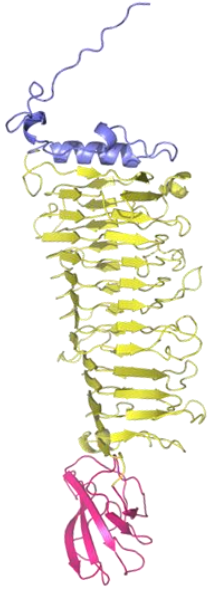 |

K23 group 1
TM-score
Central + C-term

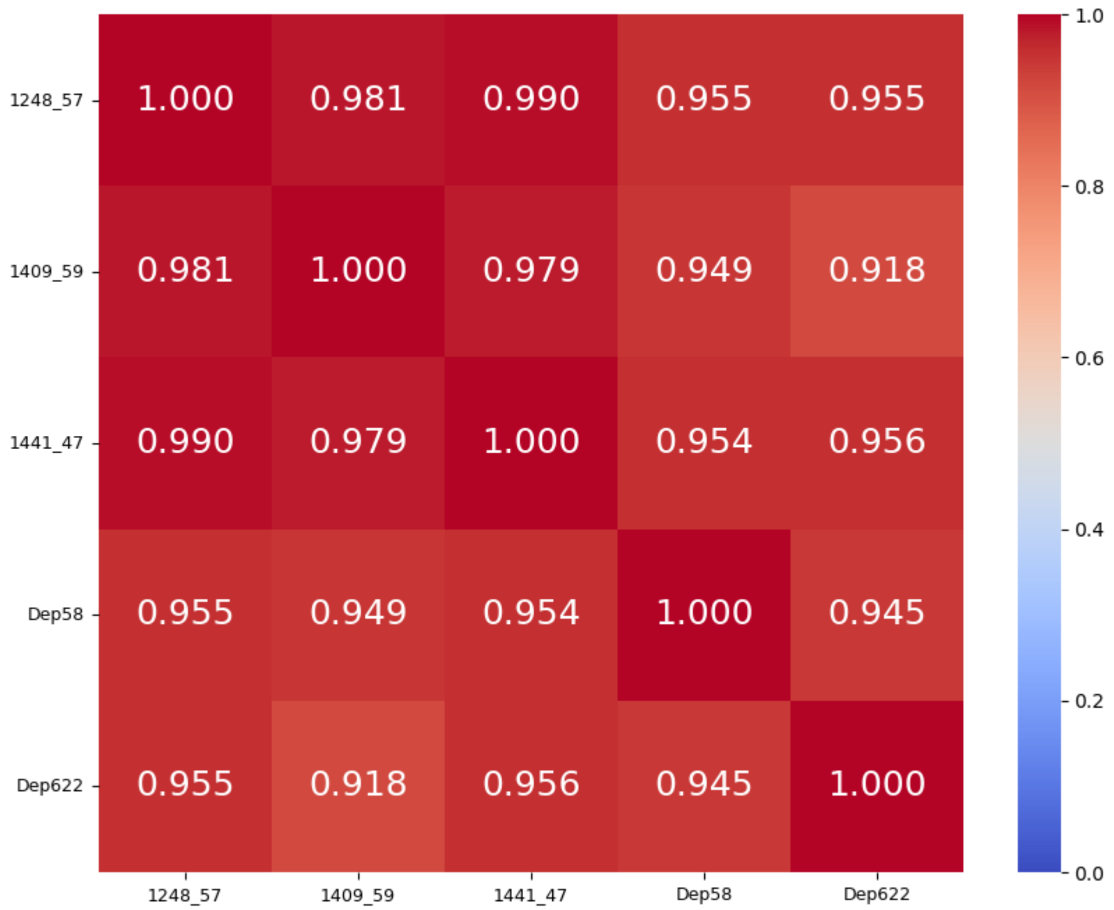


K23 group 1
%identity; cover ≥ 95%
Central + C-term

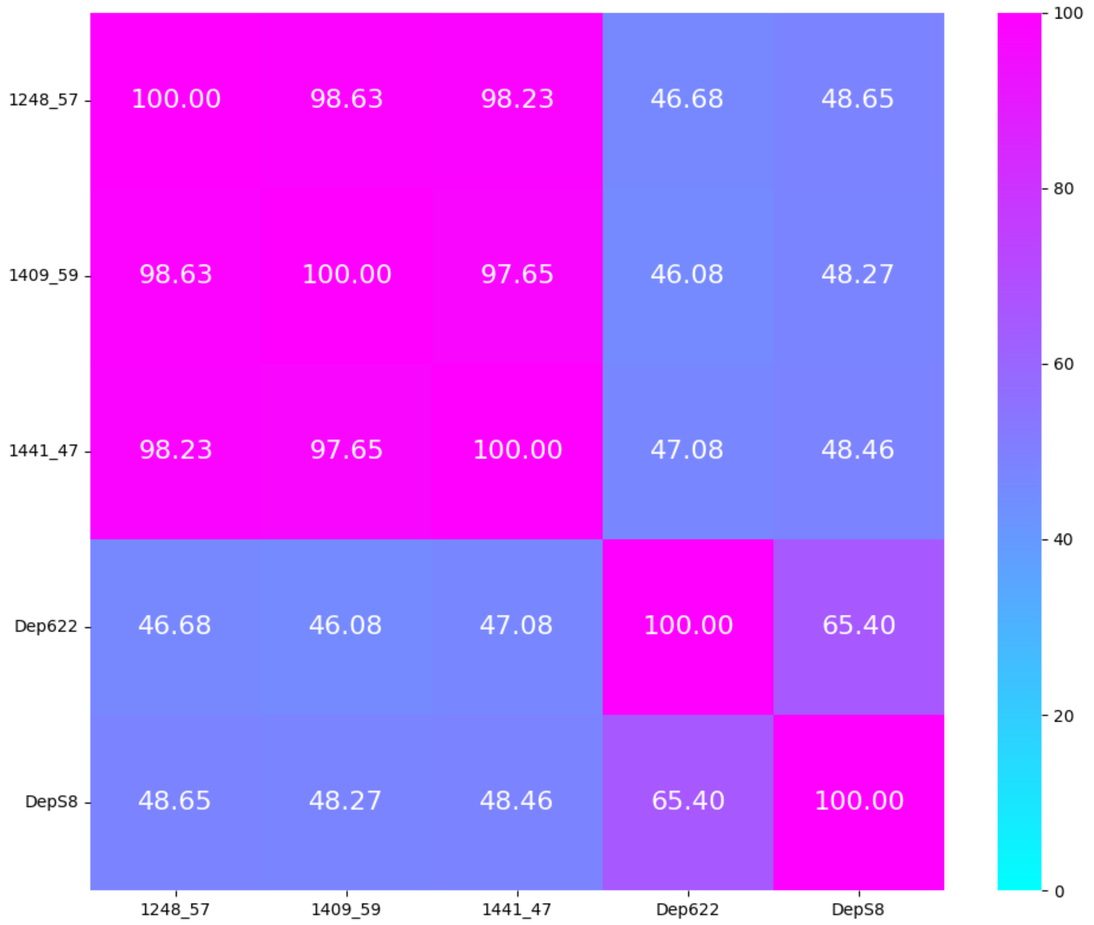

# **K25**/KL119, **K27**, **K28**

| **K-type target** | **K25/KL119** | **K25*** | **K27** | | **K28** | |
|---|---|---|---|---|---|---|
| **Group** | Group 1 | | Group 1 | | Group 1 | |
| **Protein name** | KP24gp300 | S2-2 | K27dep | gp12 | 1251_37 | KANgp225 |
| **Lenght** | 598 aa | 584 aa | 1294 aa | 1242 aa | 677 aa | 605 aa |
| **Number of proteins in the group** | 2 proteins | | 2 proteins | | 2 proteins | |
| **Predicted protein structure** | 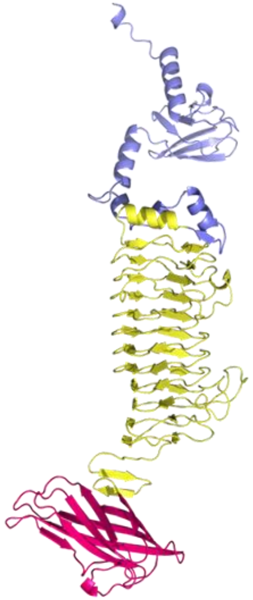 | 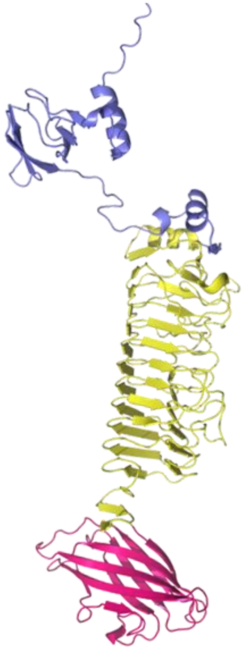 | 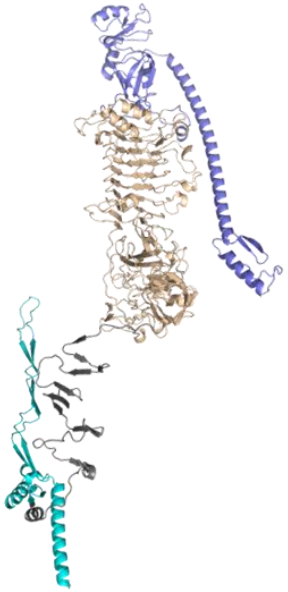 | 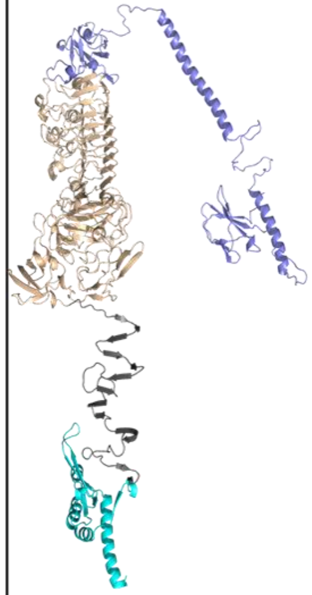 | 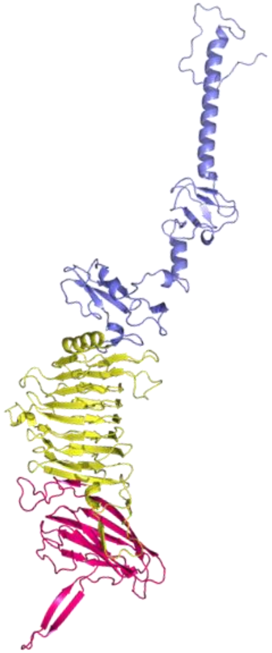 | 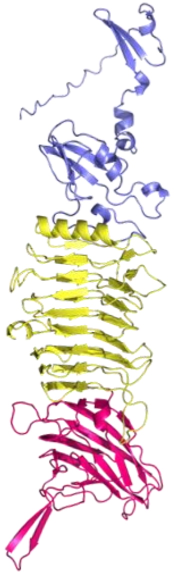 |

*protein was not tested on KL119

K25 group 1
TM-score
Central + C-term

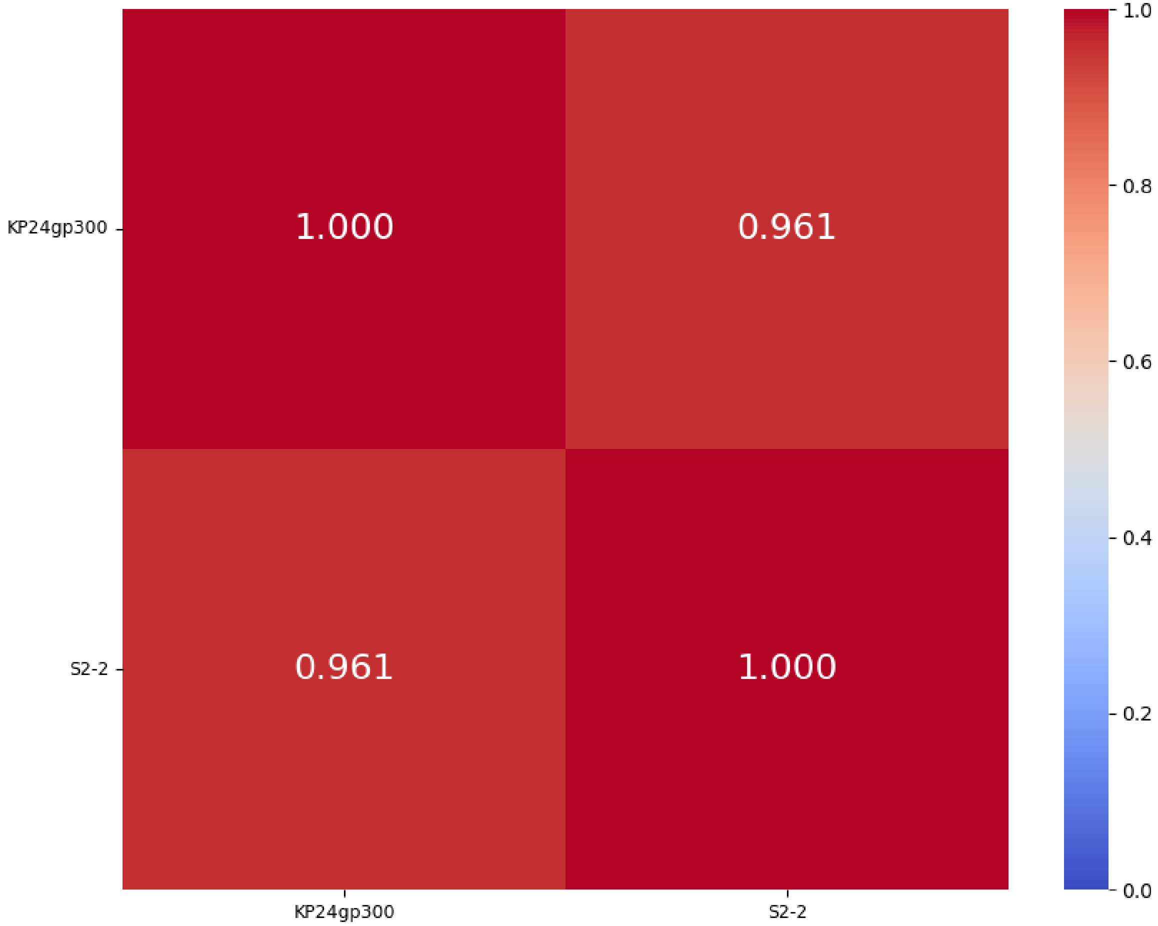


K25 group 1
%identity; cover ≥ 99%
Central + C-term

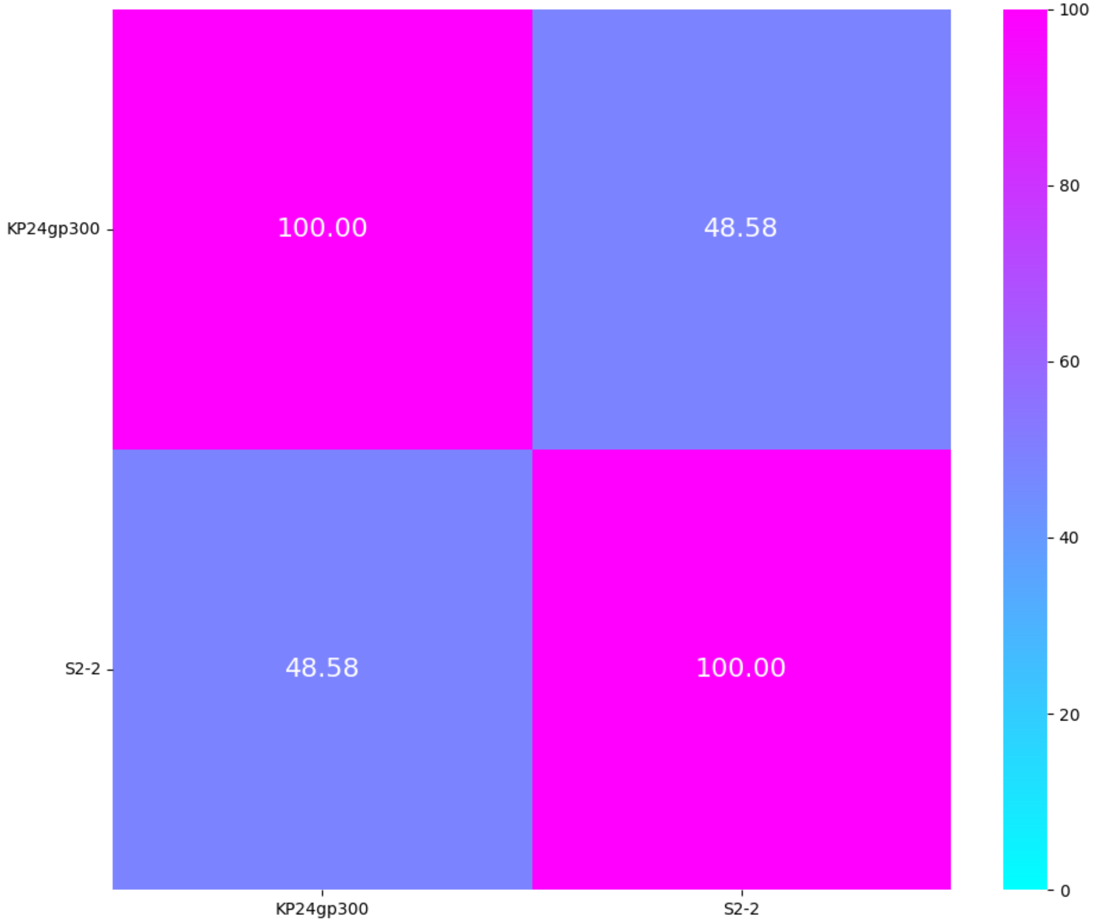

K27 group 1
TM-score
Central + C-term

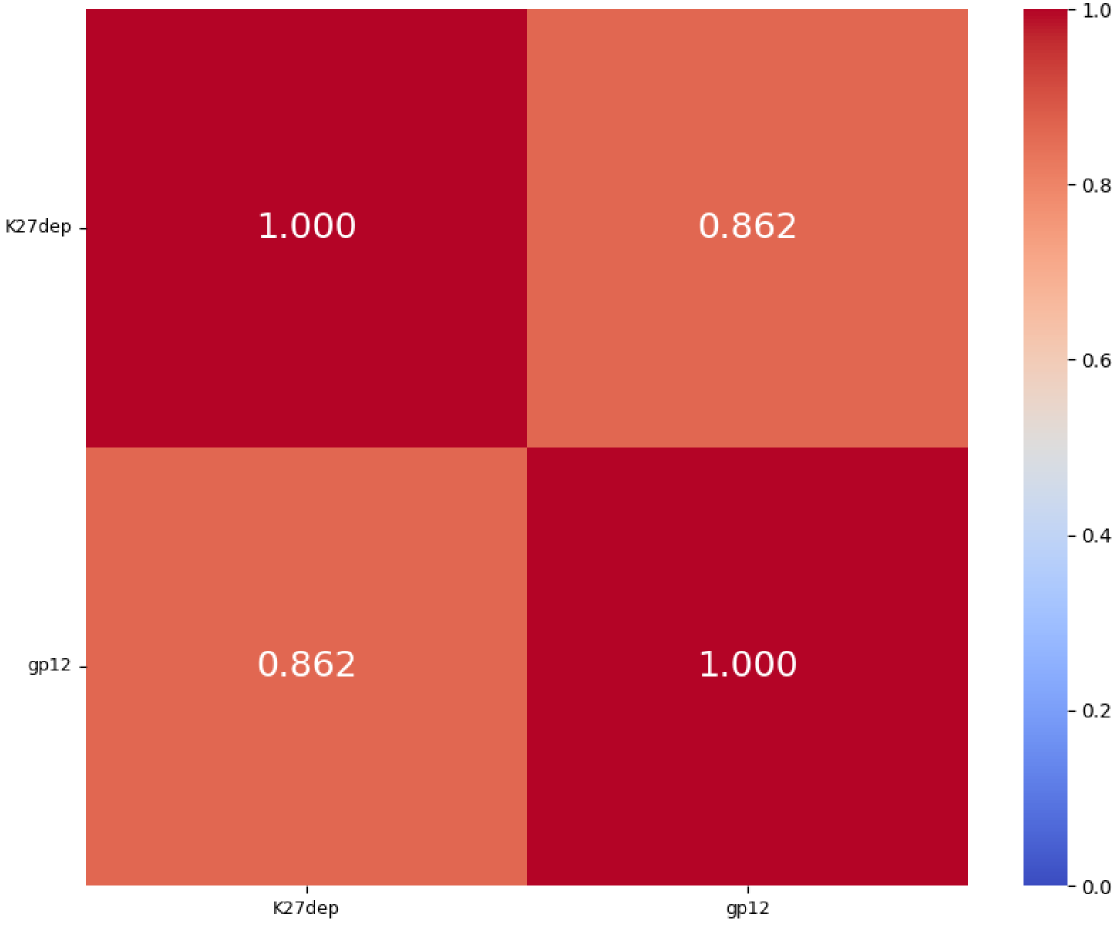


K27 group 1
%identity; cover ≥ 89%
Central + C-term

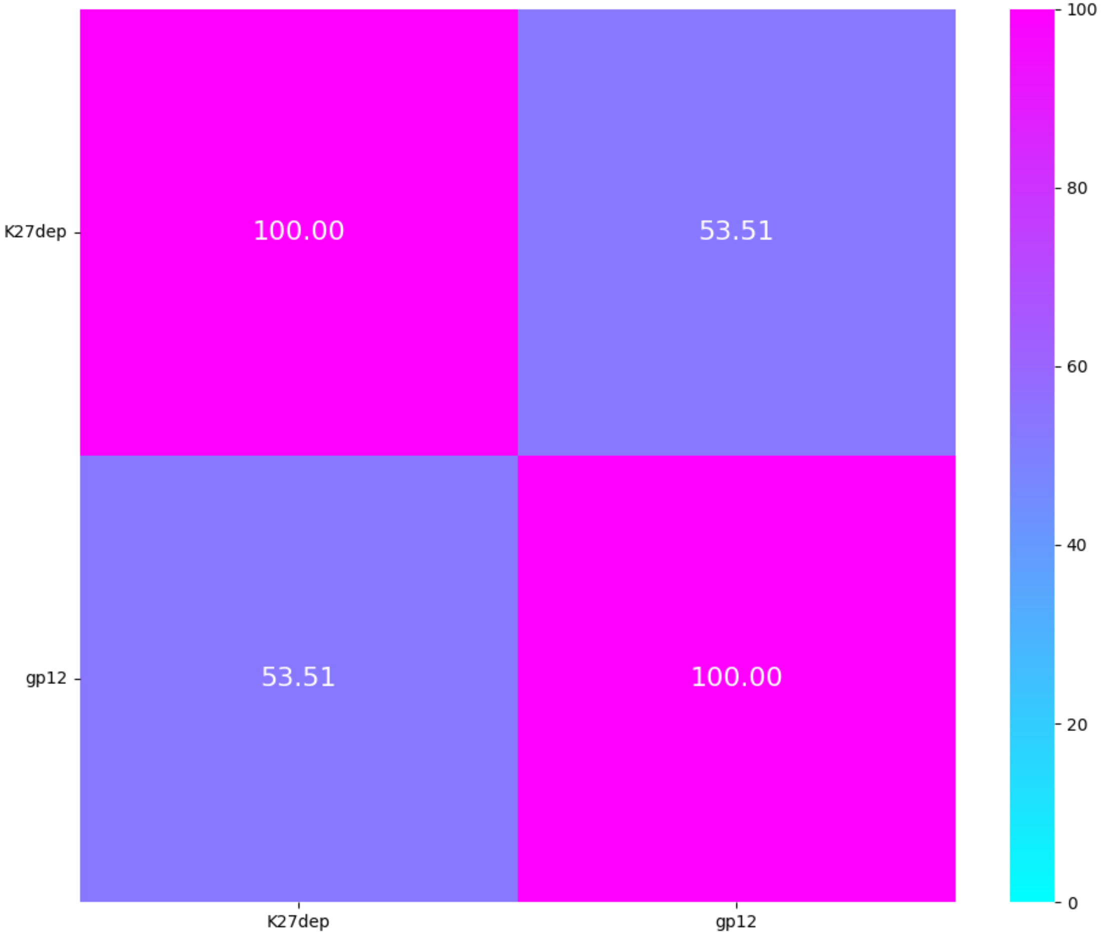

K28 group 1
TM-score
Central + C-term

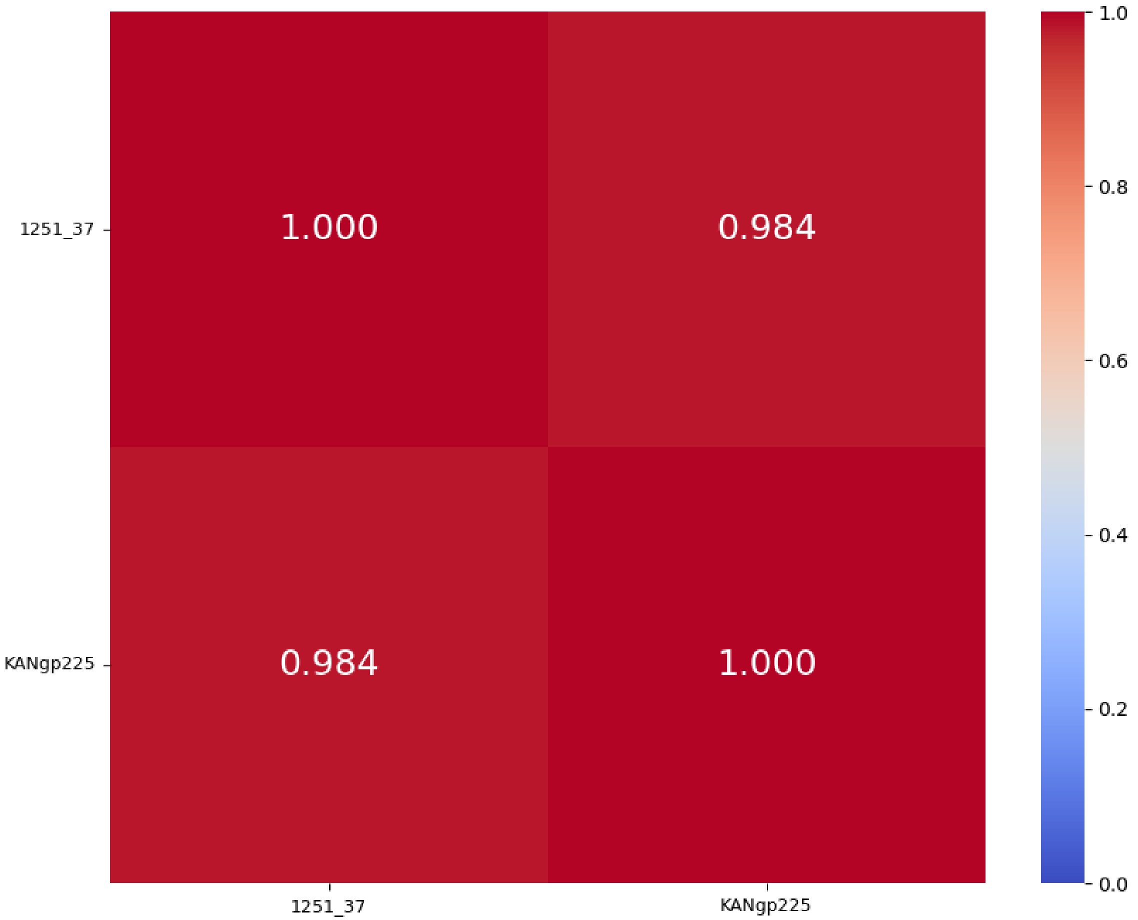


K28 group 1
%identity; cover = 100%
Central + C-term

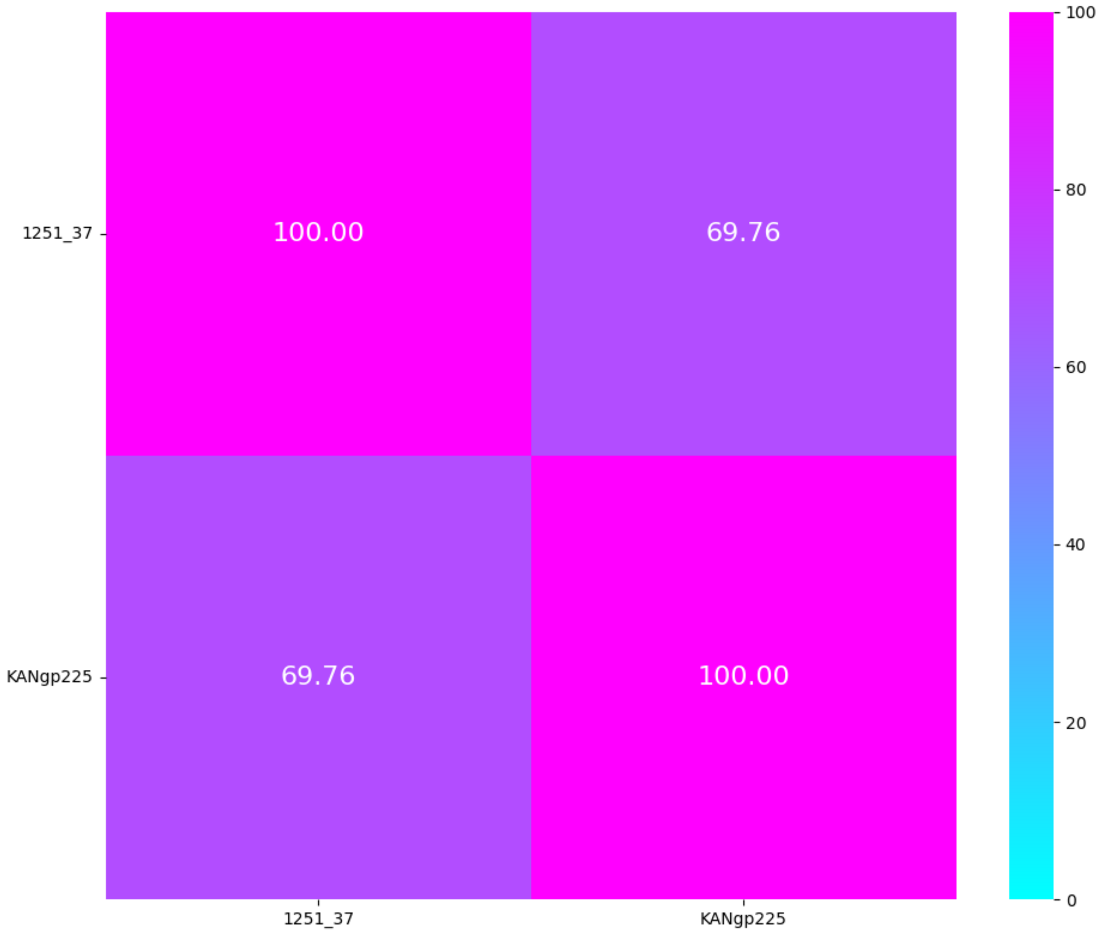

# **K30**/K69, **K31, K32, K35, K38**

| **K-type target** | **K30/69** | | **K31** | **K32** | **K35** | | **K38** |
|---|---|---|---|---|---|---|---|
| **Group** | Group 1 | | Group 1 | Group 1 | Group 1 | | Group 1 |
| **Protein name** | K5-2 ORF37 | S2-6 | K11gp0043 | 914_77 | KP24gp301 | S2-3 | 617_77 |
| **Lenght** | 792 aa | 767 aa | 596 aa | 536 aa | 915 aa | 779 aa | 1258 aa |
| **Number of proteins in the group** | 2 proteins | | 1 protein | 1 protein | 2 proteins | | 1 protein |
| **Predicted protein structure** | 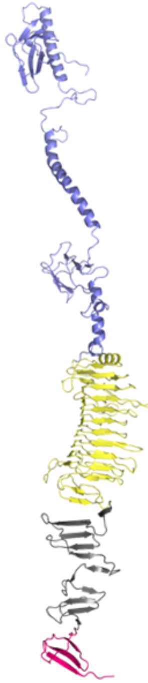 | 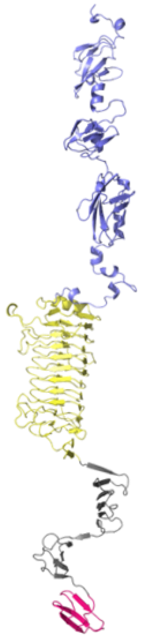 | 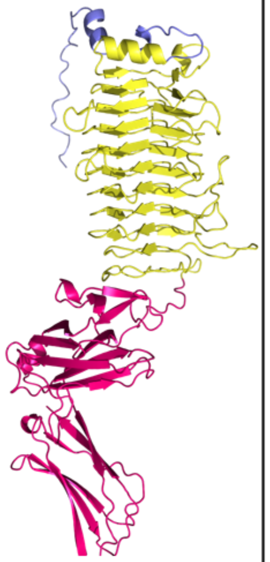 | 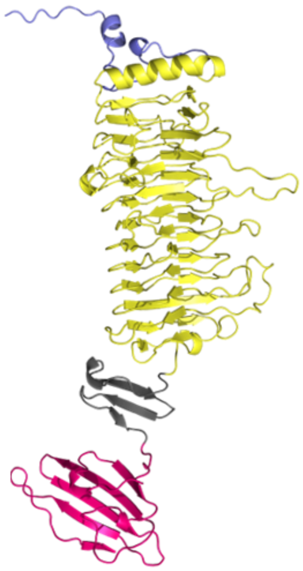 | 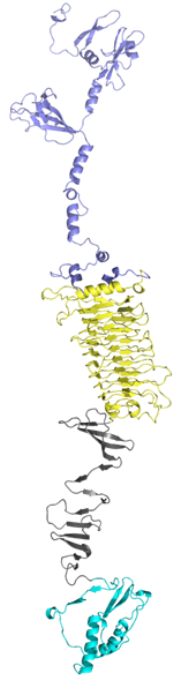 | 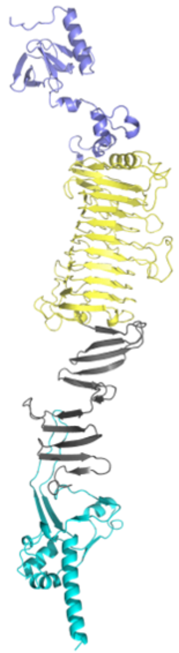 | 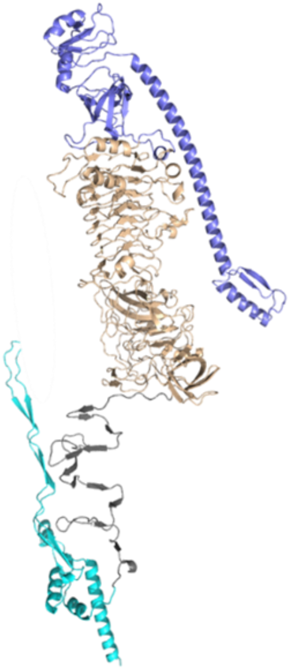 |

K30_69 group 1
TM-score
Central + C-term

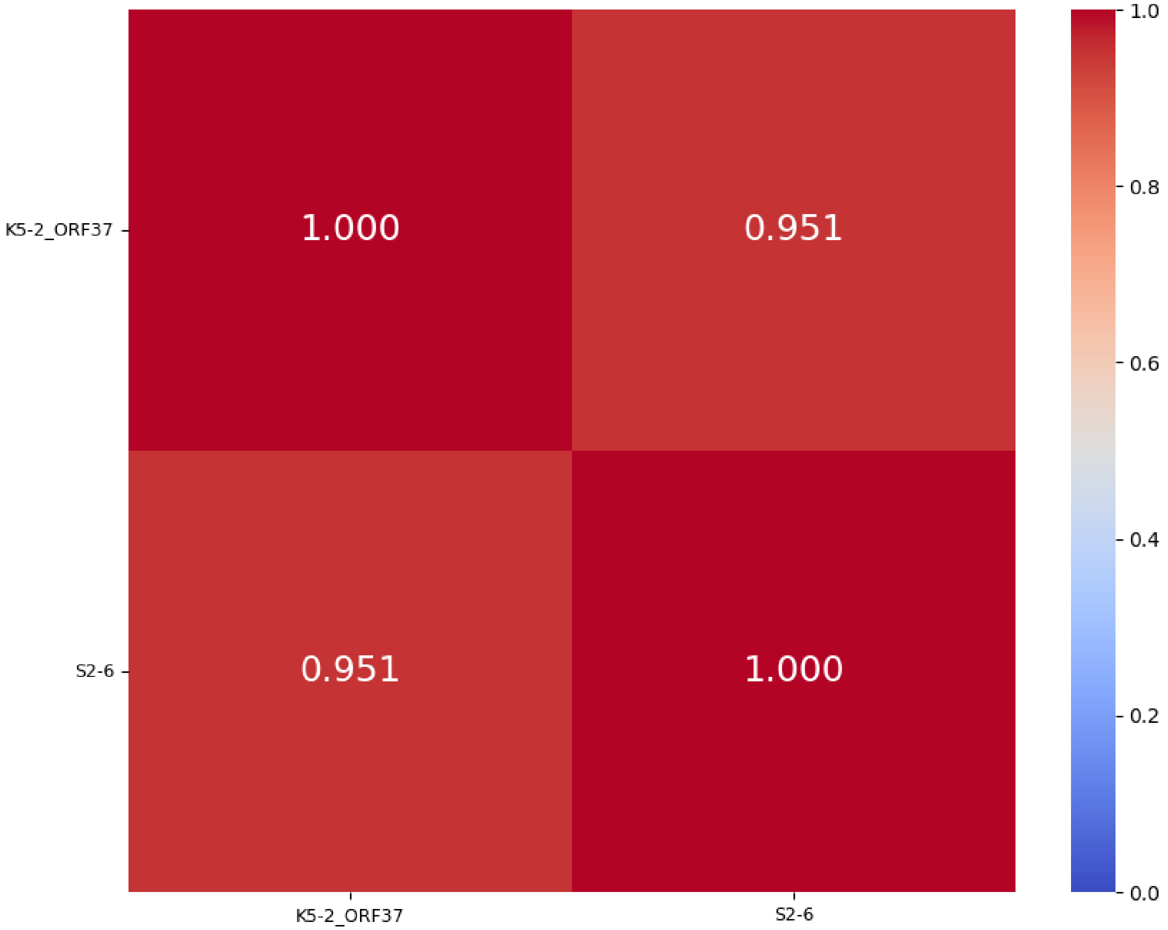


K30_69 group 1
%identity; cover = 100%
Central + C-term

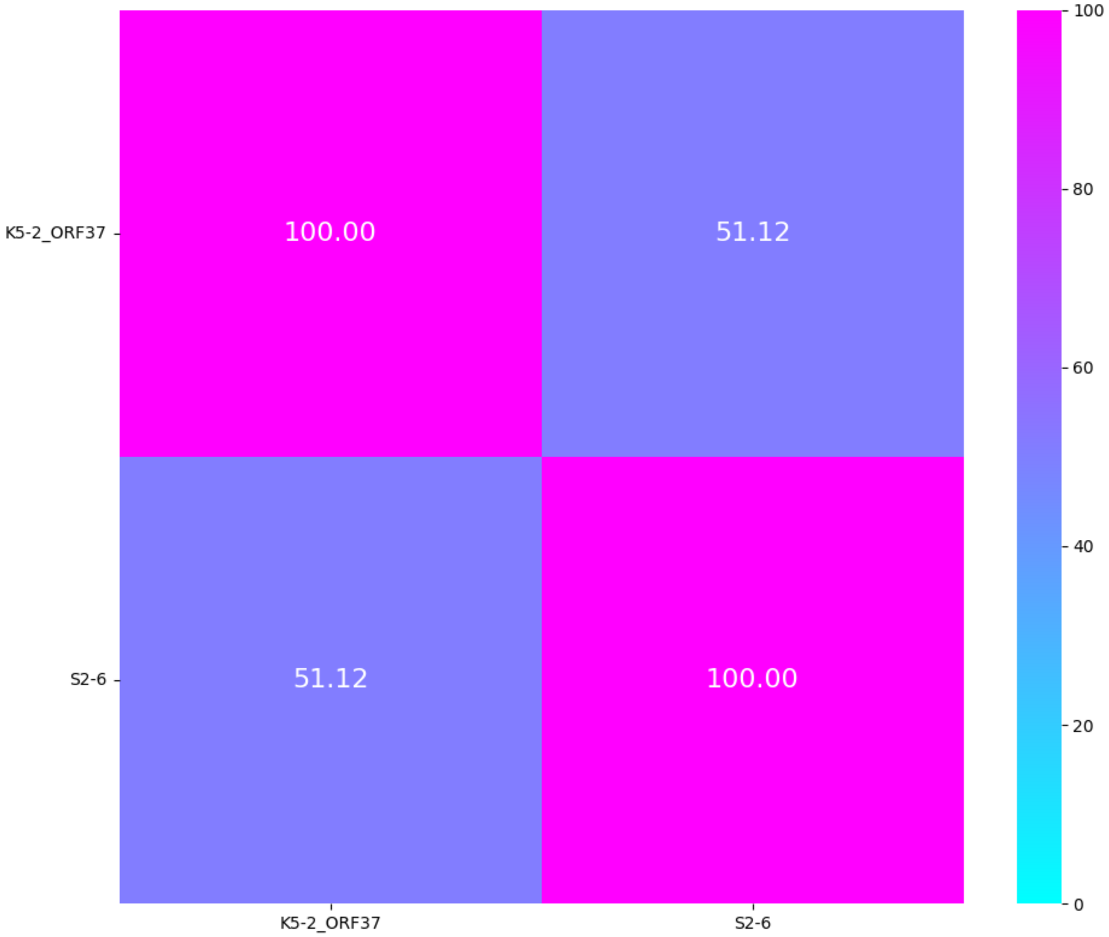

K35 group 1
TM-score
Central + C-term

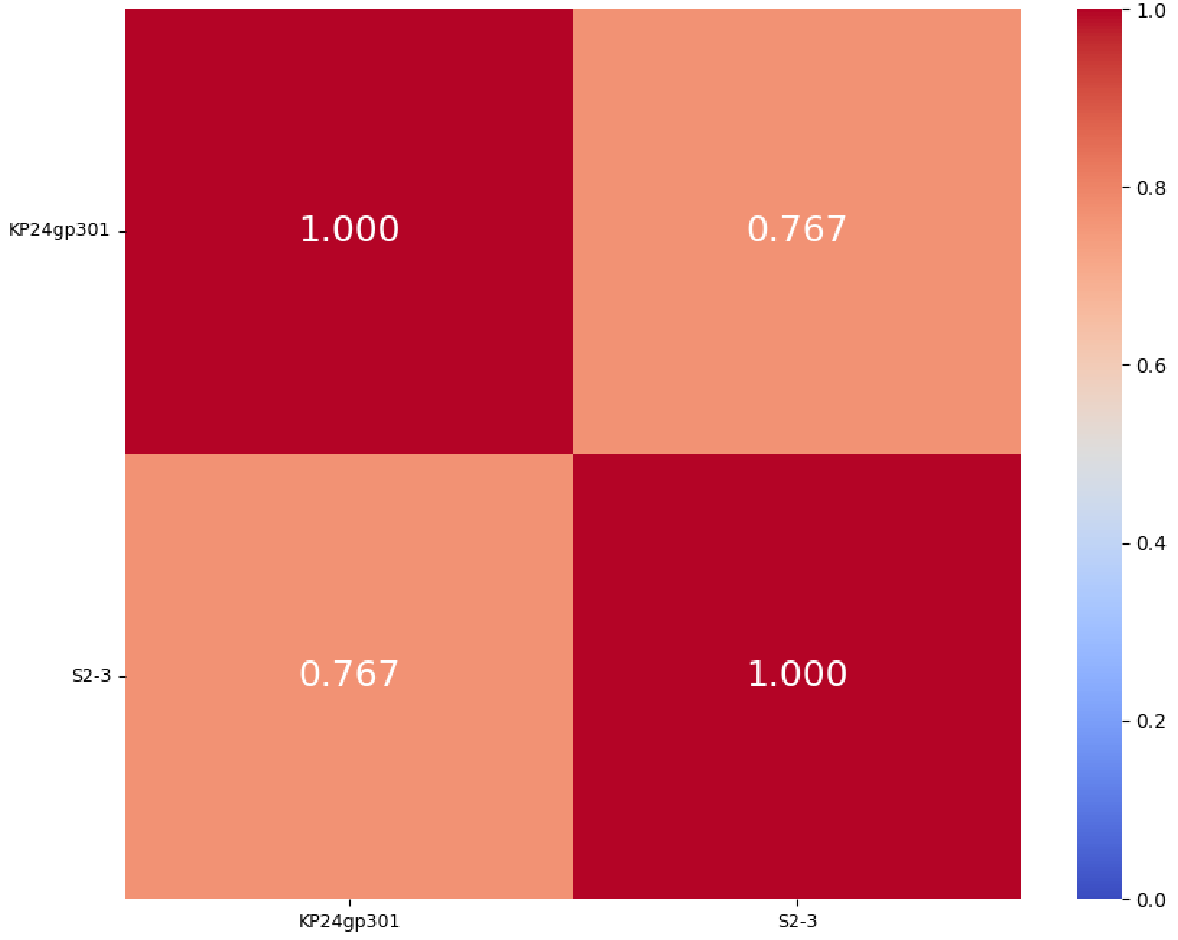


K35 group 1
%identity; cover ≥ 94%
Central + C-term

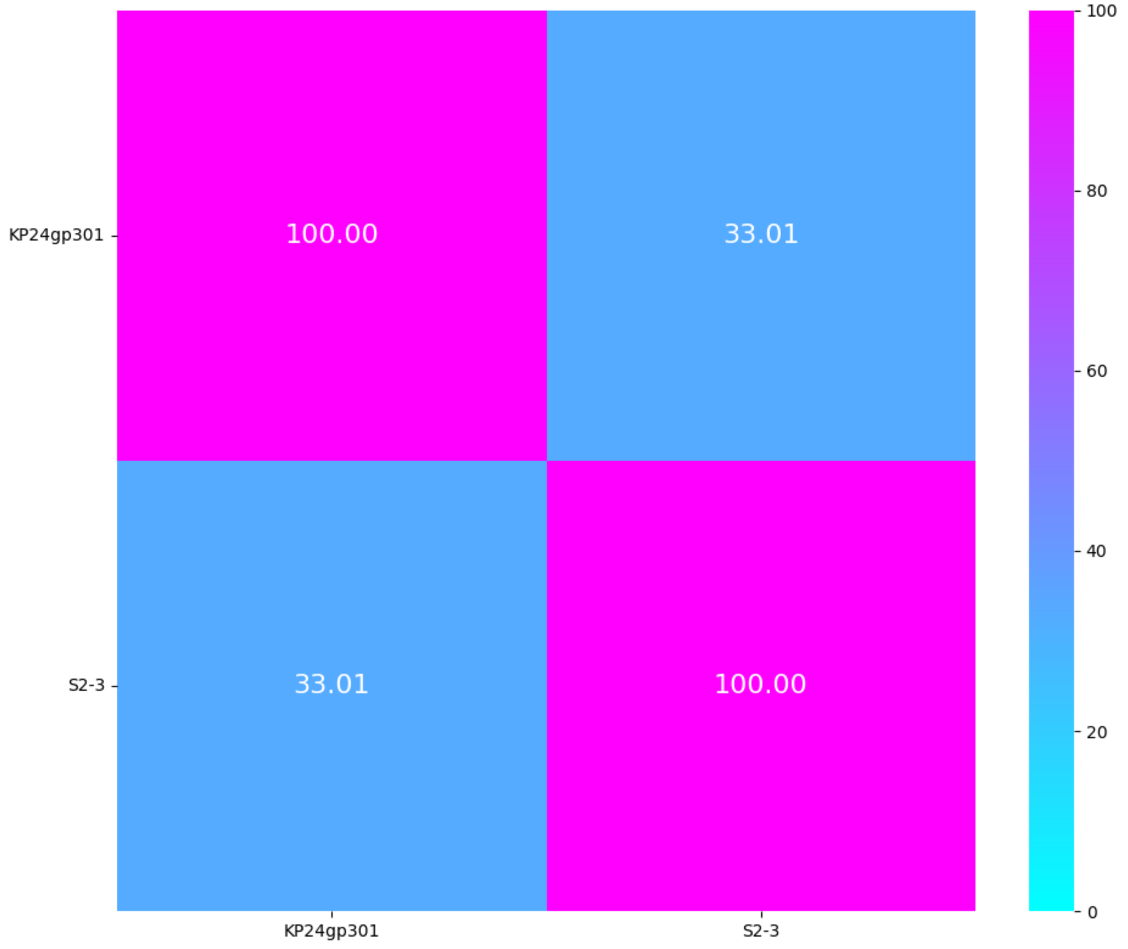

# K39, K46, K47

| K-type target | K39 | K46 | | K47 | | | |
|---|---|---|---|---|---|---|---|
| **Group** | Group 1 | Group 1 | Group 2 | Group 1 | | Group 2 | Group 3 |
| **Protein name** | KANgp227 | KP24gp308 | 248_38 | Dep42 | Dpo42 | P560dep | Dpo43 |
| **Leng** | 820 aa | 737 aa | 693 aa | 793 aa | 793 aa | 802 aa | 641 aa |
| **Number of proteins in the group** | 1 protein | 1 protein | 1 protein | 2 proteins | | 1 protein | 1 protein |
| **Predicted protein structure** | 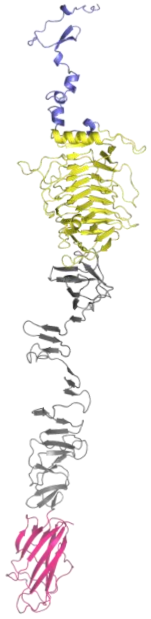 | 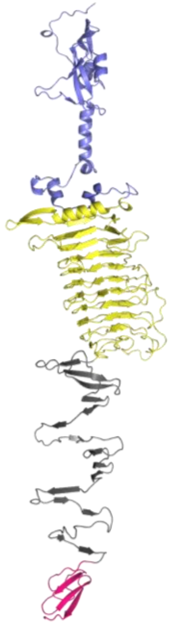 | 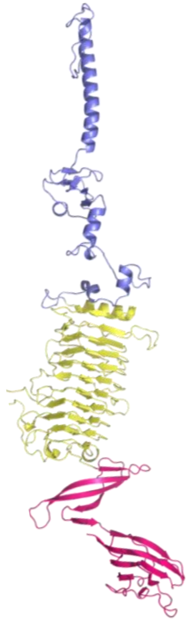 | 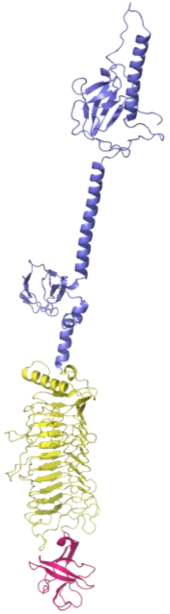 | 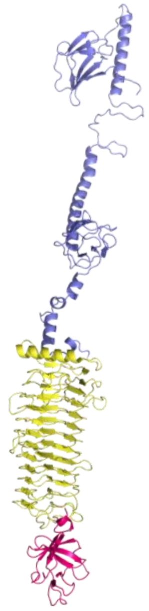 | 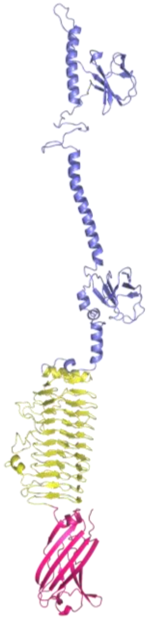 | 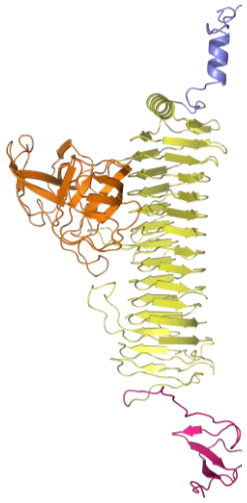 |

K46 group 1 + group 2
TM-score
Central

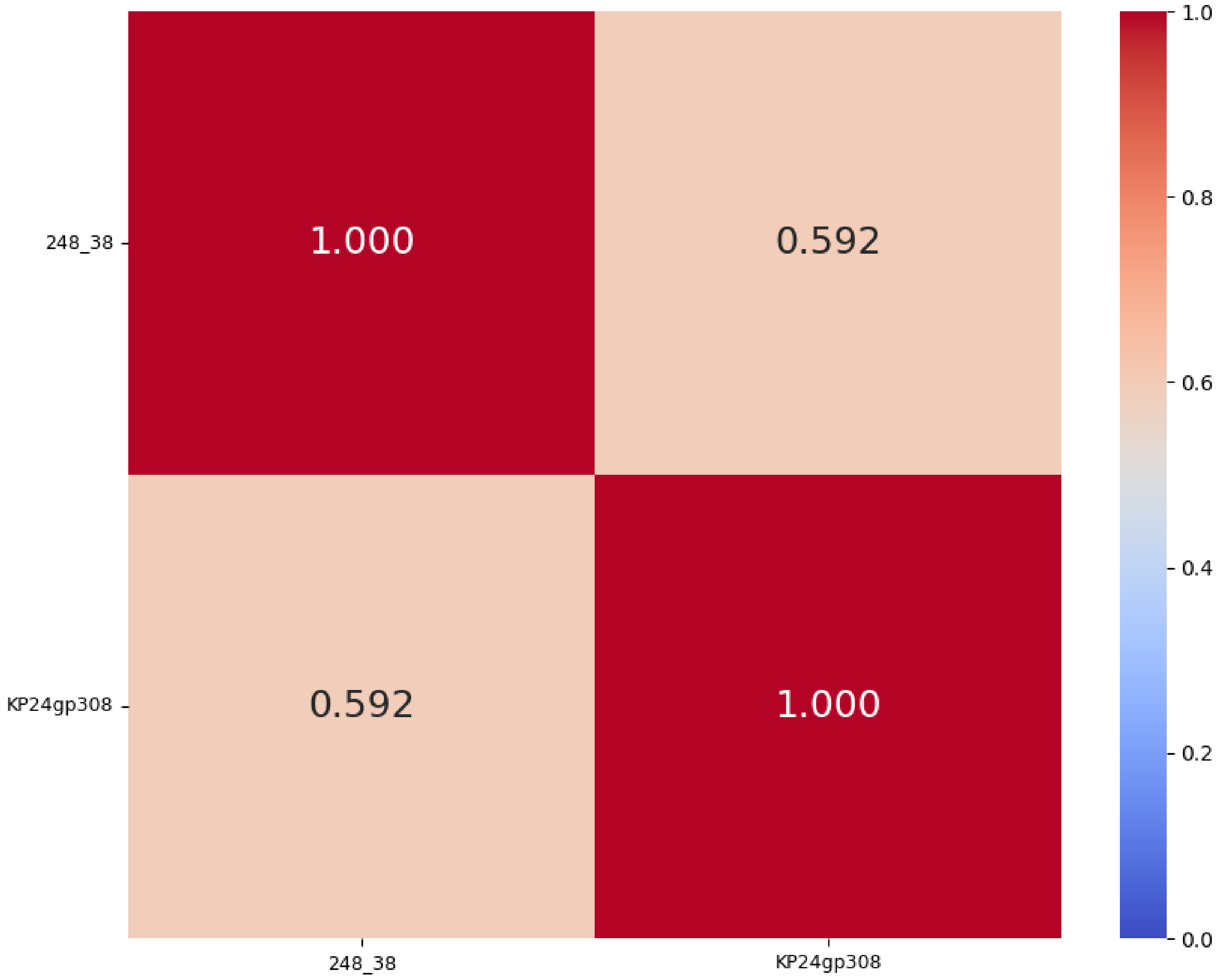

K47 group 1_2_3
TM-score
Central + C-term

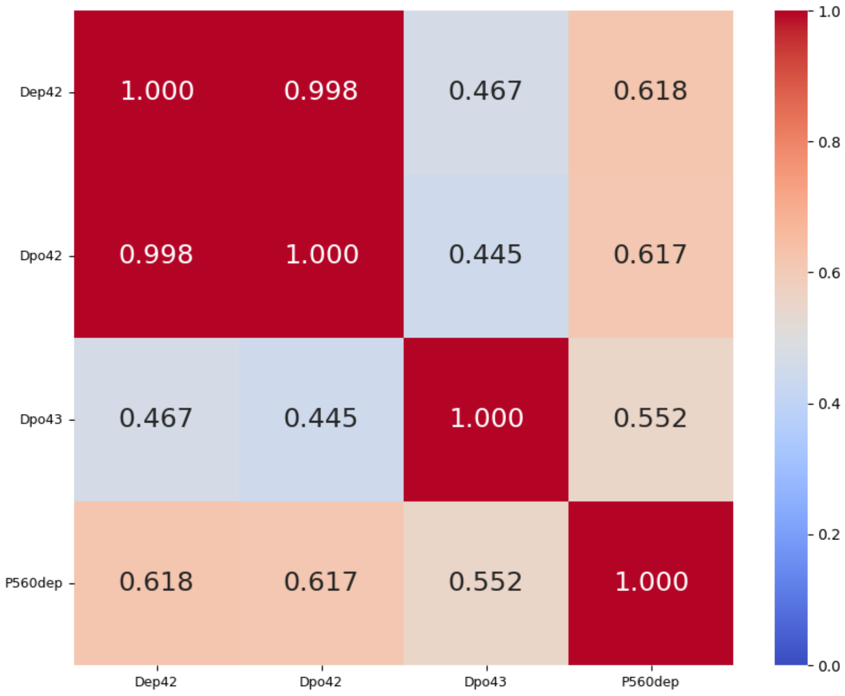


K47 group 1_2_3
%identity; cover 5-100%
Central + C-term

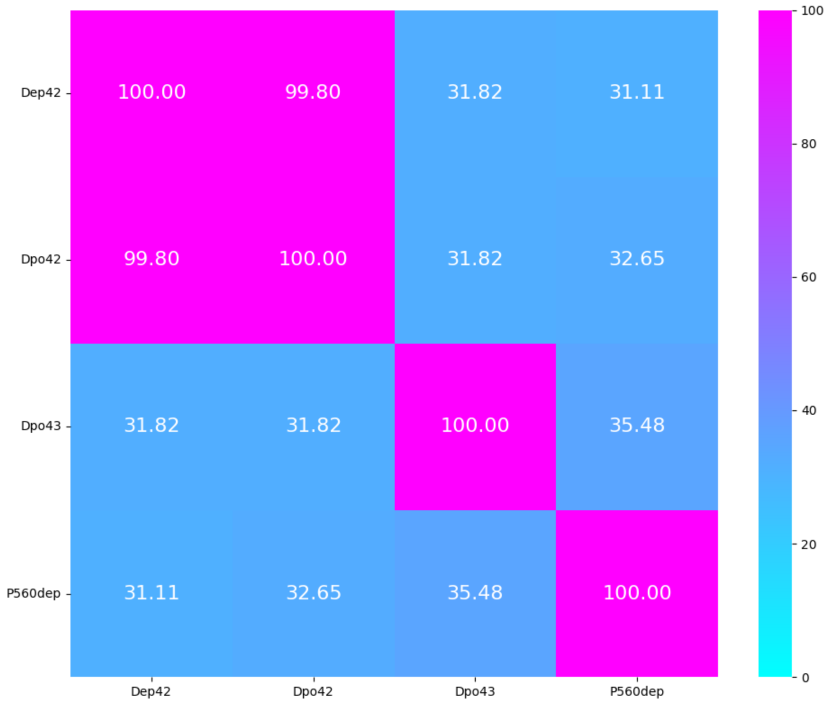


K47 group 1_2_3
TM-score
Central

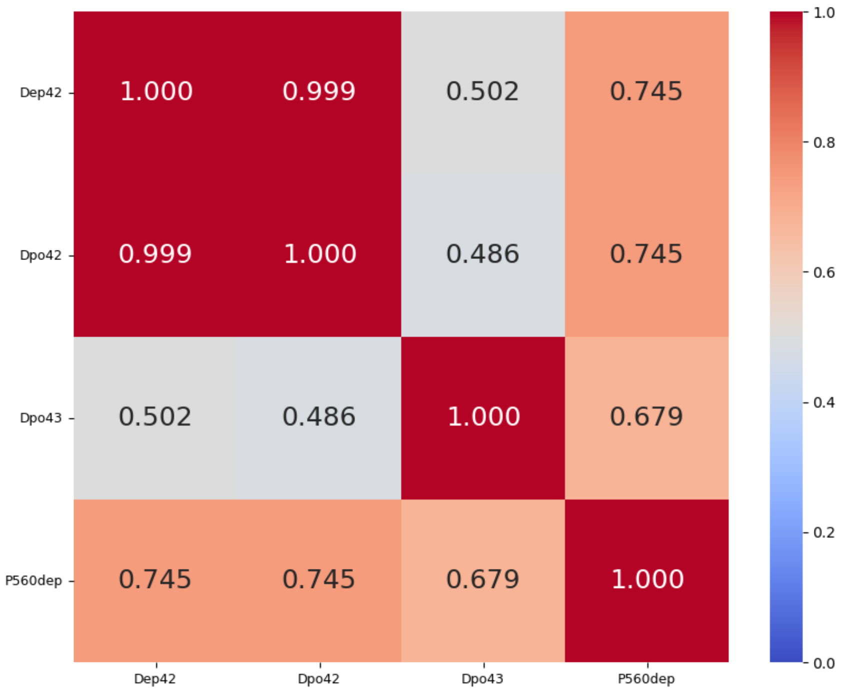

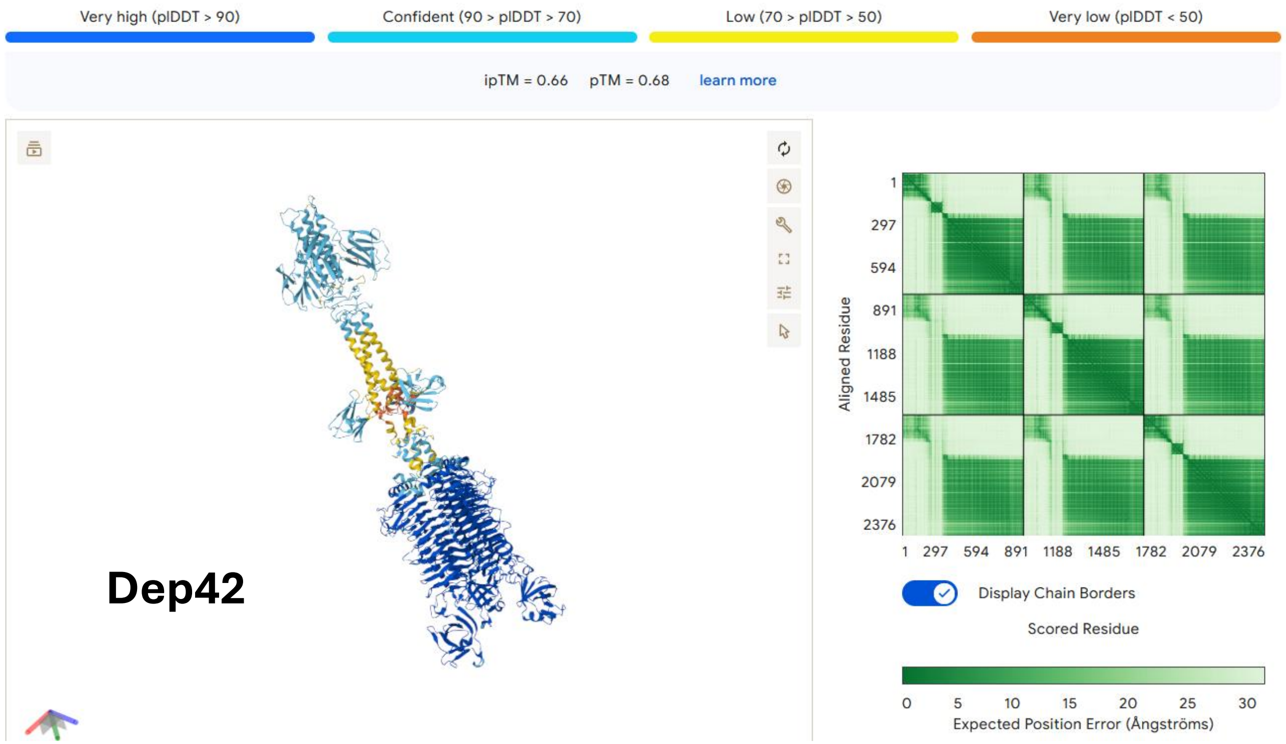
Very high (pIDDT > 90)
Confident (90 > pIDDT > 70)
Low (70 > pIDDT > 50)
Very low (pIDDT < 50)
ipTM = 0.66 pTM = 0.68 learn more
Dep42
Aligned Residue
1 297 594 891 1188 1485 1782 2079 2376
Display Chain Borders
Scored Residue
0 5 10 15 20 25 30
Expected Position Error (Ångströms)

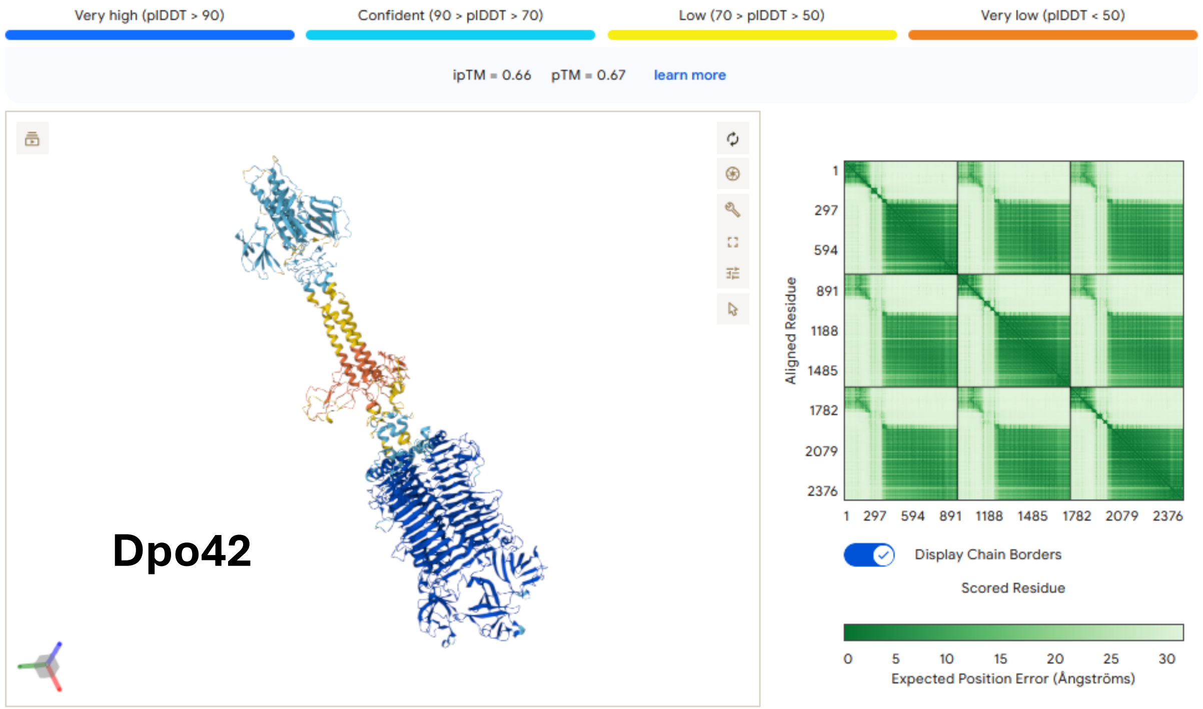
Very high (pIDDT > 90)
Confident (90 > pIDDT > 70)
Low (70 > pIDDT > 50)
Very low (pIDDT < 50)
ipTM = 0.66 pTM = 0.67 learn more
Dpo42
Aligned Residue
1 297 594 891 1188 1485 1782 2079 2376
Display Chain Borders
Scored Residue
0 5 10 15 20 25 30
Expected Position Error (Ångströms)

# K51, K52, K54, K56

| K-type target | K51 | K52 | K54 | K56 |
|---|---|---|---|---|
| **Group** | Group 1 | Group 1 | Group 1 | Group 1 |
| **Protein name** | GBH019_279 | 434_33 | gp531 | K56dep |
| **Lenght** | 809 aa | 859 aa | 895 aa | 678 aa |
| **Number of proteins in the group** | 1 protein | 1 protein | 1 protein | 1 protein |
| **Predicted protein structure** | 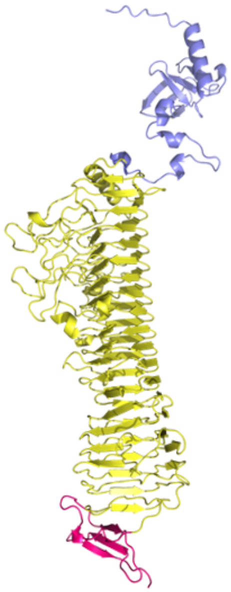 | 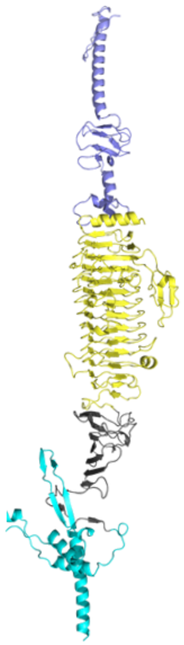 | 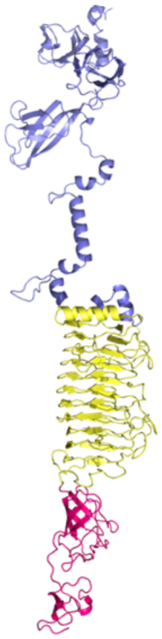 | 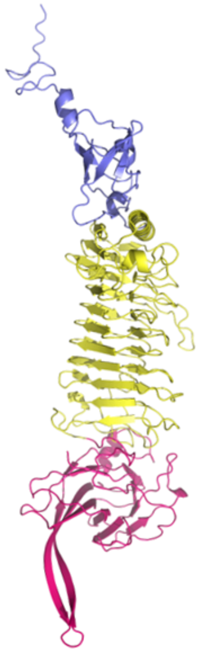 |

#### Cleavage site gp531

(Noreika et al. 2023)

**K54** CPS repeat unit

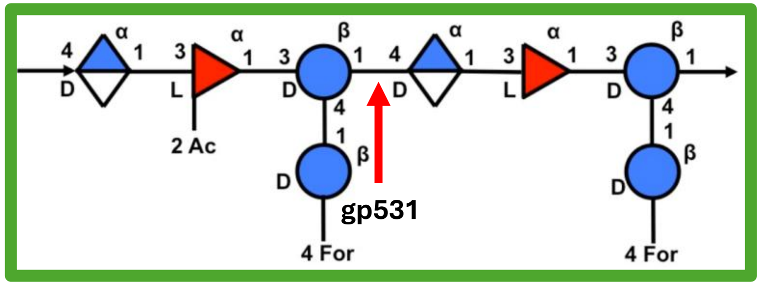


→4)-α-D-GlcpA-(1→3)-α-L-Fucp-(1→3)-β-D-Glcp-(1→4) -α-D-GlcpA-(1→3)-α-L-Fucp-(1→3)-β-D-Glcp-(1→
2 ↑ OAc
4 ↑ 1 β-D-Glcp 4 ↑ OFor
4 ↑ 1 β-D-Glcp 4 ↑ OFor

gp531

# **K57**/K68

| K-type target | K57* | K57* | K57/K68 | K57* | K57/K68 |
|---|---|---|---|---|---|
| **Group** | Group 1 | | | | |
| **Protein name** | Dep_kpv767 | Dep_kpv79 | gp157 | Dep_ZX1 | KLEO7gp24 |
| **Lenght** | 843 aa | 721 aa | 628 aa | 614 aa | 570 aa |
| **Number of proteins in the group** | 5 proteins | | | | |
| **Predicted protein structure** | 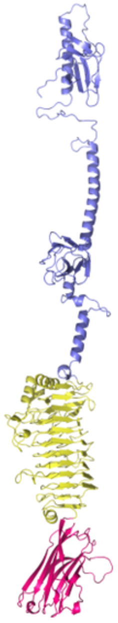 | 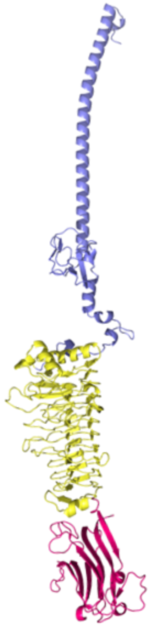 | 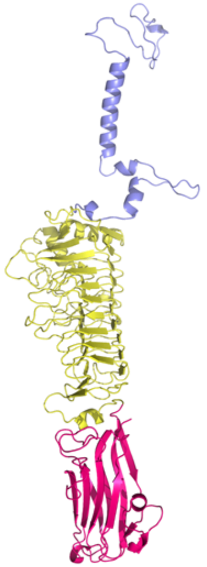 | 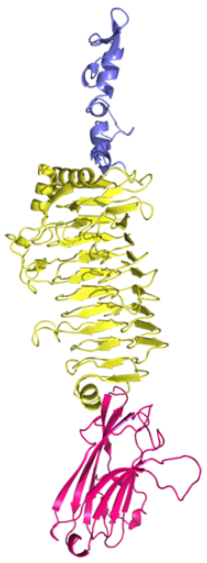 | 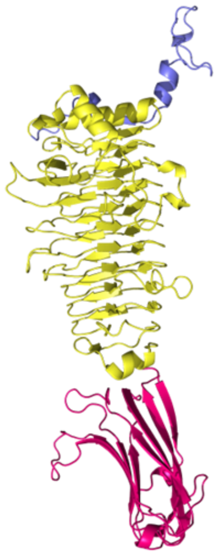 |

*protein was not tested on K68

#### Cleavage site Dep_kpv767 Dep_kpv79

(Volozhantsev et al. 2020)

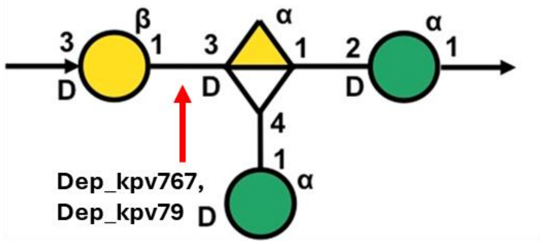


→3)-β-D-Gal*p*-(1→3)-α-D-Gal*p*A-(1→2)-α-D-Man*p*-(1→
4
↑
1
α-D-Man*p*

Dep_kpv767, Dep_kpv79

**K57**
CPS repeat unit

K57 group 1
TM-score
Central + C-term

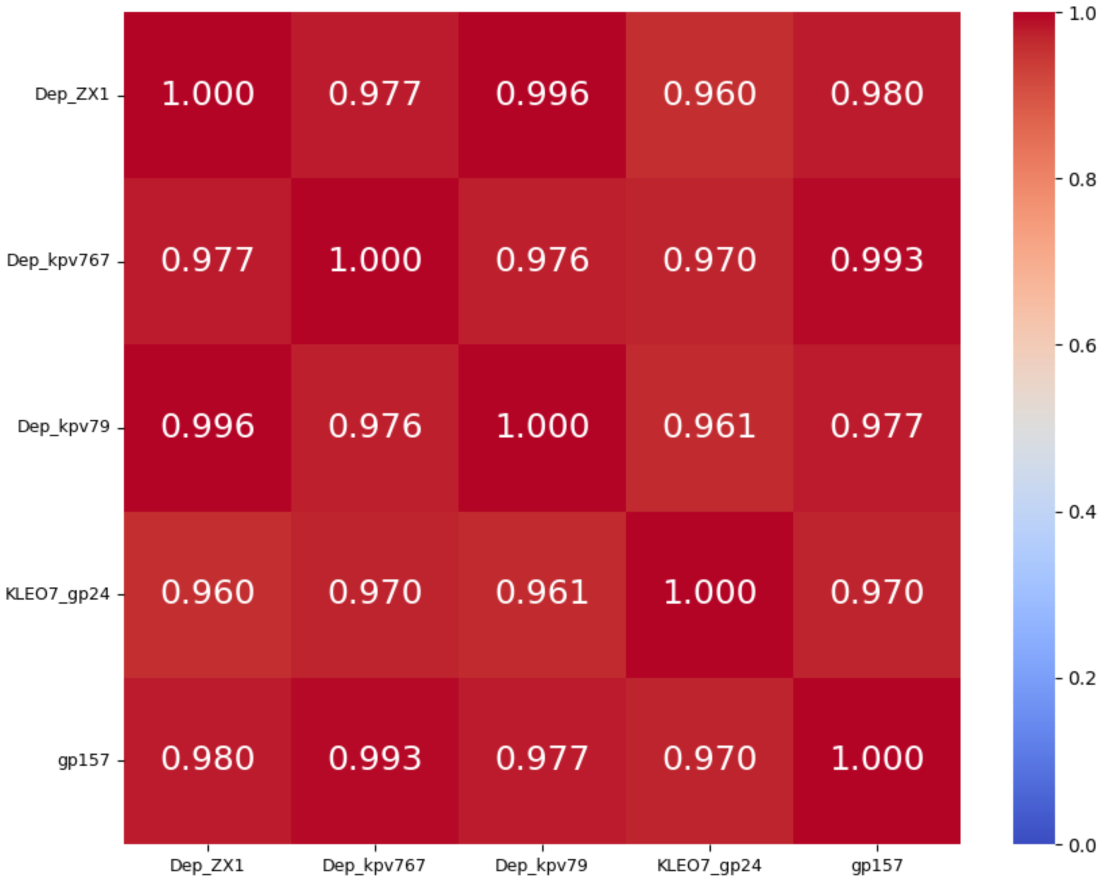


K57 group 1
%identity; cover ≥ 99%
Central + C-term

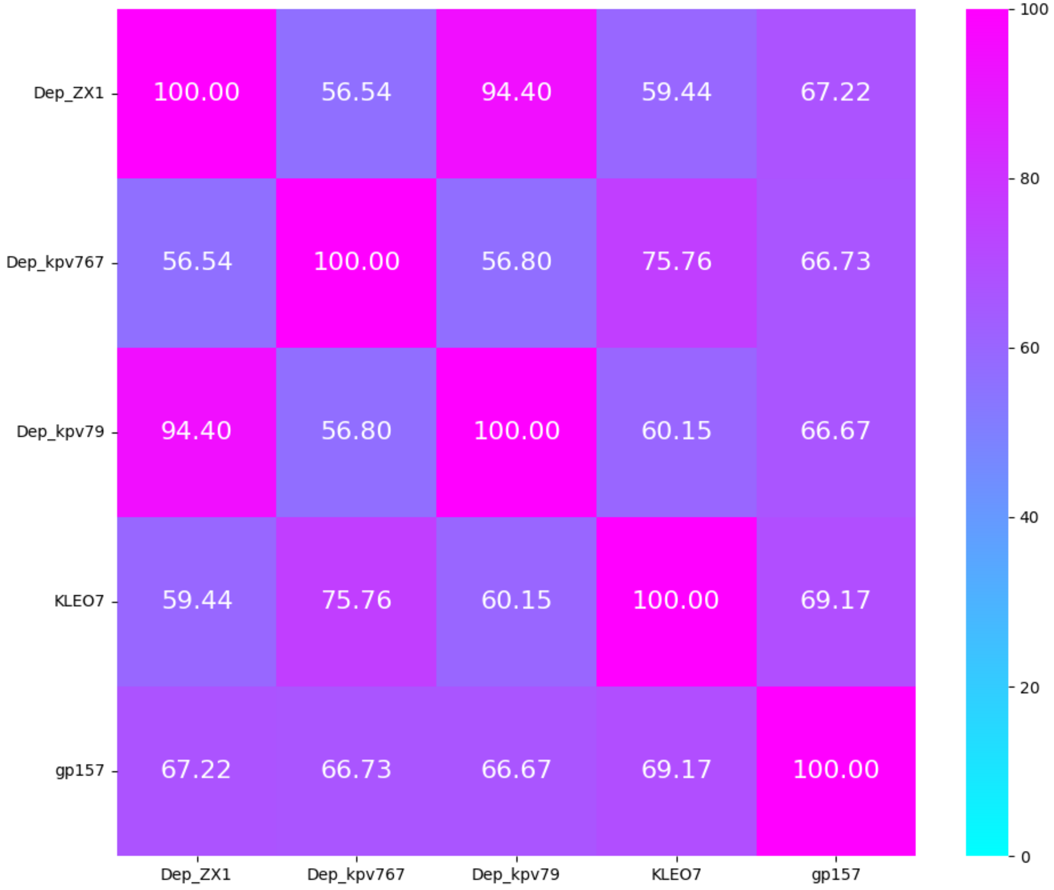

# K60, K61, K62

| K-type target | K60 | | K61 | K62 | | | |
|---|---|---|---|---|---|---|---|
| Group | Group 1 | | Group 1 | Group 1 | | Group 2 | |
| Protein name | 1723_59 | 1724_71 | KP24gp310 | 0367_12 | 0391_11 | K62-Dpo30 | 914_74 |
| Lenght | 951 aa | 843 aa | 679 aa | 851 aa | 839 aa | 692 aa | 666 aa |
| Number of proteins in the group | 2 proteins | | 1 protein | 2 proteins | | 2 proteins | |
| Predicted protein structure | 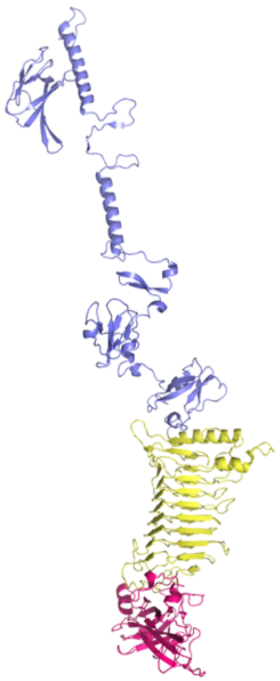 | 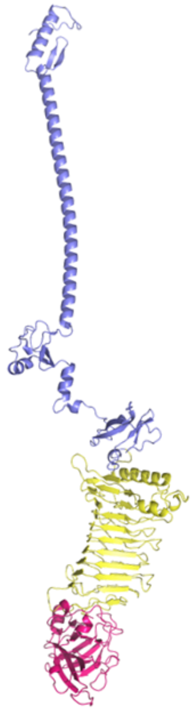 | 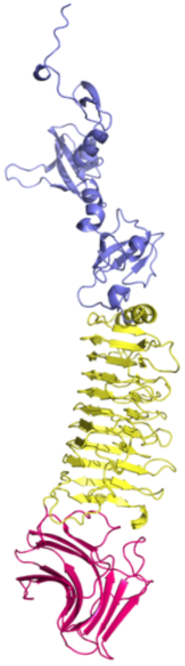 | 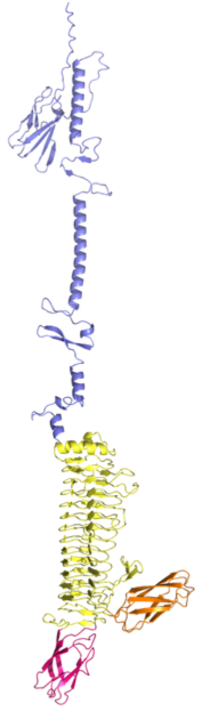 | 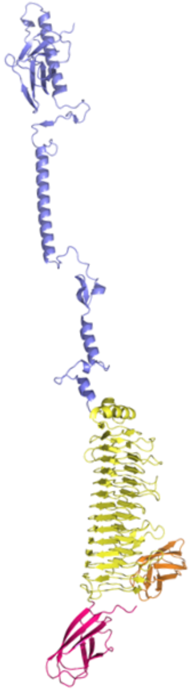 | 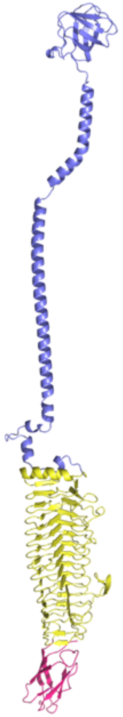 | 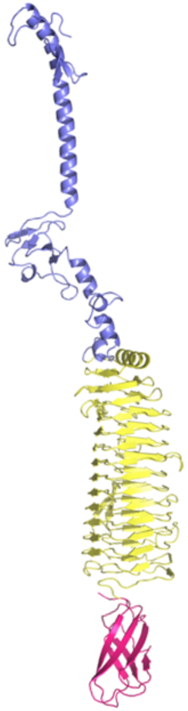 |

K60 group 1
TM-score
Central + C-term

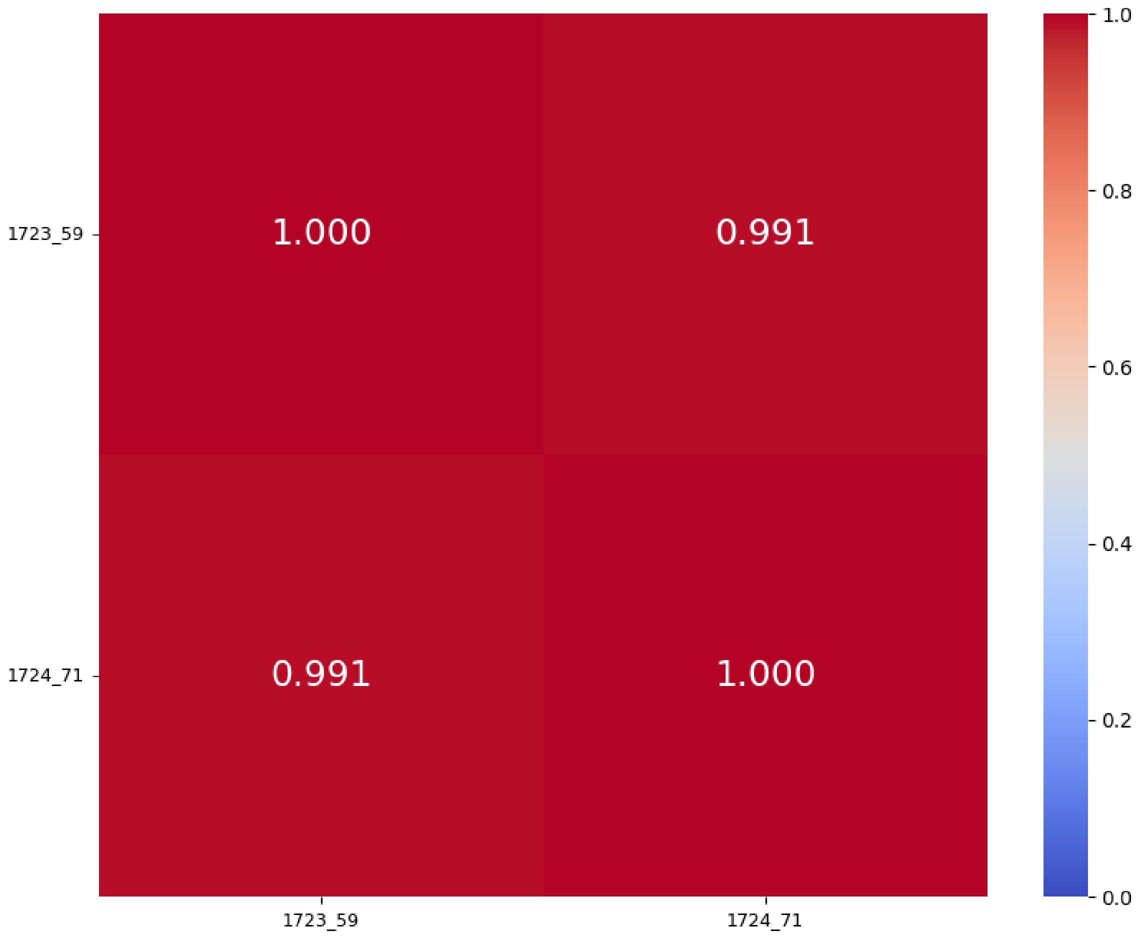


K60 group 1
%identity; cover ≥ 99%
Central + C-term

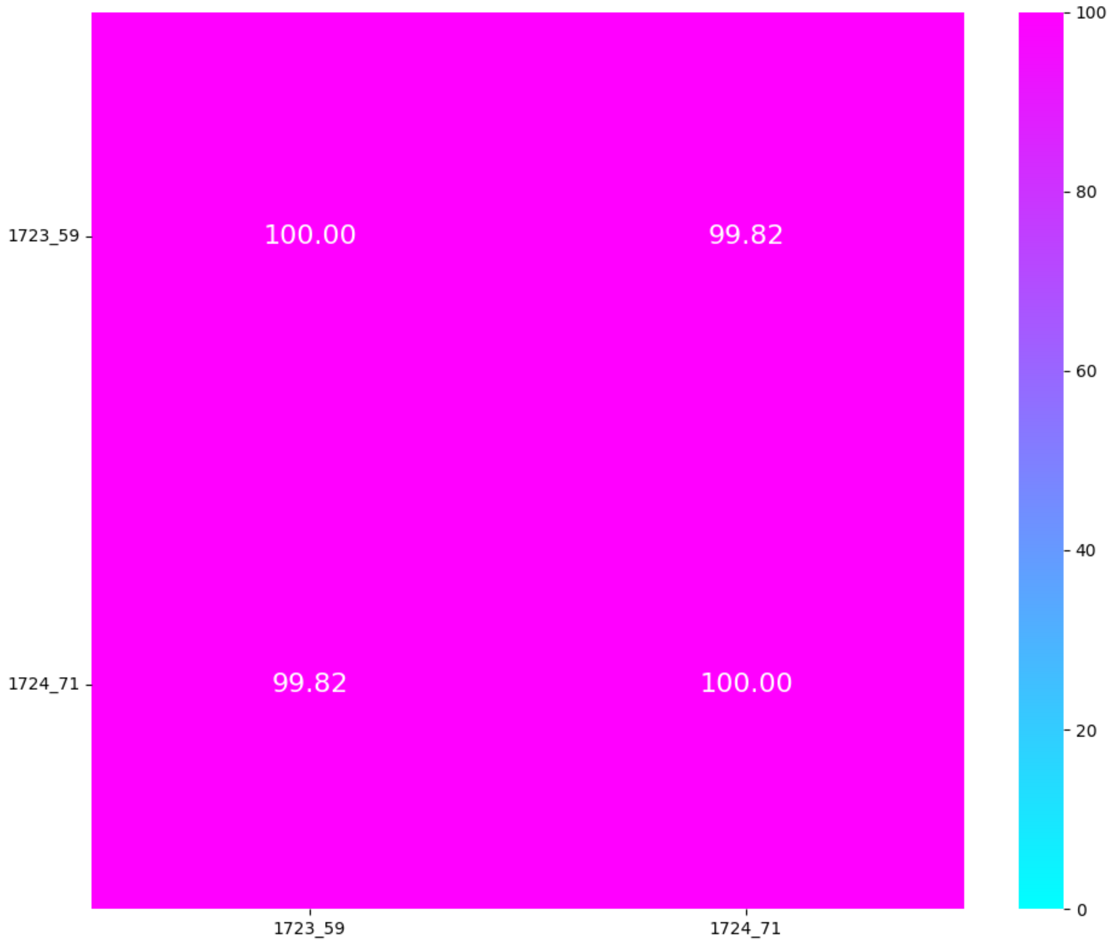

K62 group 1
TM-score
Central + C-term

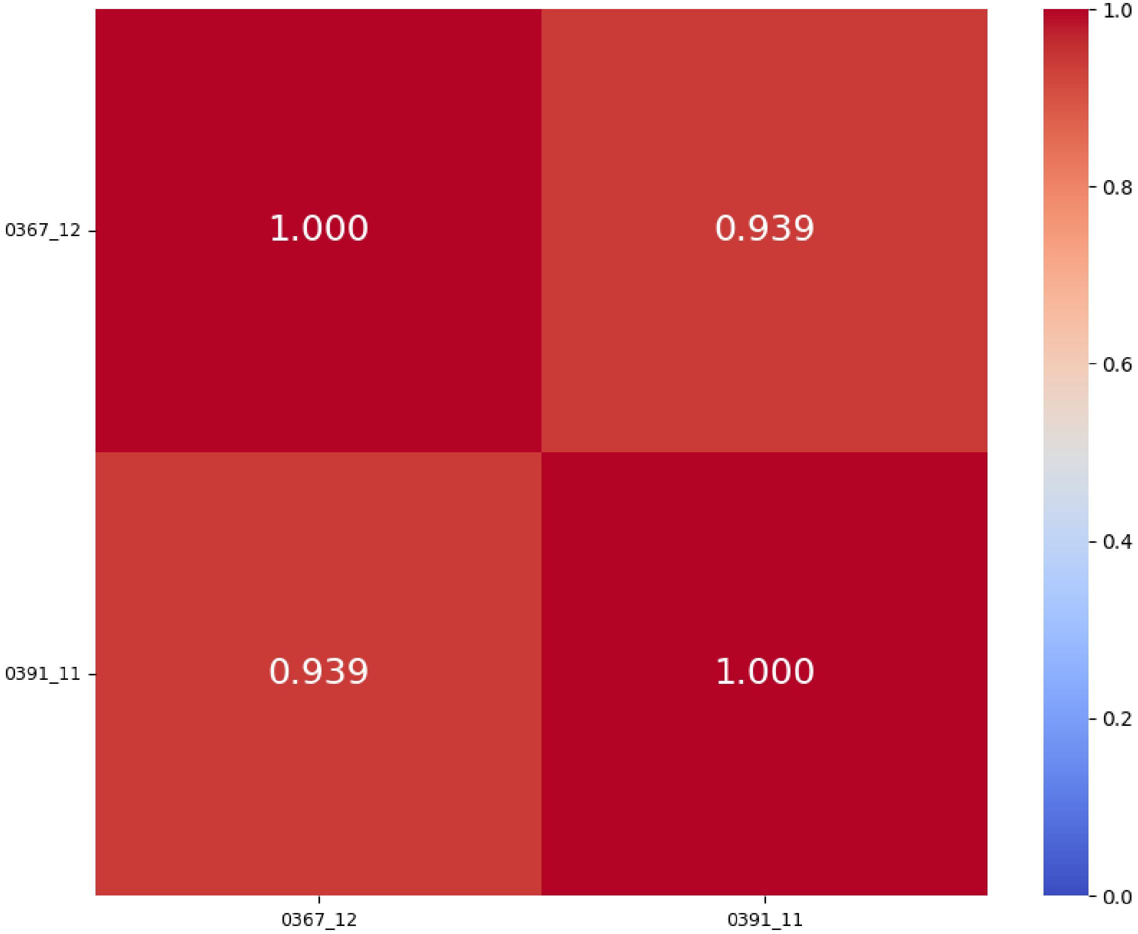


K62 group 1
%identity; cover ≥ 99%
Central + C-term

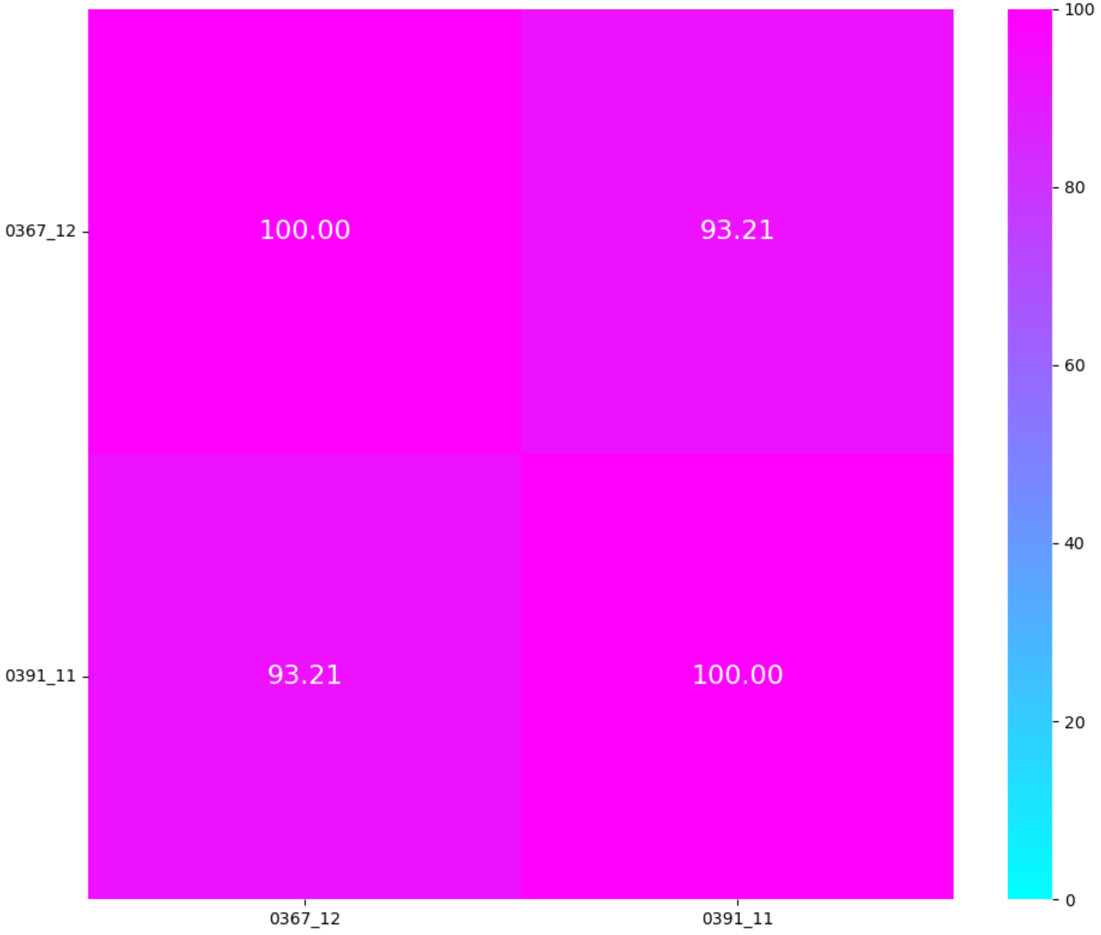

K62 group 2
TM-score
Central + C-term

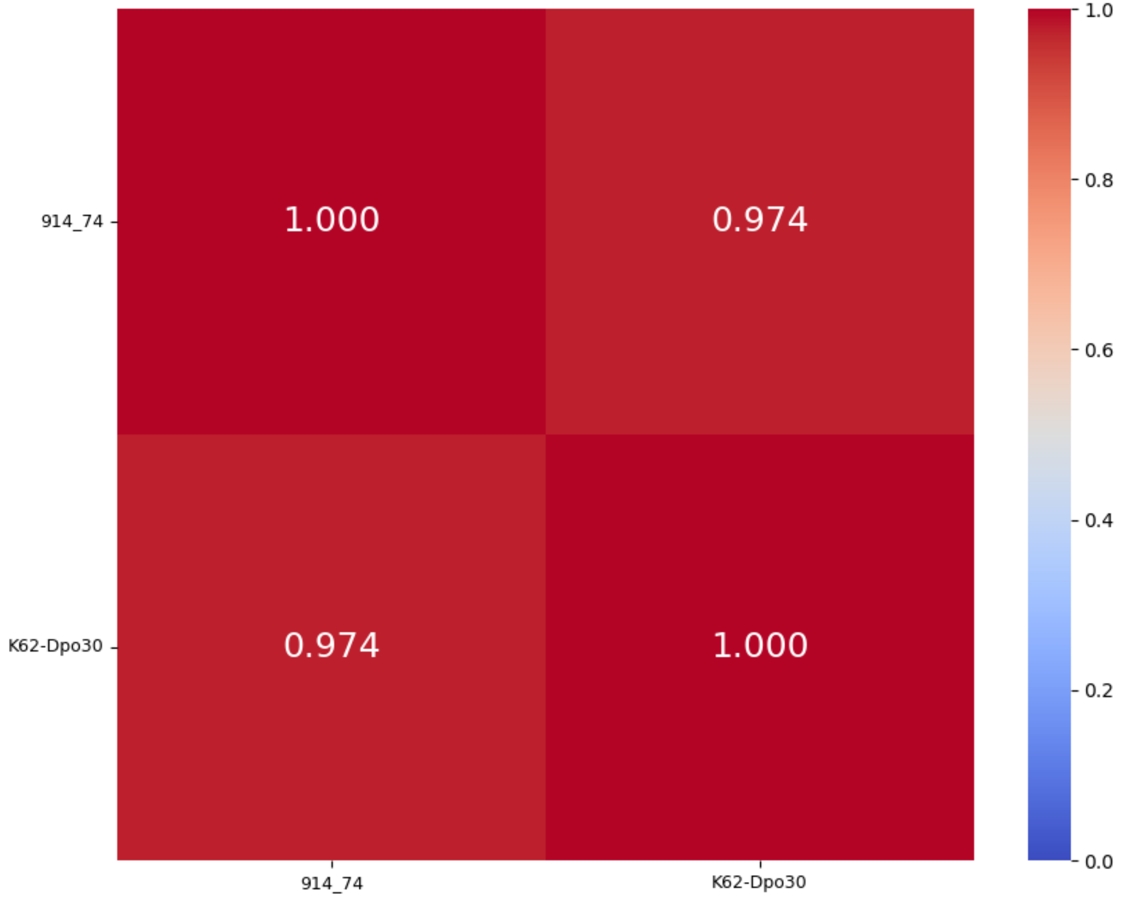


K62 group 2
%identity; cover ≥ 93%
Central + C-term

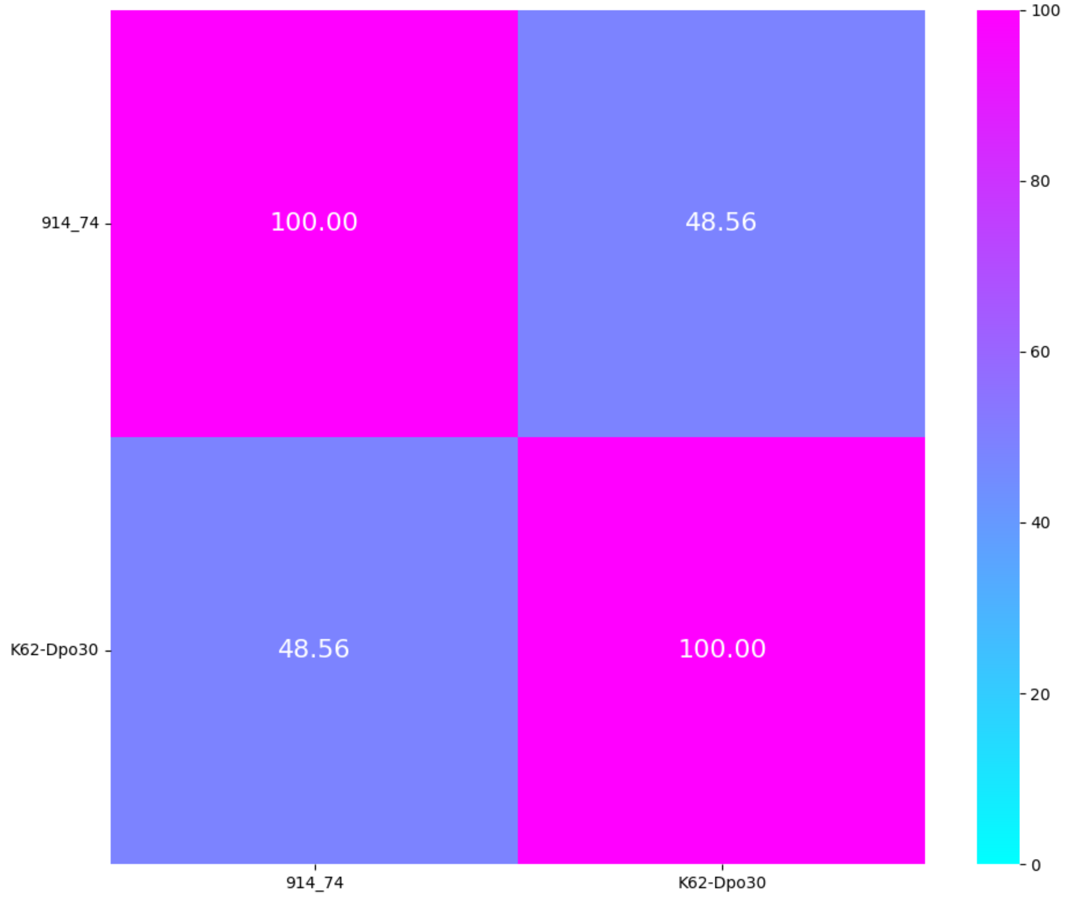

# K63

| K-type target | K63 | |
|---|---|---|
| Group | Group 1 | |
| Protein name | KP36gp50 | KP34gp57 |
| Lenght | 883 aa | 630 aa |
| Number of proteins in the group | 2 proteins | |
| Predicted protein structure | 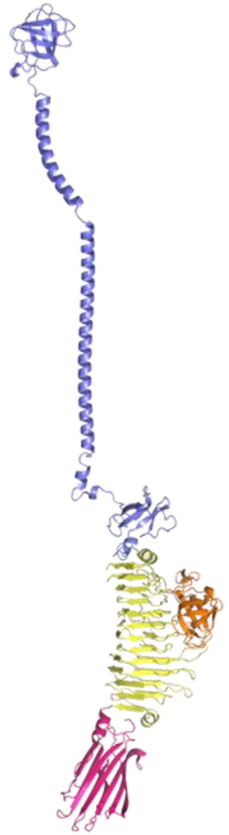 | 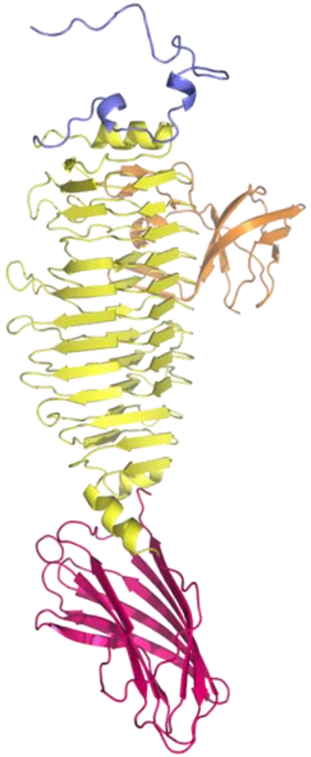 |

#### Cleavage site KP34gp57

(Maciejewska et al. 2023)

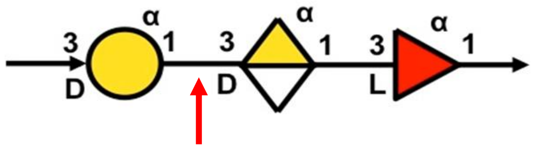


→3)-α-D-Gal*p*-(1→3)-α-D-Gal*p*A-(1→3)-α-L-Fuc*p*-(1→

KP34gp57

**K63**

CPS repeat unit

Active Site Residues **KP34gp57**

**Glu266/Glu300**

K63 group 1
TM-score
Central + C-term

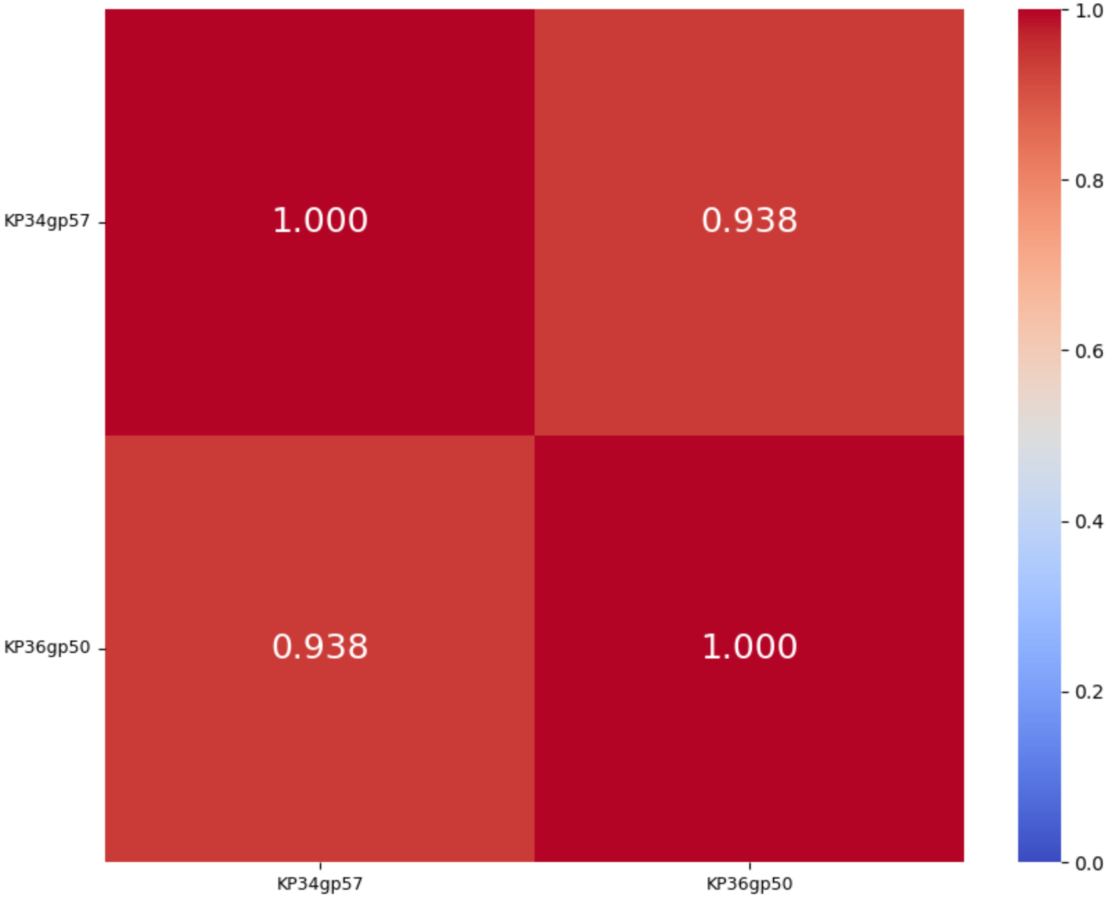


K63 group 1
%identity; cover ≥ 96%
Central + C-term

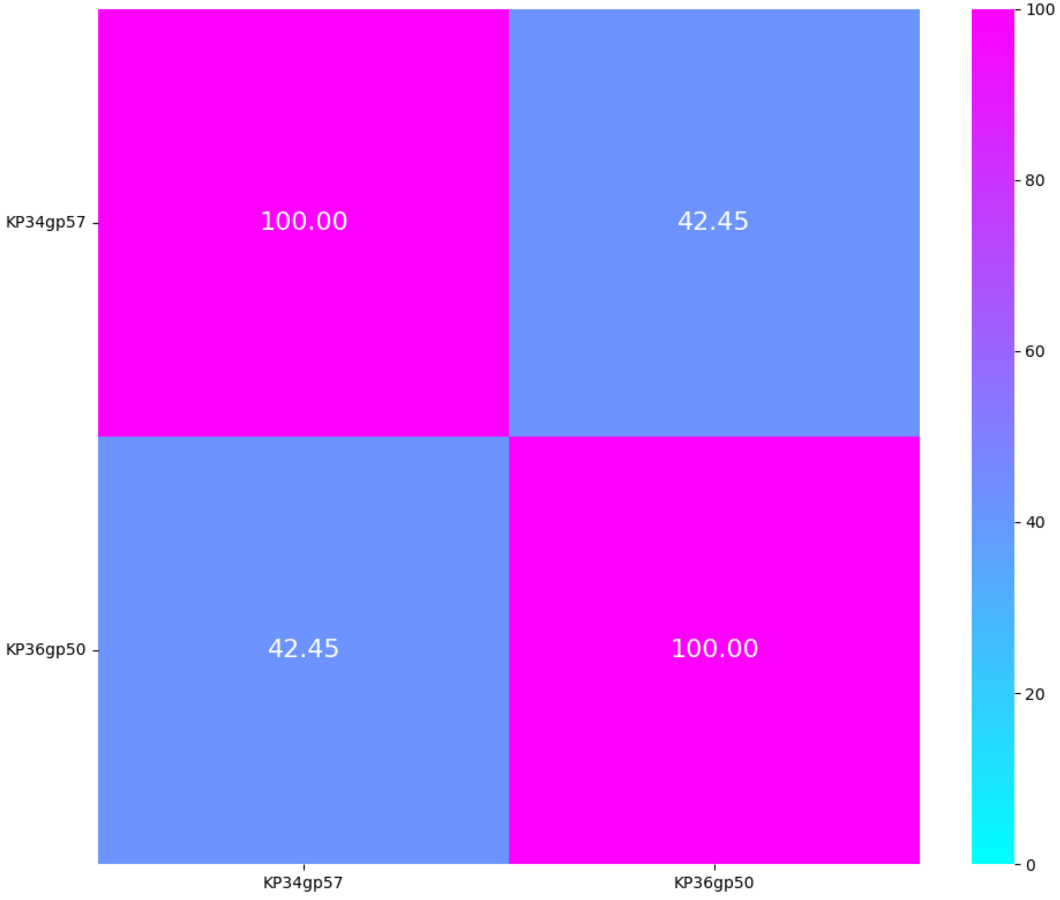

# K64

| K-type target | K64 | | | | |
|---|---|---|---|---|---|
| **Group** | Group 1 | | | Group 2 | |
| **Protein name** | P510dep | K64-ORF41 | S2-5 | 1091_44 | KP24gp303 |
| **Lenght** | 1017 aa | 1017 aa | 996 aa | 880 aa | 661 aa |
| **Number of proteins in the group** | 3 proteins | | | 2 proteins | |
| **Predicted protein structure** | 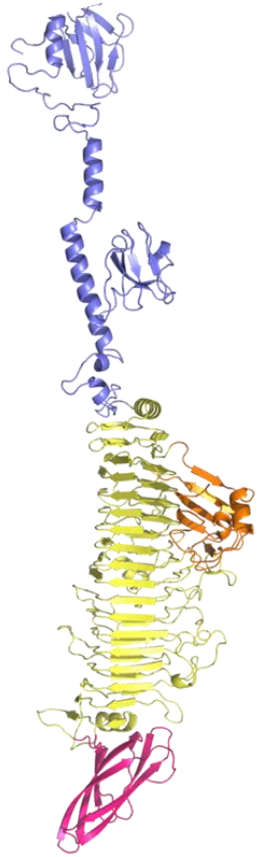 | 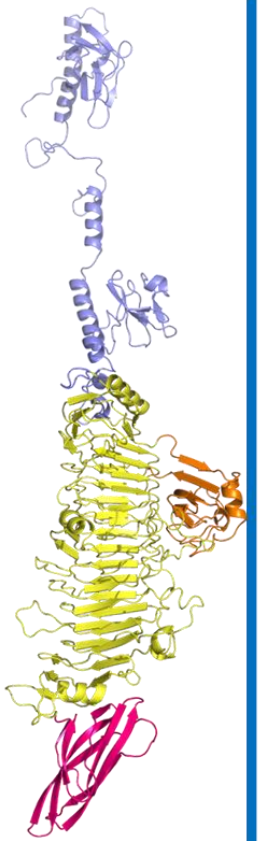 | 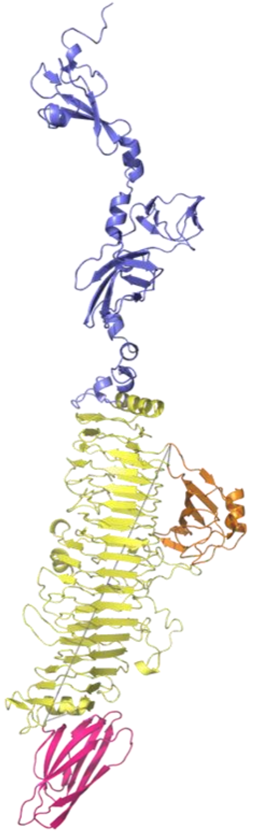 | 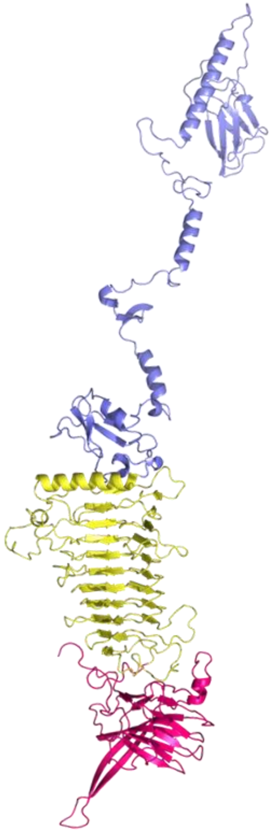 | 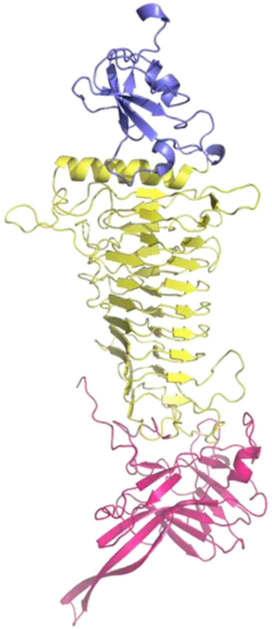 |

Active Site Residues
**K64-ORF41**
**Tyr528, His574, Arg628**

K64 group 1
TM-score
Central + C-term

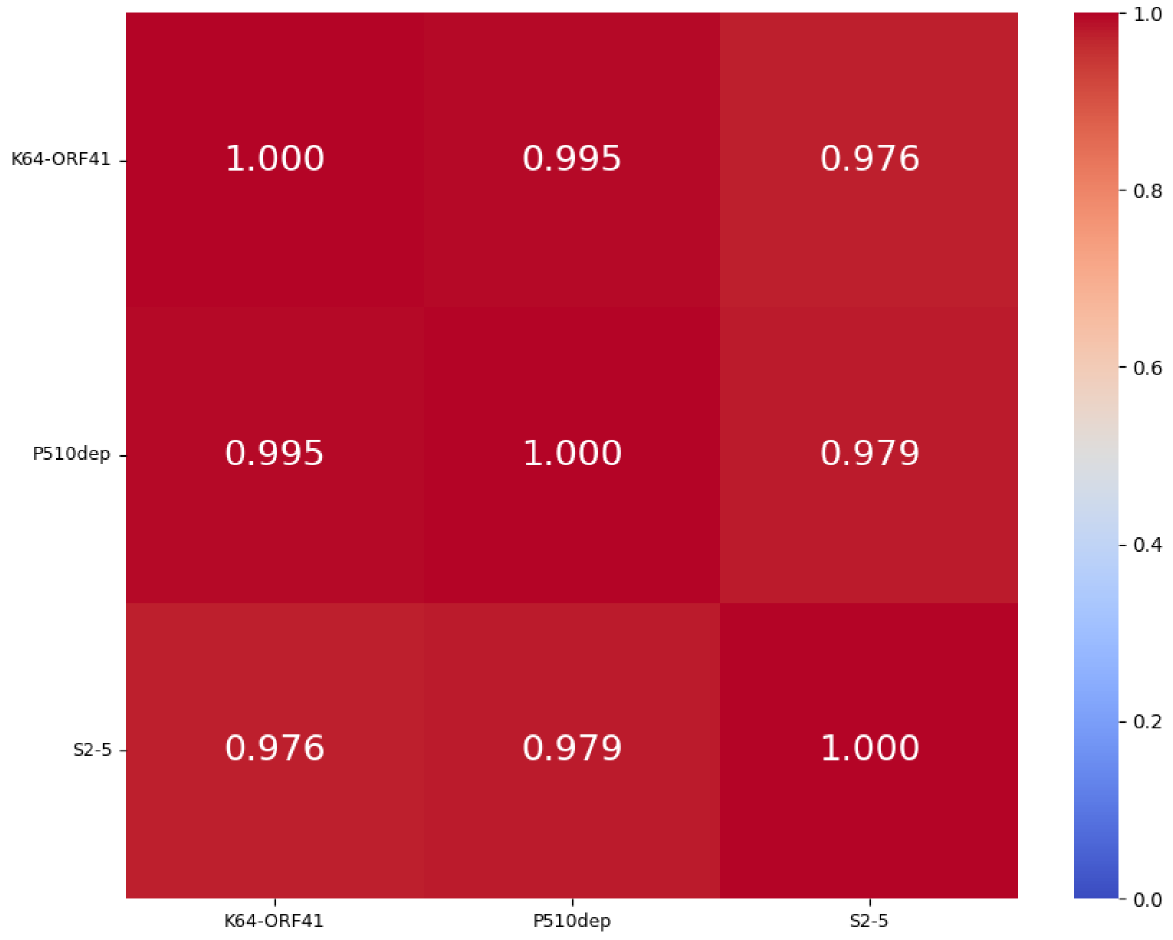


K64 group 1
%identity; cover ≥ 99%
Central + C-term

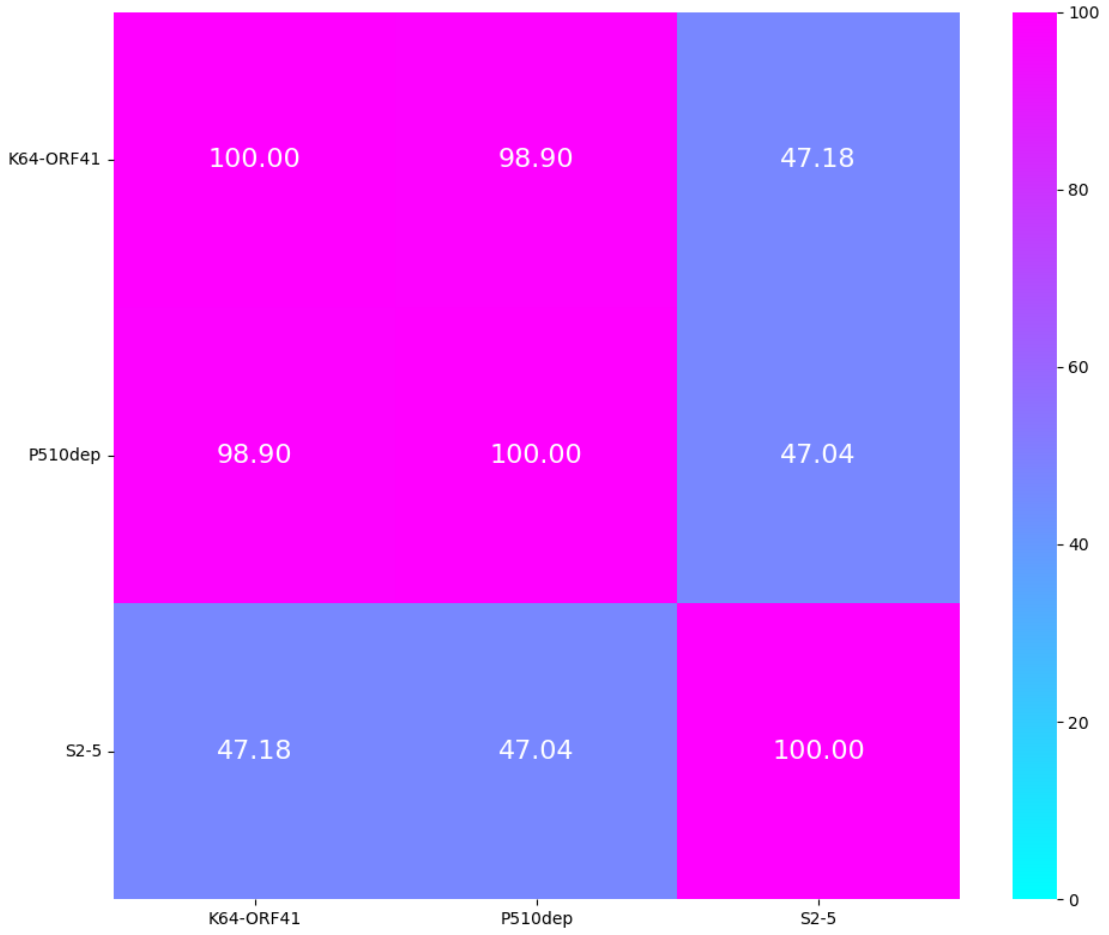

K64 group 2
TM-score
Central + C-term

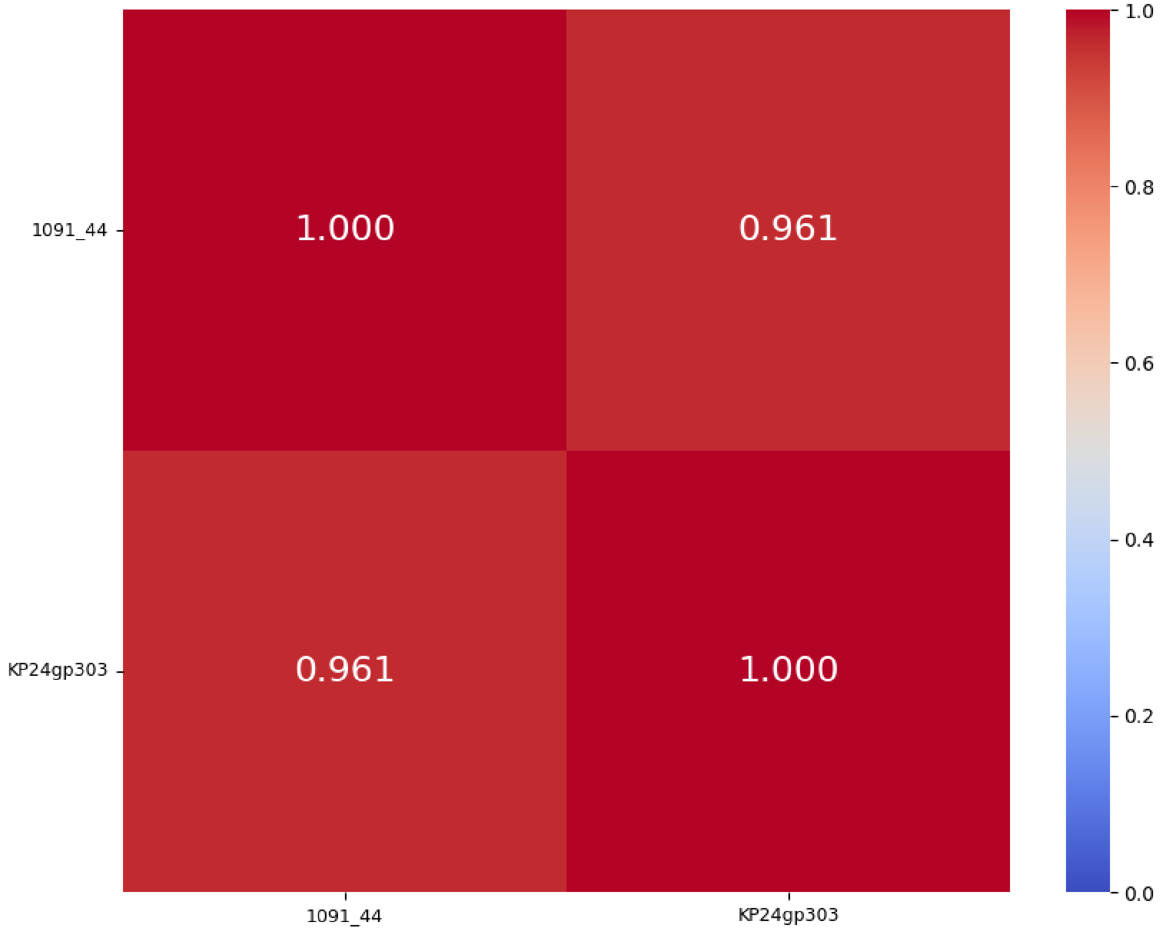


K64 group 2
%identity; cover ≥ 99%
Central + C-term

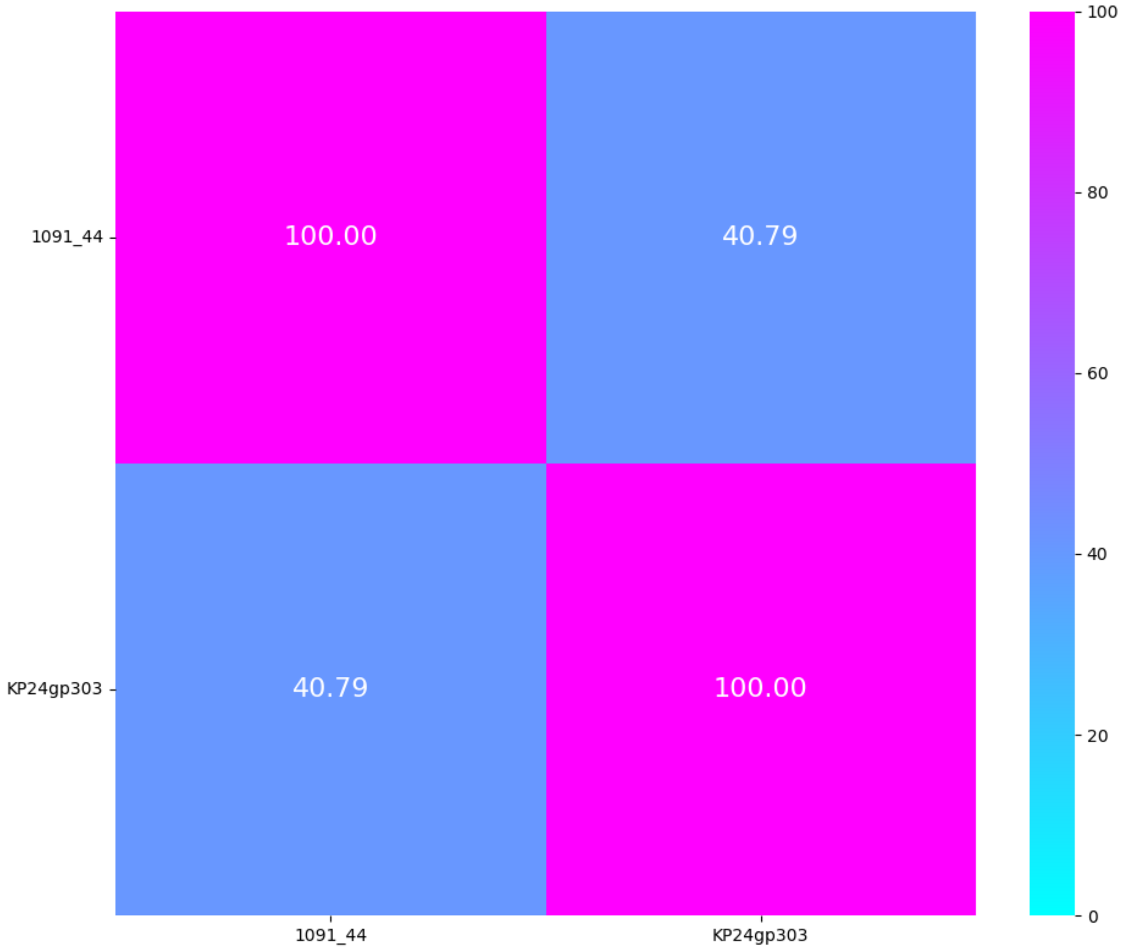

# KL101, KL102, KL103, KL104

| K-type target | KN1 = KL101 | | KN2 = KL102 | KN3 = KL103 | KN4 = KL104 | |
|---|---|---|---|---|---|---|
| **Group** | Group 1 | | Group 1 | Group 1 | Group 1 | Group 2 |
| **Protein name** | KN1dep | Dp42 | ORF96 | KN3dep | KN4dep | S1-2 |
| **Lenght** | 820 aa | 820 aa | 1245 aa | 792 aa | 850 aa | 736 aa |
| **Number of proteins in the group** | 2 proteins | | 1 protein | 1 protein | 1 protein | 1 protein |
| **Predicted protein structure** | 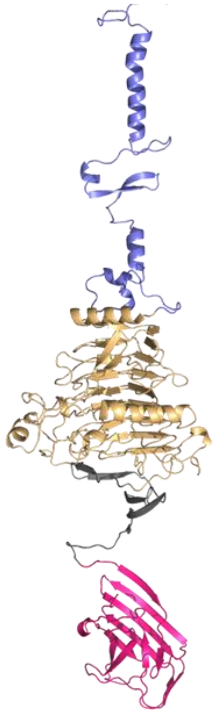 | 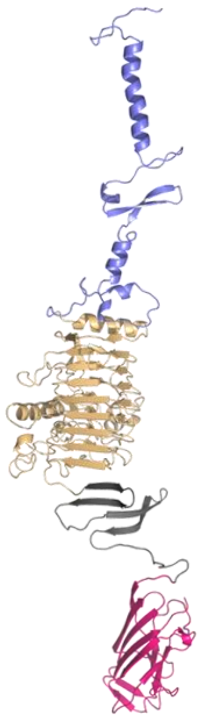 | 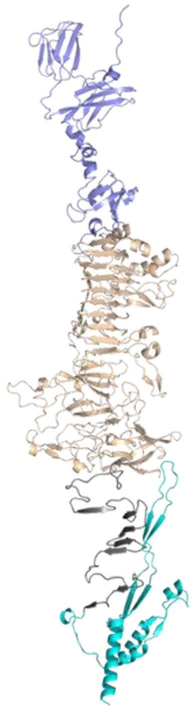 | 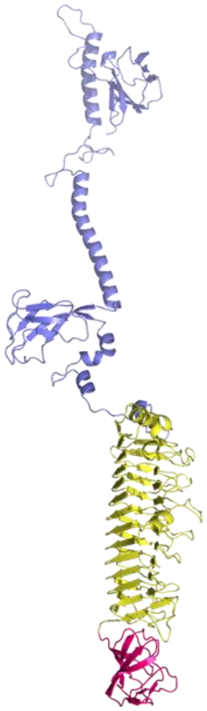 | 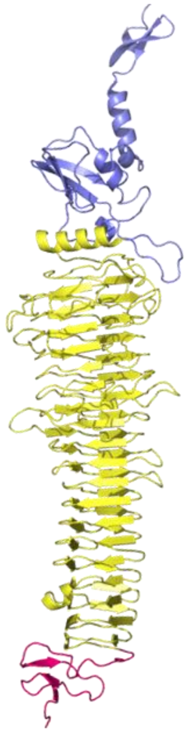 | 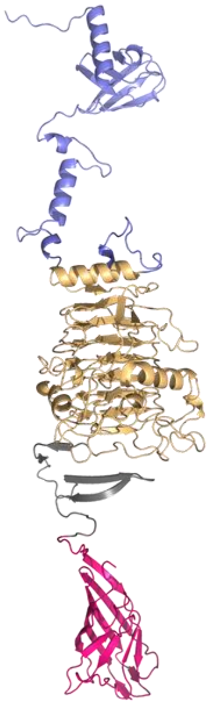 |

KN1_KL101 group 1
TM-score
Central + C-term

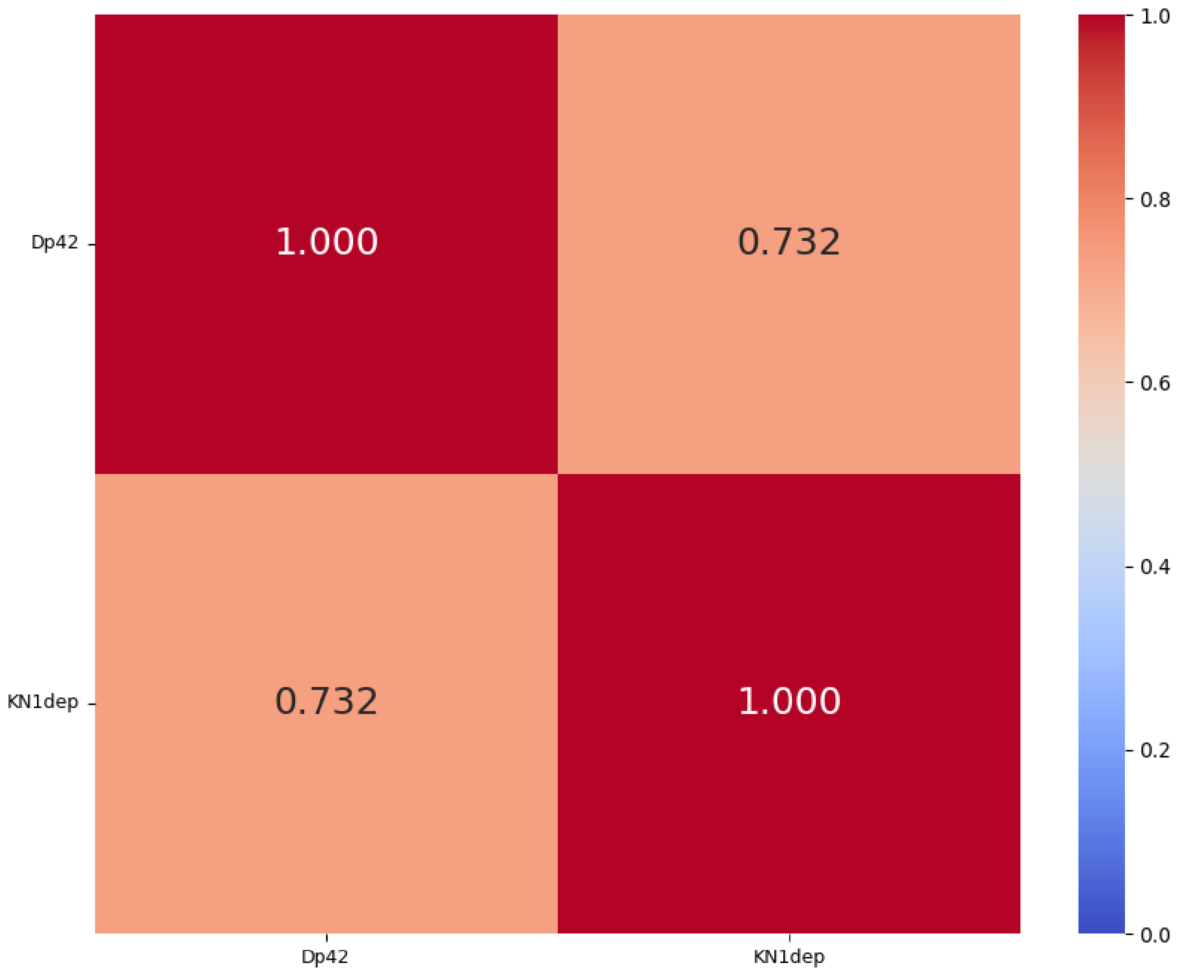


KN1_KL101 group 1
%identity; cover ≥ 99%
Central + C-term

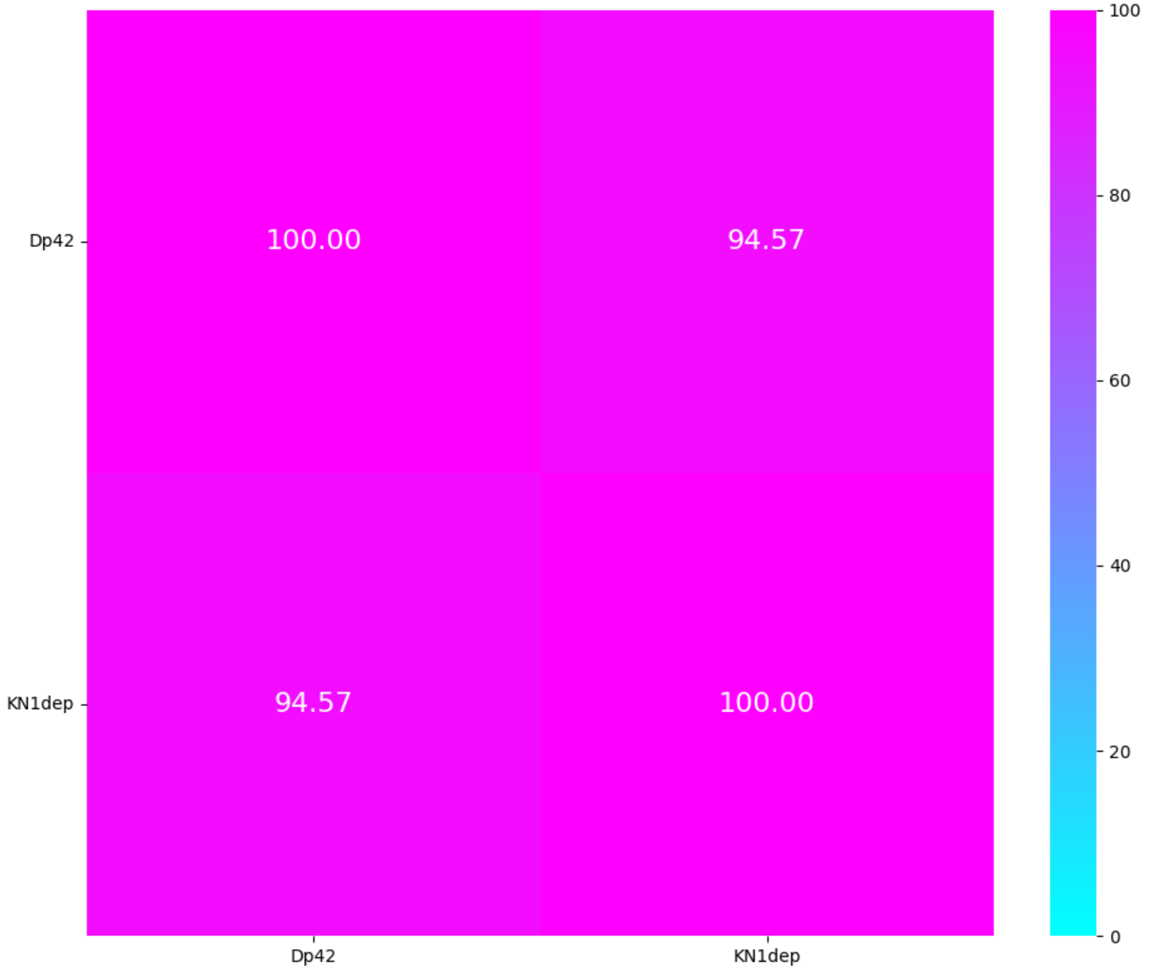

# KL105, KL108

| Serotype | KN5 = KL105 | KL108 | |
|---|---|---|---|
| Group | Group 1 | Group 1 | |
| Protein name | S2-1 | Dep108.2 | Dep108.1 |
| Lenght | 1193 aa | 592 aa | 590 aa |
| Number of proteins in the group | 1 protein | 2 proteins | |
| Predicted protein structure | 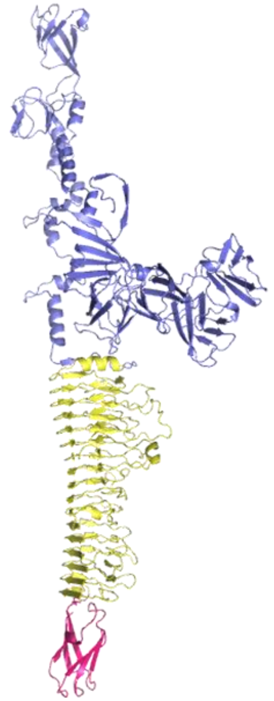 | 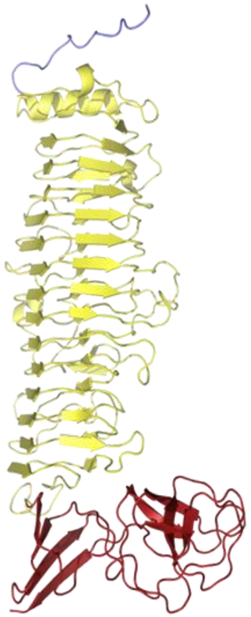 | 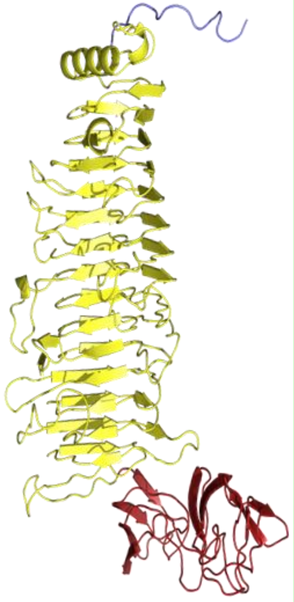 |

#### Cleavage site Dep108.1, Dep108.2

(Kasimova et al. 2023)

**KL108**
CPS repeat unit

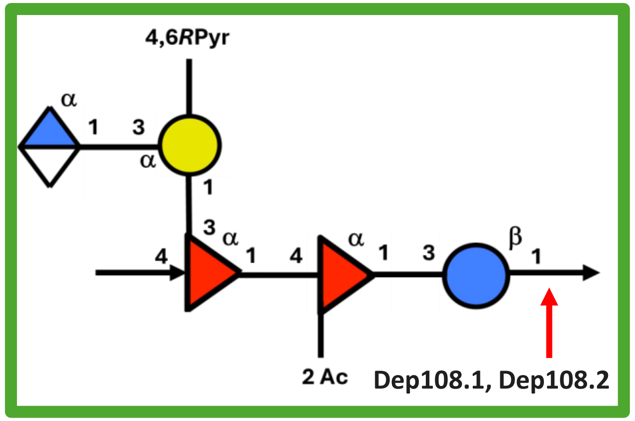


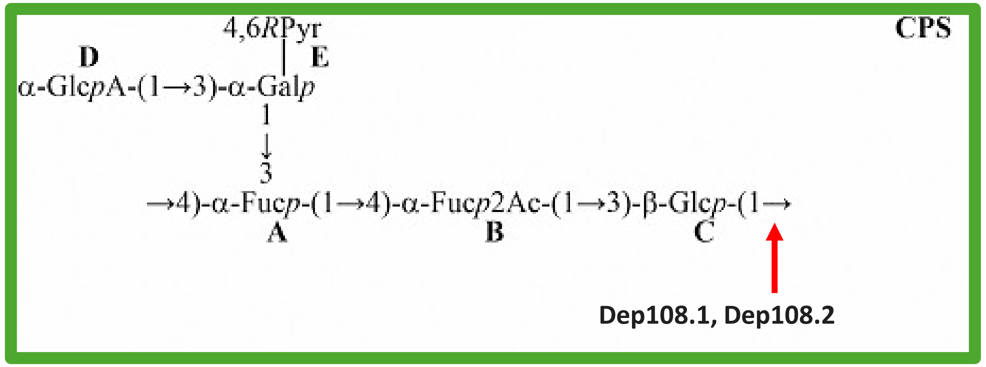

KL108 group 1
TM-score
Central + C-term

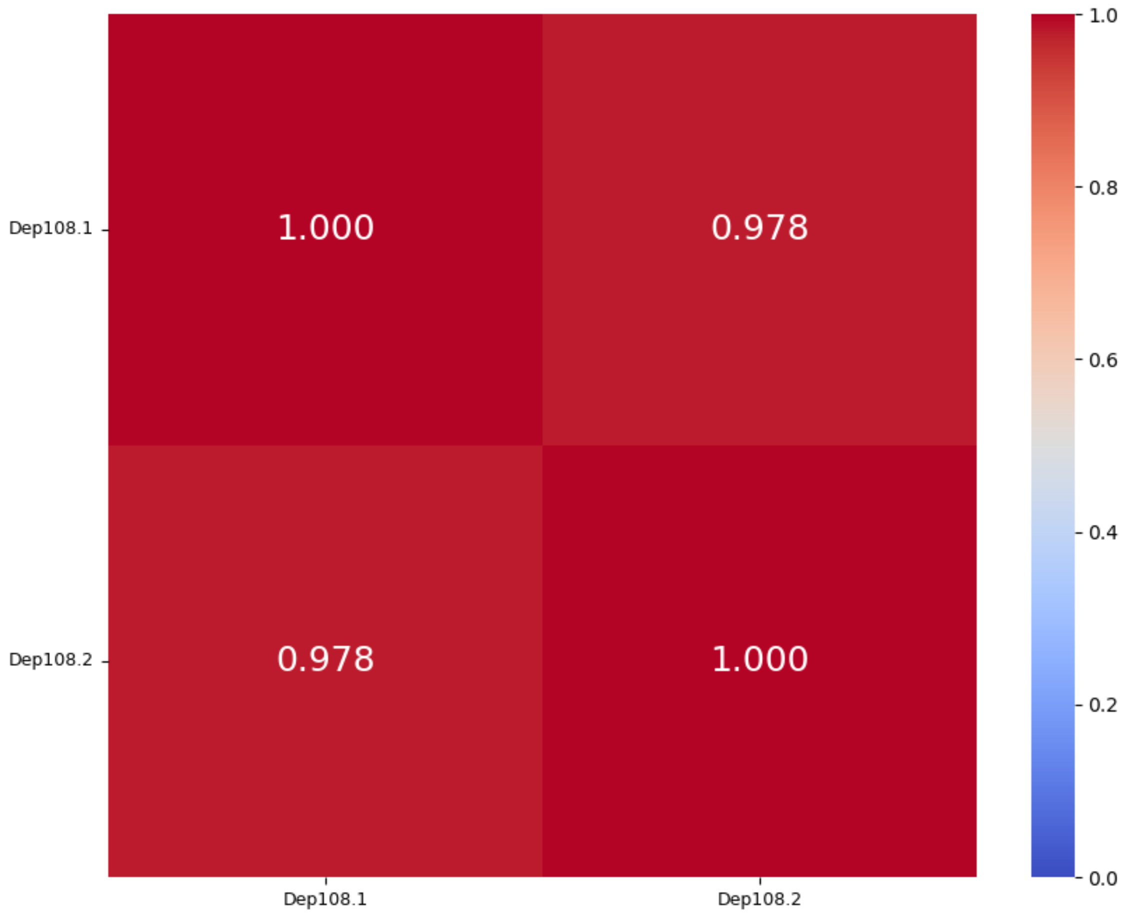


KL108 group 1
%identity; cover ≥ 99%
Central + C-term

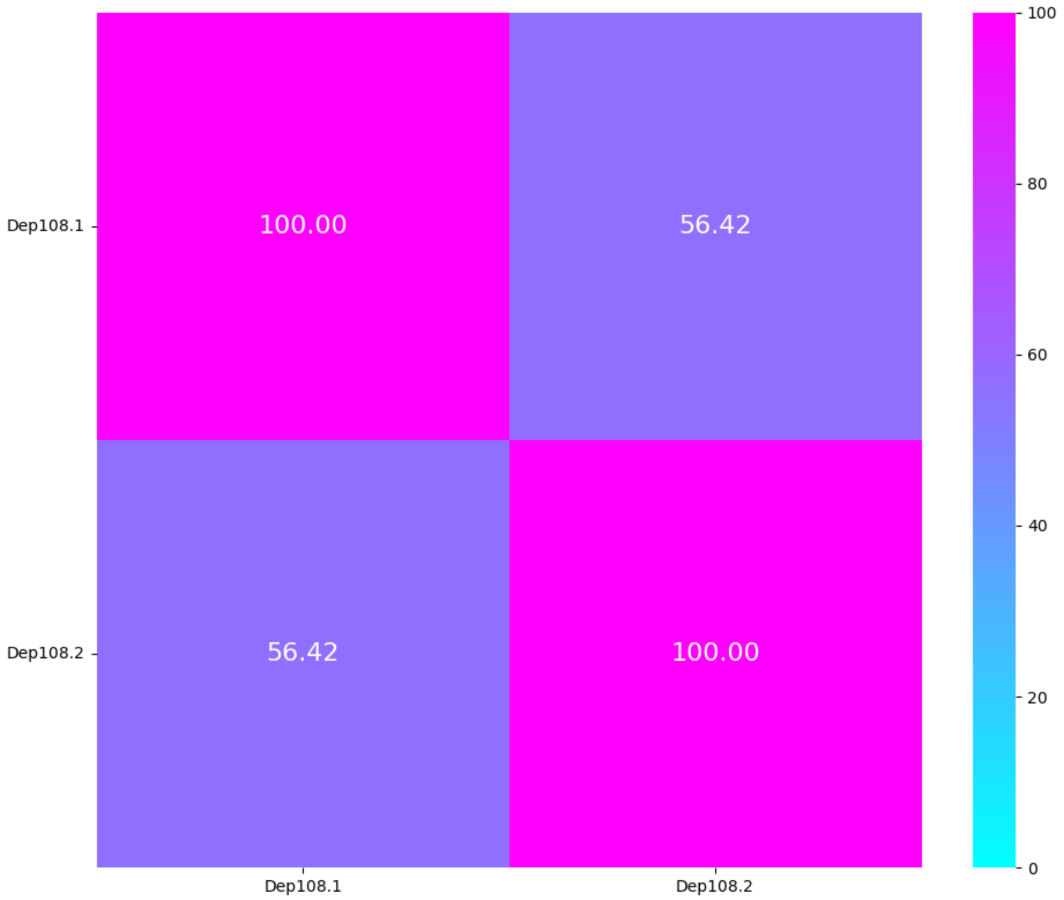

# KL111, KL114, KL116, KL122

| K-type target | K22/K37/KL111 | KL111 | | KL114 | KL116 | KL122 |
|---|---|---|---|---|---|---|
| Group | Group 1 | | | Group 1 | Group 1 | Group 1 |
| Protein name | KLEO13gp10 | 184_34 | KANgp232 | KP24gp307 | KANgp223 | 0496_72 |
| Lenght | 777 aa | 639 aa | 623 aa | 742 aa | 581 aa | 968 aa |
| Number of proteins in the group | 1 protein | 2 proteins | | 1 protein | 1 protein | 1 protein |
| Predicted protein structure | 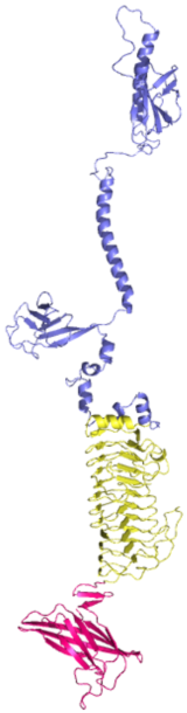 | 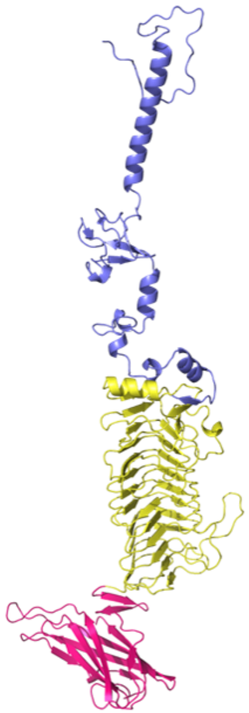 | 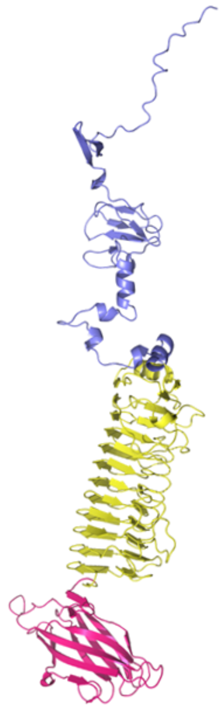 | 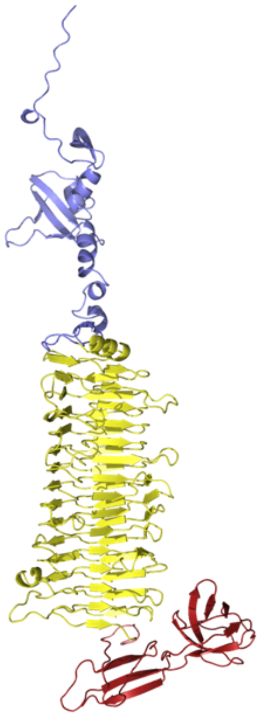 | 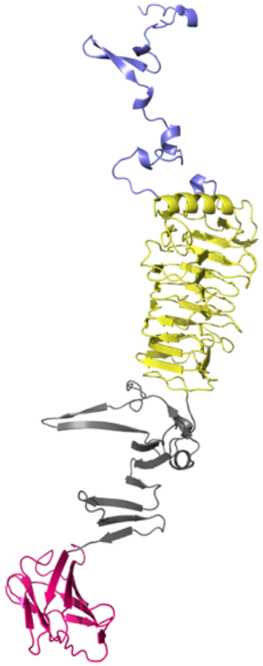 | 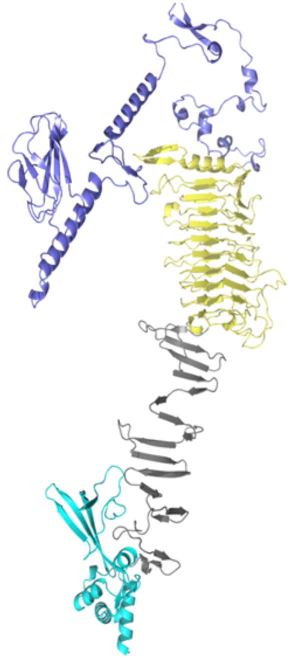 |

KL111 group 1
TM-score
Central + C-term

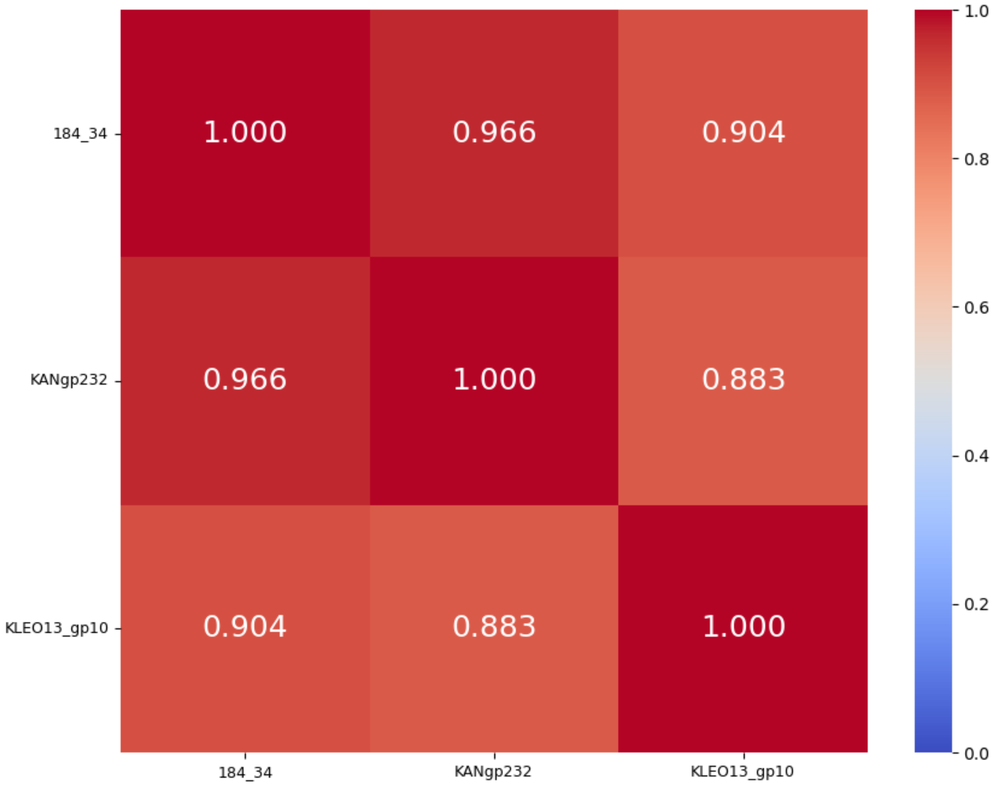


KL111 group 1
%identity
Central + C-term

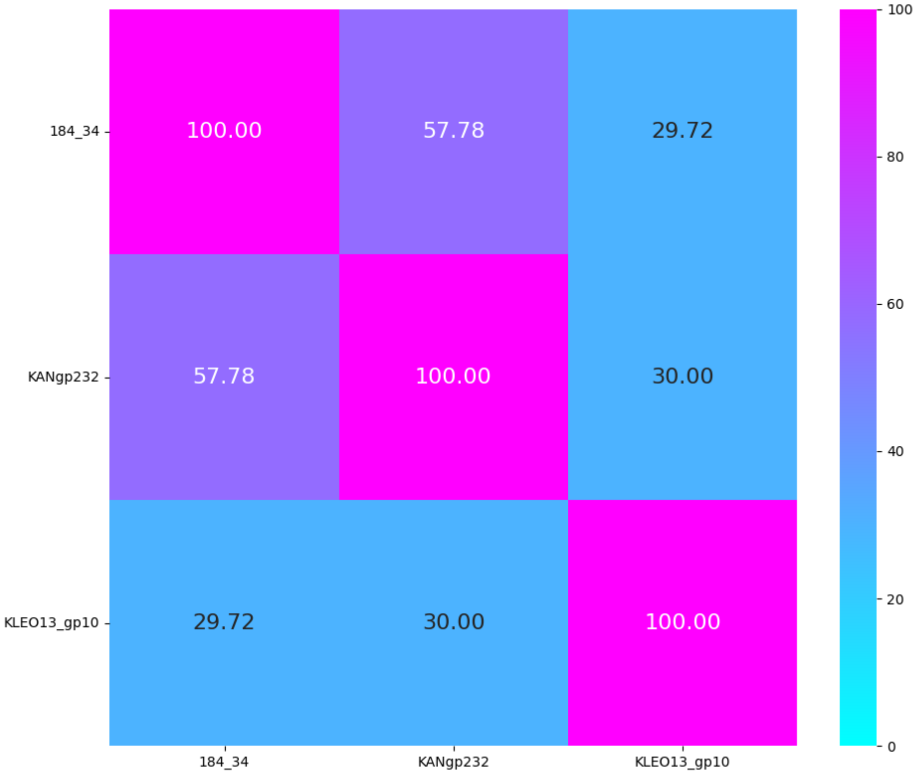

# KL127, KL134, KL137, KL143, KL153, KL158

| K-type target | KL127 | KL134 | KL137 | KL143 | K153 | KL158 |
|---|---|---|---|---|---|---|
| Group | Group 1 | Group 1 | Group 1 | Group 1 | Group 1 | Group 1 |
| Protein name | 319_37 | KP24gp168 | KP24gp306 | 738_68 | KANgp220 | KP24gp309 |
| Lenght | 794 aa | 881 aa | 883 aa | 742 aa | 644 aa | 751 aa |
| Number of proteins in the group | 1 protein | 1 protein | 1 protein | 1 protein | 1 protein | 1 protein |
| Predicted protein structure | 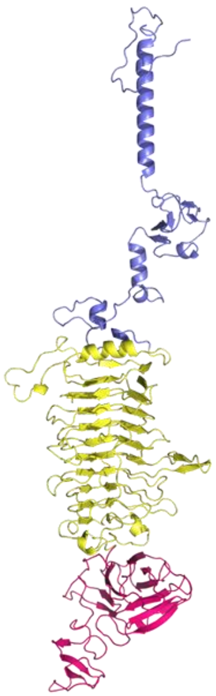 | 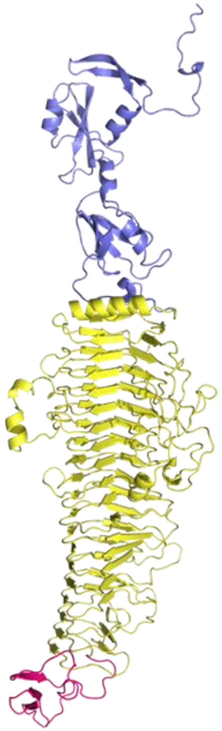 | 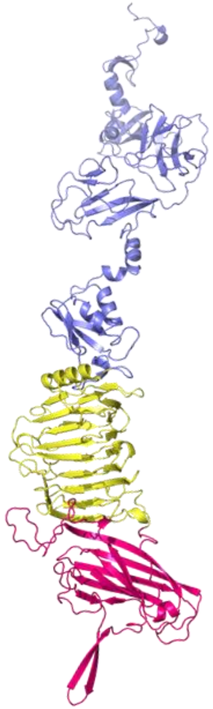 | 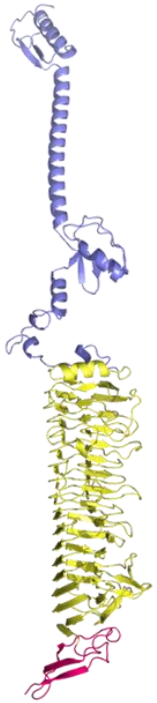 | 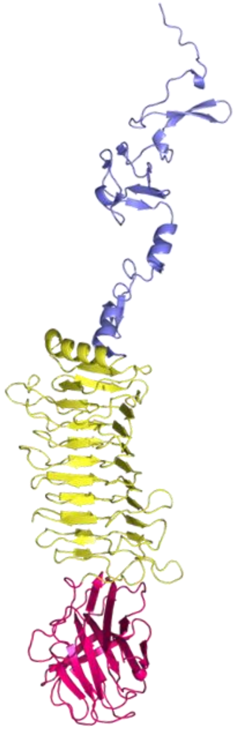 | 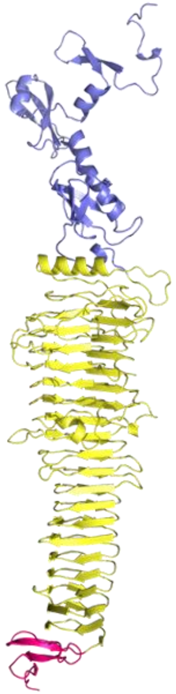 |

Only Central domain

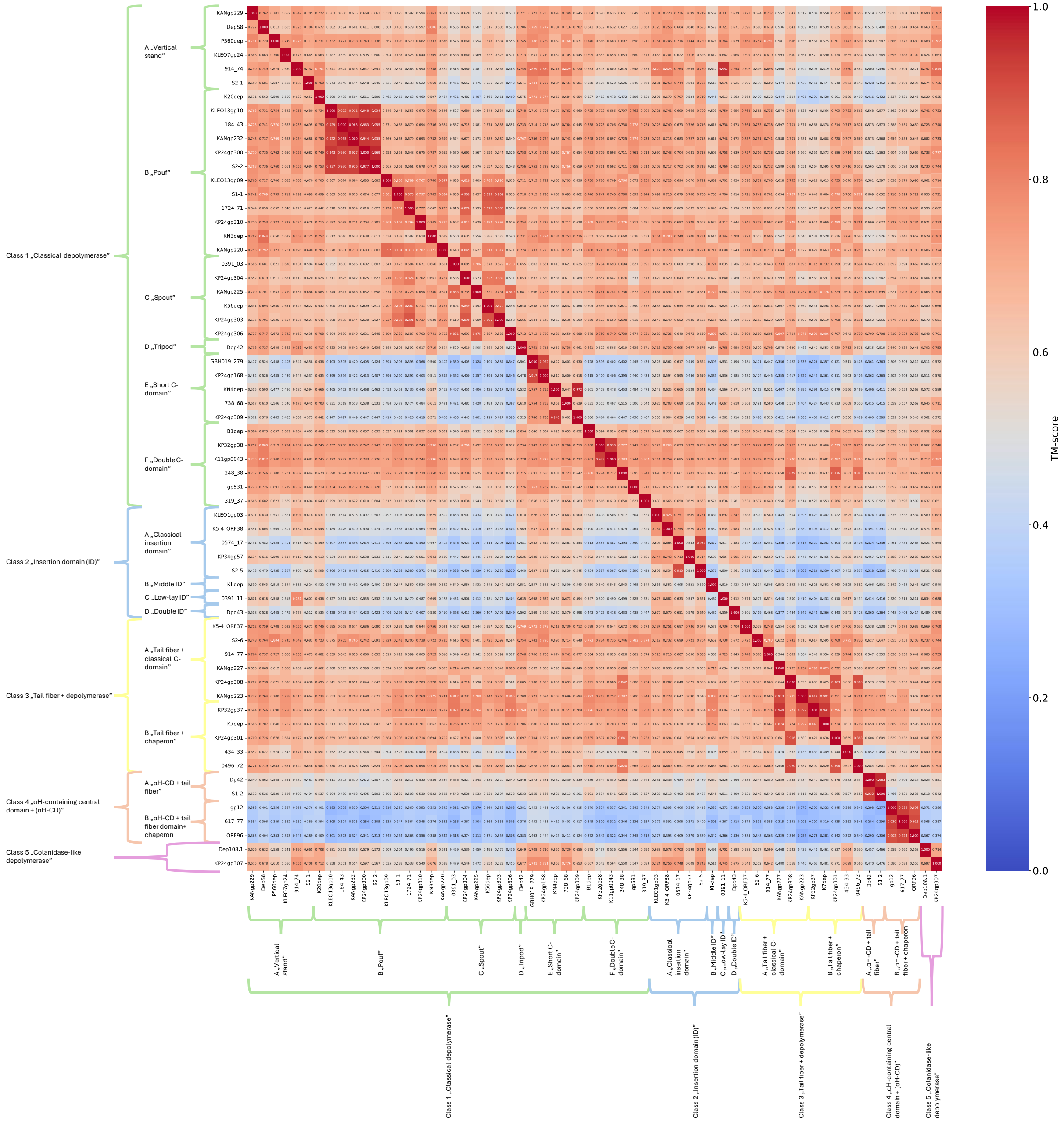

Only C-terminal domain

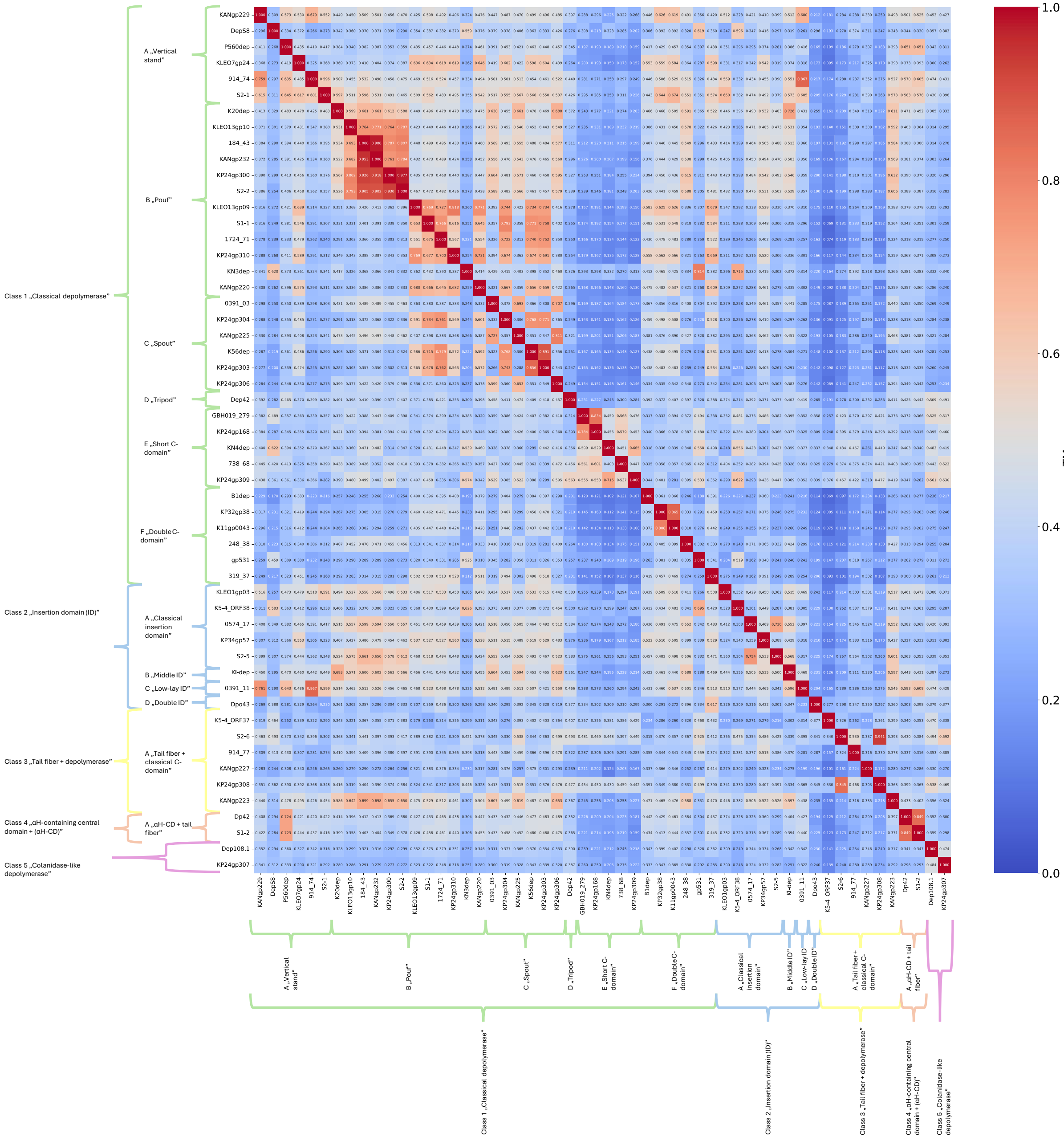

Only Chaperone

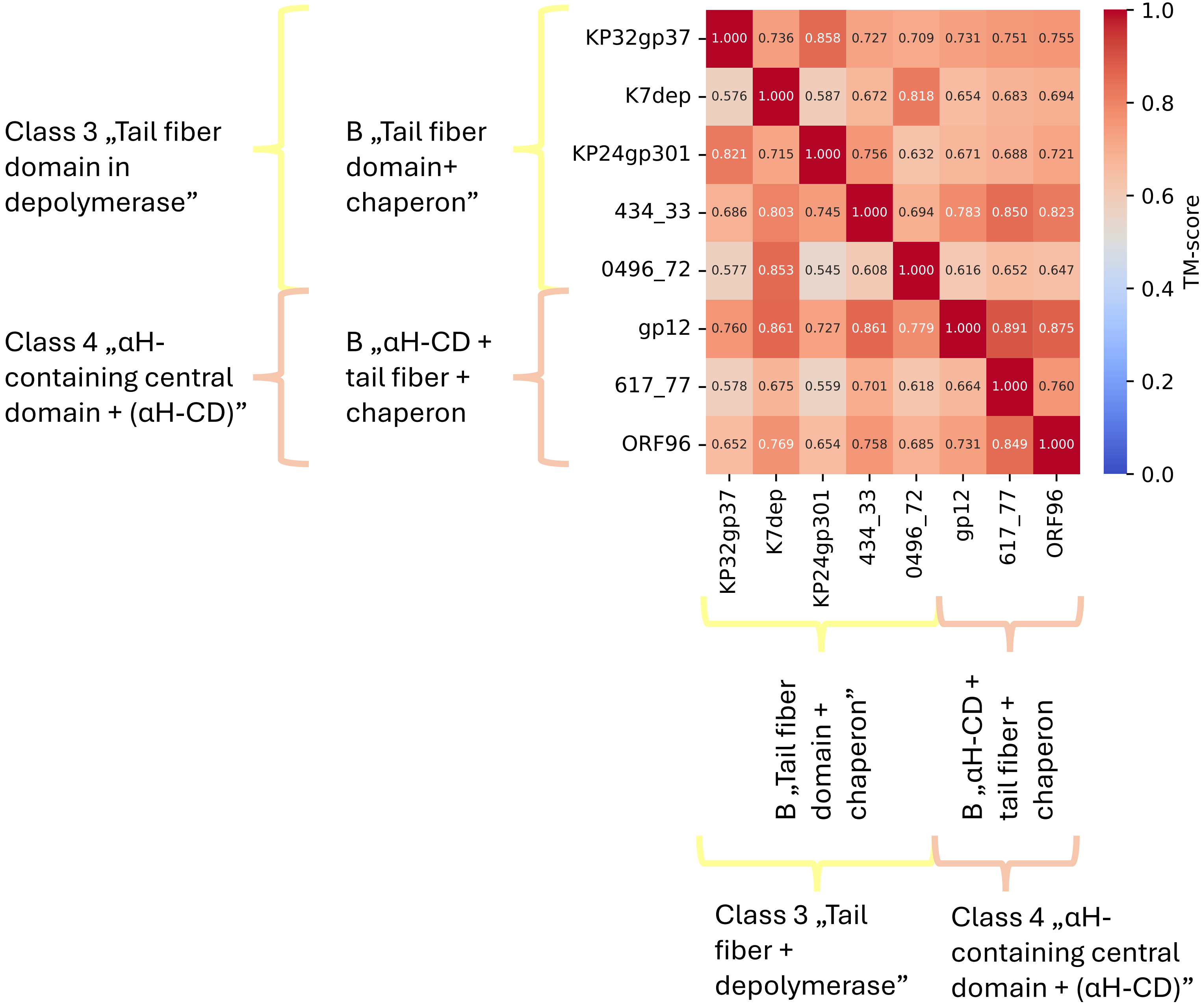
