## Supplementary material for "DepoCatalog: Mapping the Diversity of 105 Recombinant Klebsiella Phage Depolymerases Across Sequence, Structure, and Substrate Specificity": Suppl Figures S1-S5

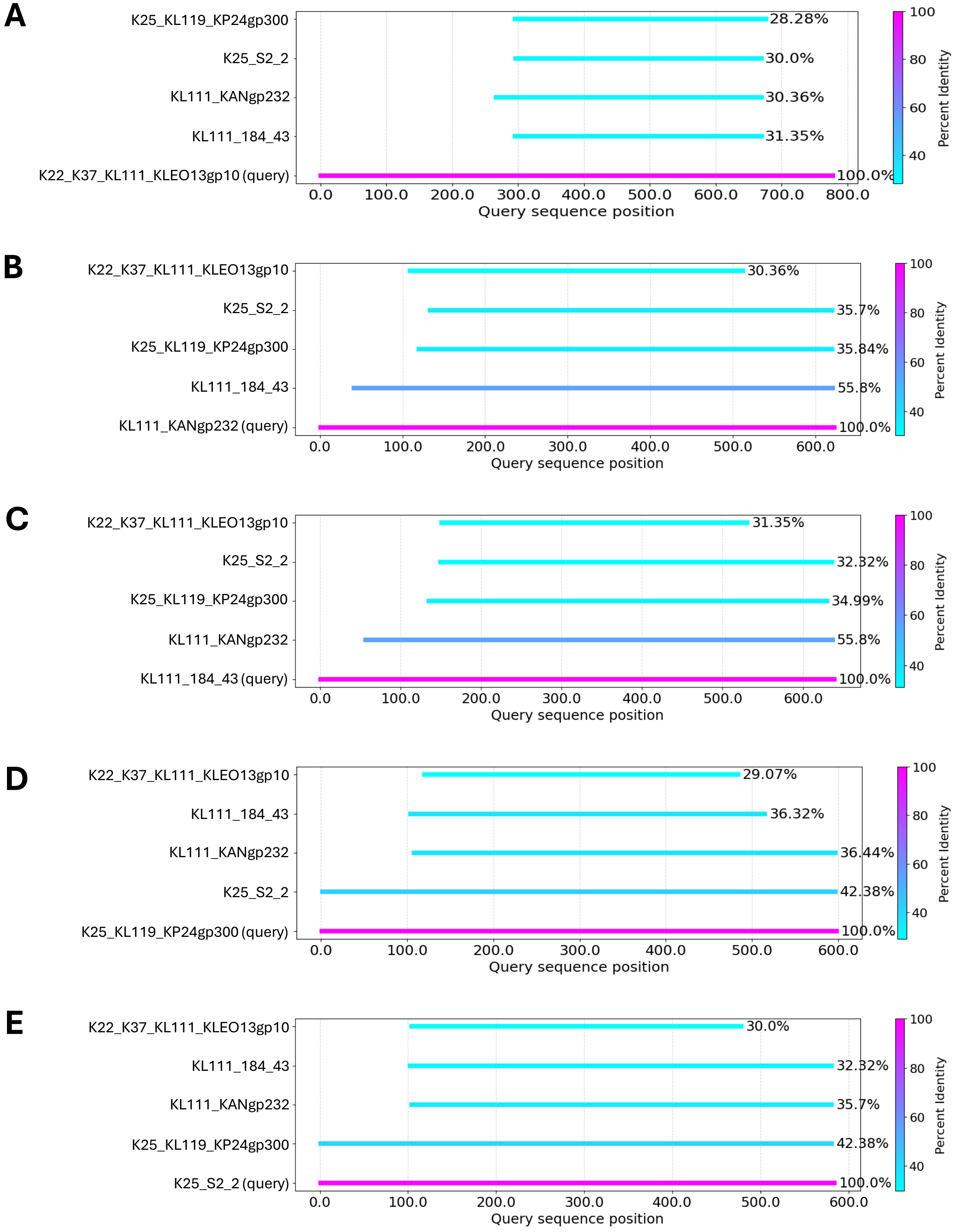


**Figure S1.** Pairwise amino acid sequence comparison of five *Klebsiella pneumoniae* CPS- degrading depolymerases using BLASTP. Each panel (A-E) displays an alignment of one query protein against remaining four.

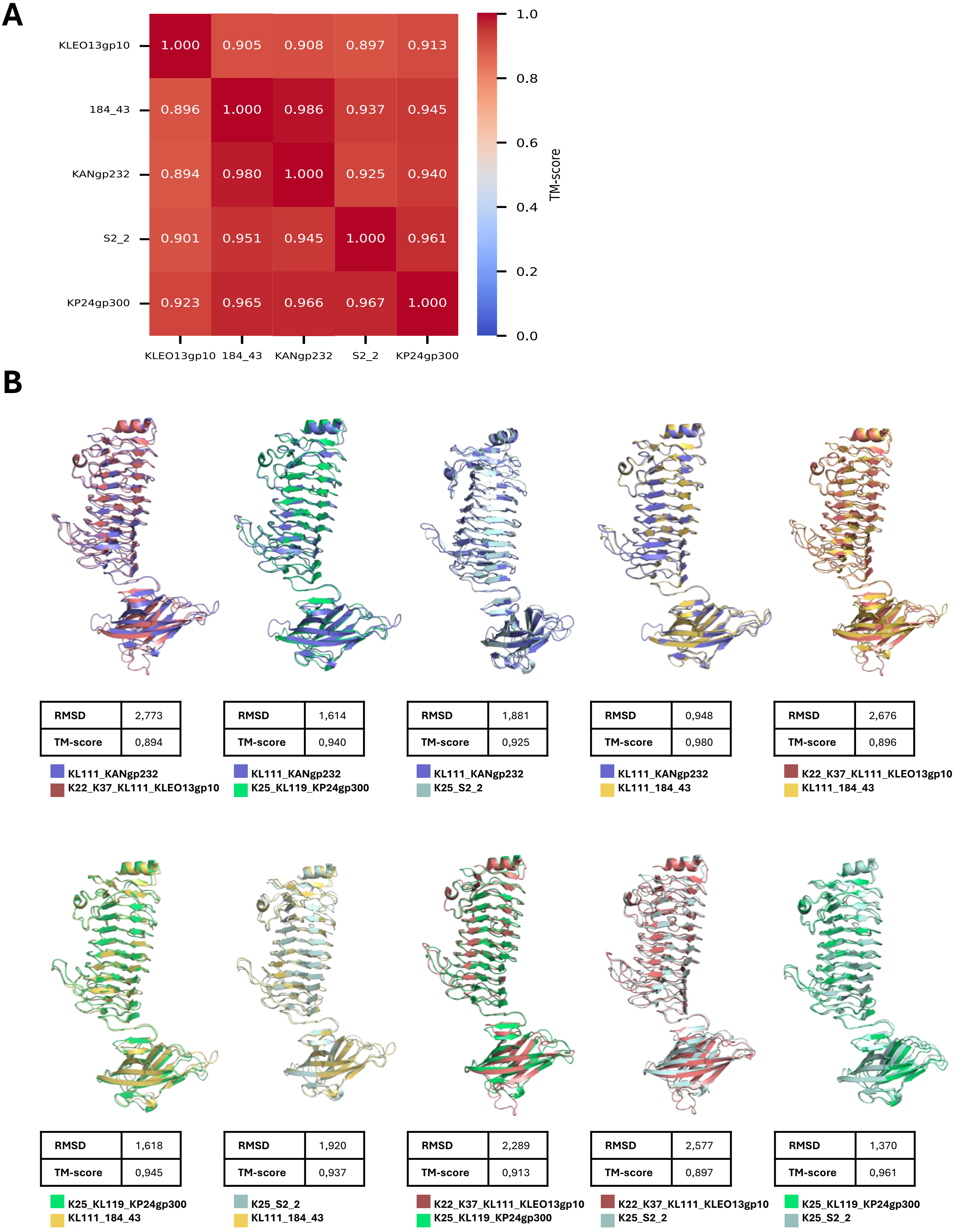


**Figure S2.** Structural comparison of five *K. pneumoniae* CPS- degrading depolymerases (excluding N-terminal domains). (A) Heatmap of pairwise structural similarity between depolymerases, calculated using USalign. Color intensity corresponds to TM- scores, with higher values indicating greater structural similarity. (B) Structural alignments of protein pairs visualised in PyMOL. Each alignment includes root-mean-square-deviation (RMSD) and TM- score values to quatify structural congruence.

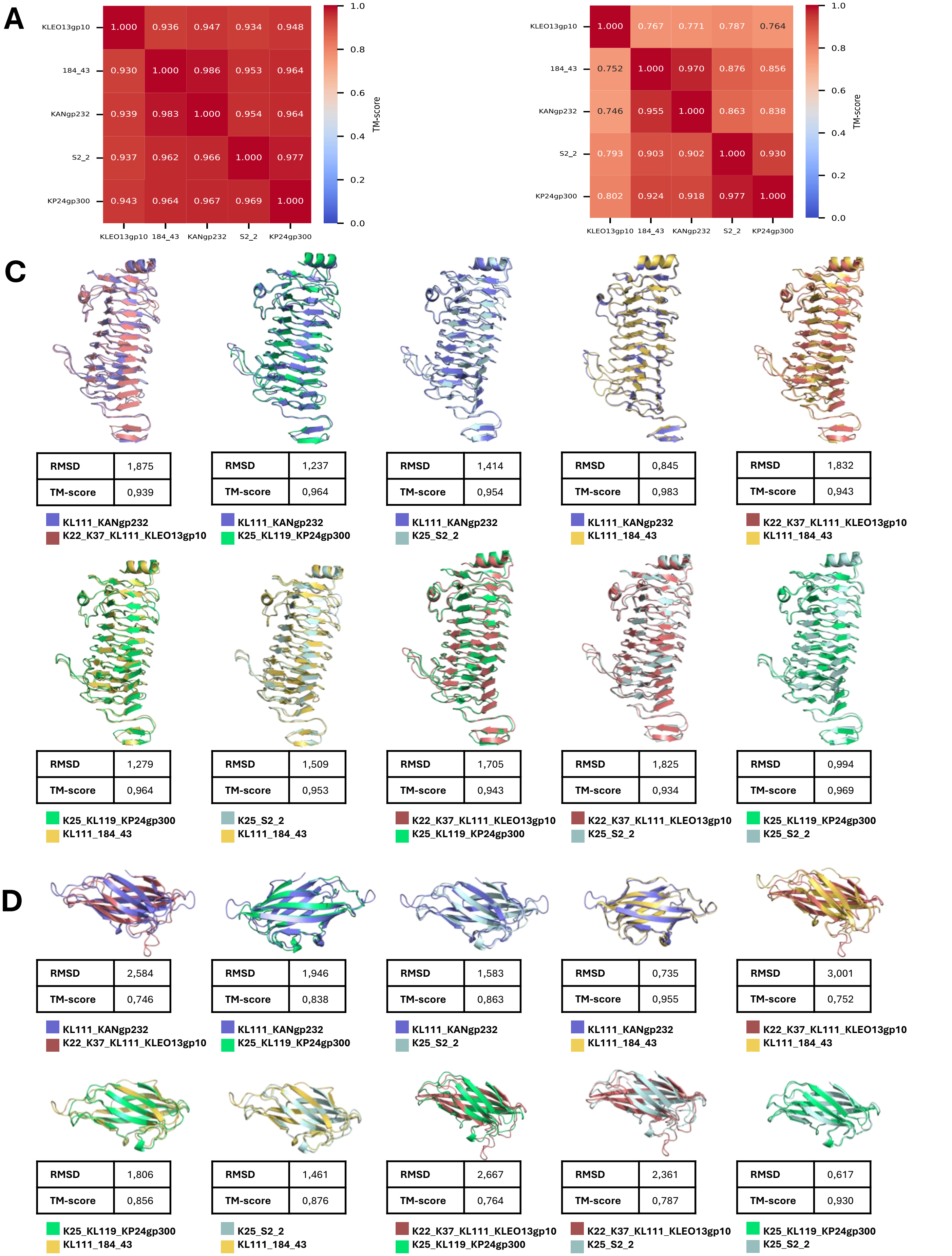


**Figure S3**. Domain-specific structural comparison of *K. pneumoniae* CPS- degrading depolymerases. (A) Heatmap of pairwise structural similarity between the central domains of depolymerases, computed using USalign. TM-scores are color-coded, with higher values indicating greater structural similarity. (B) Heatmap of C-terminal domains comparisons, also based on TM-scores calculated using USalign. (C) Structural alignments of the central domains visualized in PyMOL, with associated root-mean-square deviation (RMSD) and TM-score values provided for each pairwise alignment. (D) Structural alignments of the C-terminal domains, also shown with root-mean-square deviations (RMSD) and TM-scores.

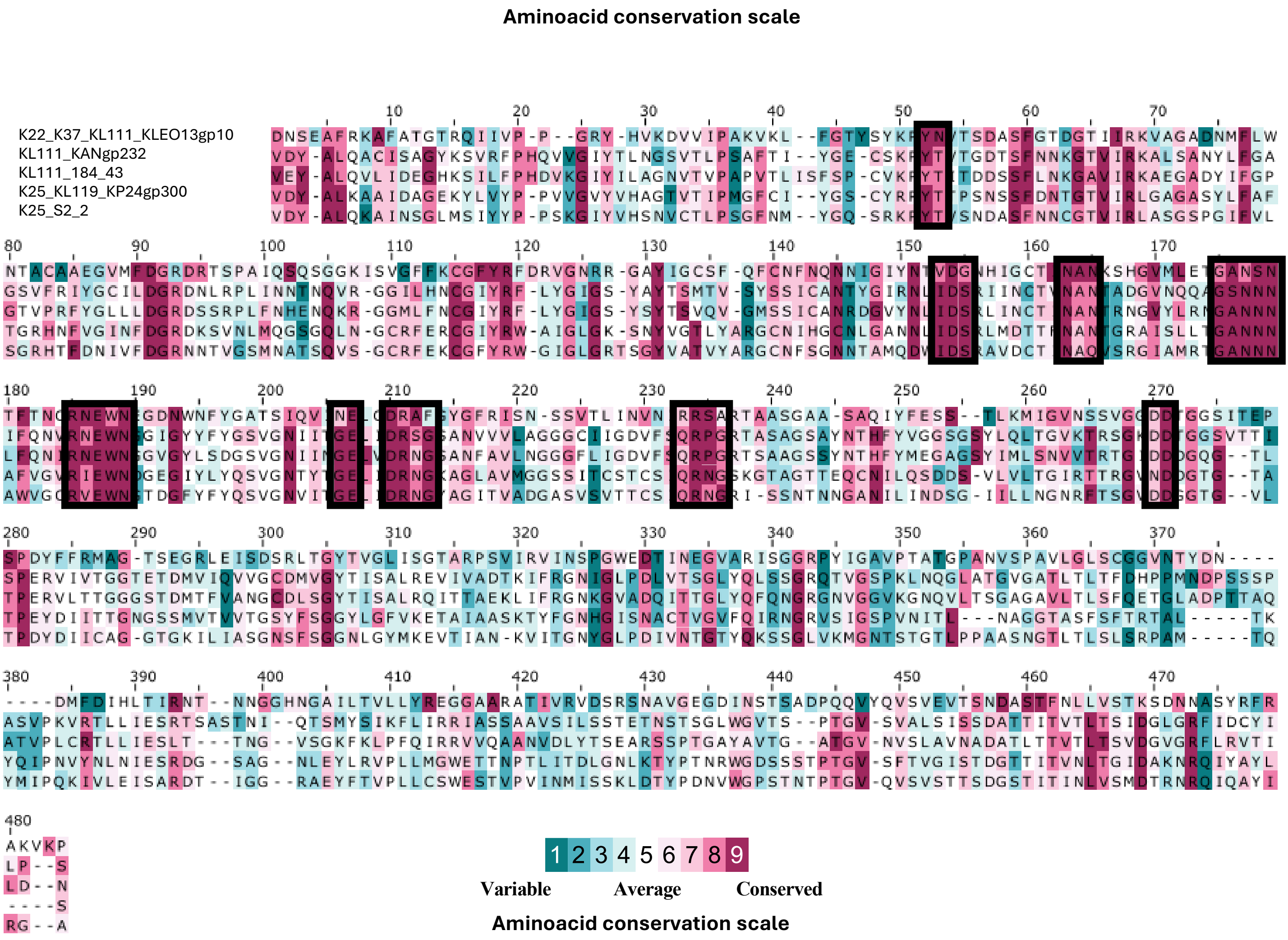


**Figure S4.** Conservation-based analysis of *K. pneumoniae* CPS- degrading depolymerases. Multiple sequence alignments (MSA) of five depolymerases, with amino acid conservation colouring according to ConSurf scores. Conservation level range from variable (low scores) to highly conserved (high scores), visually emphasizing conserved motifs. Residues corresponding to the predicted active site regions marked with black rectangles, highlighting their conservation across sequences and spatial co- localization in the structures. This representation provides a sequence-level view of conserved, functionally relevant regions across the protein set.

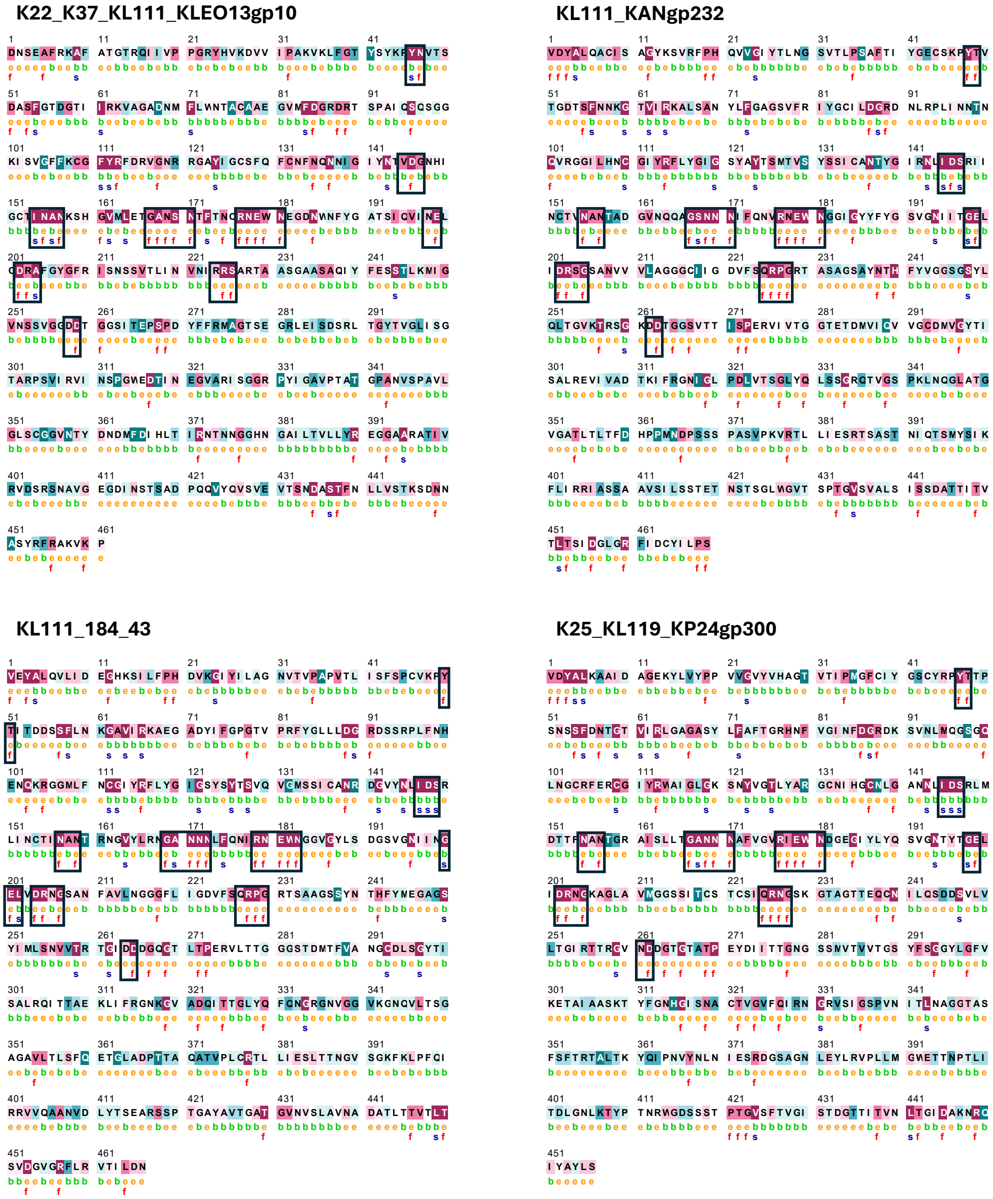


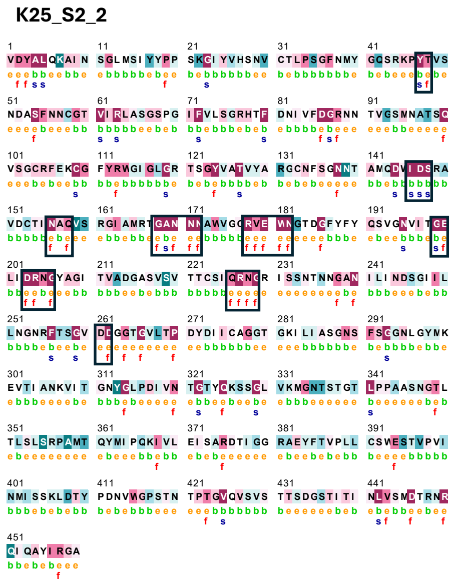


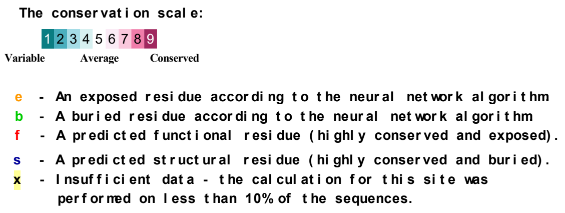


**Figure S5.** Conservation-based analysis of *K. pneumoniae* CPS- degrading depolymerases. Each panel displays amino acid sequence of depolymerase, colored according to residue conservation scores from ConSurf analysis. Below each amino acid, additional annotations indicate whether the residue is exposed or buried, or is classified as functionally, or structurally important. Residues corresponding to regions predicted as possible active site.
